# Massively parallel quantification and prediction of monomeric protein yield in *Escherichia coli*

**DOI:** 10.64898/2026.11.02.763507

**Authors:** Selena Setsu, Kaori Kiyokawa, Andy Y. W. Lam, Emi Mishiro-Sato, Yukihide Tomari, Kotaro Tsuboyama

## Abstract

Obtaining sufficient quantities of purified proteins in their intended molecular states is essential for biological research and protein engineering^1–4^. Advances in computational protein design have expanded the functional capabilities of de novo proteins and accelerated candidate generation^5–10^. However, reliable methods for ranking expected monomeric yield of candidate de novo designs, including molecular binders^11–15^, fluorescent proteins^16–19^ and enzymes^20–22^, have yet to be established^23^. Standardized measurements of monomeric yield across diverse sequences remain scarce, limiting our understanding of its determinants and the development of accurate predictors for candidate selection. Here we developed a scalable pooled assay combining peptide barcodes^24,25^, size-exclusion chromatography and mass spectrometry to quantify monomeric yield for over 39,000 natural protein domain variants, de novo proteins and nanobodies under uniform and standardized Escherichia coli production conditions. We used this dataset to (1) quantify the relationship between folding stability and monomeric yield, (2) benchmark inverse-folding methods^5,7,9,26–30^, (3) evaluate the utility of structure-prediction confidence metrics^31–33^ in selecting high-yield design candidates, and (4) identify sequence and structural patterns in nanobodies associated with monomeric yield and monomer-to-dimer yield ratios. We further developed ESMC-MonoYield, a protein language model^33^ fine-tuned to predict monomeric yield from amino acid sequence alone, and prospectively demonstrated enrichment of high-yield candidates among BindCraft-designed binders^8^. Our approach enables standardized, large-scale measurement of monomeric yield across diverse proteins, and establishes monomeric yield as a sequence-encoded property that can be learned, predicted and ultimately designed.

## Introduction

Producing proteins in sufficient quantities in their intended molecular states is essential for biochemical research, therapeutic development and industrial biocatalysis^2,34–36^. For natural proteins whose amino acid sequences must be preserved, improving yield largely involves optimizing expression constructs, hosts and production conditions^4,37–39^. De novo protein design, by contrast, allows many alternative structures and sequences to be generated for a common structural or functional objective^5,6,8–10,13,40–45^. Yield in the intended molecular state is therefore a critical criterion for ranking candidate sequences^46,47^. For proteins expected to be monomeric, the relevant endpoint is the yield of purified monomeric protein. This increases the value of predicting this yield under a uniform, low-cost production protocol^1,48^.

Predicting monomeric yield is challenging because it integrates several coupled factors and processes, and success at one stage does not guarantee success at the next. *Escherichia coli* (*E. coli*) is widely used because of its rapid growth, low cultivation cost and established expression tools^4,49^. Protein yield depends on host genotype^50–52^, plasmid architecture and promoter activity^3,53^, induction conditions and temperature^3,54^, and the cellular proteostasis machinery, including molecular chaperones^55–58^. Protein synthesis, solubility and monomeric recovery also depend on protein properties such as amino acid sequence, folding stability, hydrophobicity and aggregation propensity^46,59–74^, as well as on messenger RNA sequence and secondary structure^75–79^. Importantly, these aspects of protein production do not necessarily improve together: stabilizing mutations can increase total expression while reducing solubility^80,81^. Similarly, higher soluble yield does not necessarily translate into higher monomeric yield because soluble proteins can also form unintended oligomers^13^. Thus, measurements of protein abundance in total lysates or soluble fractions alone are insufficient to understand or predict monomeric recovery.

A range of experimental and computational approaches has been developed to assess protein production and related properties. Reporter-based assays enable high-throughput assessment of soluble expression^82–86^, but do not directly quantify purified monomeric yield. Systematic measurements across large panels of proteins provide expression and solubility data^64,87–89^, but differences in experimental conditions and readouts complicate data integration across studies. This variability has motivated calls for standardized production and characterization protocols^1^ and recent efforts to establish such workflows^47^. Even with standardized and efficient protocols, throughput remains constrained by DNA synthesis costs and the need to produce and analyse each protein in a separate well. Despite the limited availability of large-scale experimental data, several computational models have been developed to predict protein expression levels^90^, solubility^91–102^, aggregation propensity^103^, absolute protein folding stability^104^ and the functional effects of mutations^105,106^. Nevertheless, these predictions concern expression, solubility, aggregation or folding stability rather than purified monomeric yield. Structure-prediction confidence metrics, such as the predicted local distance difference test (pLDDT) score from AlphaFold models^31,32^, are also widely used to evaluate designed proteins. However, a benchmark of de novo designed proteins found that pLDDT provided only moderate discrimination between experimentally successful and unsuccessful designs, with performance varying across topologies^23^. These limitations underscore the need for large-scale measurements and predictive models of purified monomeric yield using a broadly applicable, uniform, and low-cost expression and purification protocol.

Here, we established a workflow based on peptide-barcode mass spectrometry (MS)^13,24,25,107–109^ to quantify purified monomeric yield of proteins. The workflow enables pooled analysis of up to approximately 2,600 proteins per flask following cytosolic expression in *E. coli*. Under uniform expression, induction and purification conditions, we obtained monomeric yield estimates, reported in mg/L, equivalent to individual expression of each protein, for 34,705 proteins spanning natural protein domains, their variants and de novo proteins. Using these data, we compared sequence-design methods and analyzed how folding stability and sequence variation relate to monomeric recovery. We then fine-tuned protein language models (pLMs)^33,110^ to predict monomeric yield and prospectively evaluated model-guided selection by individually producing newly designed binders. We also obtained monomeric yield estimates for an additional dataset of 2,590 nanobody sequences. Within this dataset, sequences satisfying rules previously proposed for yeast-display libraries^111^ showed higher monomeric recovery and monomer-to-dimer ratios than sequences that did not. We further used these measurements to develop nanobody-specific pLM predictors for both outcomes. Together, this workflow, the resulting datasets and the predictive models provide a basis for selecting sequences with improved monomeric recovery under general and uniform production conditions from the *E. coli* expression system.

## Results

### A calibrated pooled assay for monomeric protein yield

To quantify monomeric protein yield at scale, we adapted a peptide barcode-based mass spectrometry approach (Fig. 1a). Peptide barcoding has previously enabled pooled characterization and selection of target proteins, including binders and nanobodies^24,25,108,109^. Building on these applications, we sought to calibrate and validate barcode signals for absolute quantification of soluble monomeric protein yield. In this workflow, proteins carrying pre-assigned peptide barcodes were expressed in pooled *E. coli* cultures, purified by His-tag affinity beads, and separated by size-exclusion chromatography (SEC). The monomeric fraction was collected, and peptide barcodes were isolated by sequential proteolysis and affinity purification before quantification by liquid chromatography–tandem mass spectrometry (LC–MS/MS).

**Figure 1.**
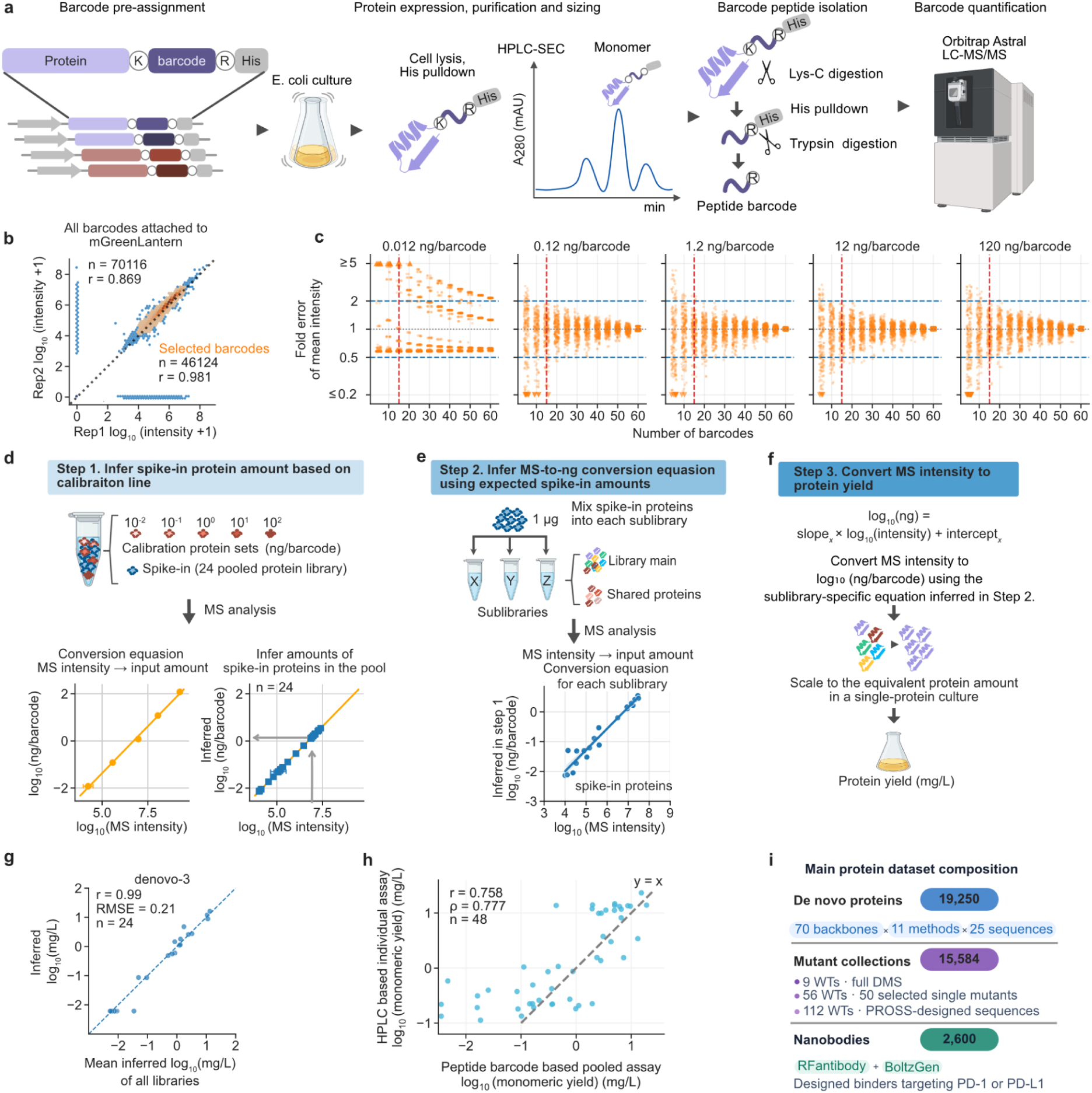
Development and validation of a calibrated peptide-barcode assay for monomeric protein yield. **a,** Overview of the pooled assay. Protein sequences were assigned unique peptide barcodes and expressed in *E. coli*. Following cell lysis and His-tag affinity purification, monomeric proteins were isolated by high-performance liquid chromatography–size-exclusion chromatography (HPLC–SEC). Barcode peptides were released and isolated by sequential Lys-C digestion, His-tag pulldown and trypsin digestion, and quantified by Orbitrap Astral LC–MS/MS. **b,** Selection of peptide barcodes with intermediate MS signal intensities for robust quantification. A total of 70,116 candidate barcodes from Feldman et al. (2025)^25^ were attached to mGreenLantern and compared between replicate measurements. Orange points indicate the 46,124 selected barcodes. Pearson correlations were *r* = 0.869 for all candidate barcodes and *r* = 0.981 for the selected subset. **c,** Effect of barcode number on quantification accuracy. Five GFP-nanobody calibration sets, each containing 60 barcodes, were pooled at input levels of 0.012, 0.12, 1.2, 12 and 120 ng per barcode, respectively. The pool was measured four times. For each measurement and input level, barcode subsets were computationally sampled 50 times at each barcode number. Points show the fold error of the mean MS intensity for each subset, using the mean across the four replicates at the corresponding input level as the reference. At the lowest input level (0.012 ng per barcode), fold errors remained large even when all 60 barcodes were used. Blue dashed lines mark 0.5-fold and twofold error; the red dashed line marks 15 barcodes. **d–f,** Conversion of MS intensities to absolute monomeric yields. **d,** Step 1: The intensity-to-amount relationship established using GFP-nanobody sets at known input amounts was used to infer the amounts of 24 spike-in proteins. **e,** Step 2: A 1-µg aliquot of the spike-in pool was added to each sample after SEC. Measured spike-in intensities and the amounts inferred in Step 1 were used to establish a sublibrary-specific intensity-to-amount relationship. **f,** Step 3: Barcode intensities were converted to protein amounts and scaled by sampling fraction, culture volume and pooled library size to obtain monomeric yields expressed as individual-culture equivalents (mg/L). **g,** Agreement of yield estimates for shared control proteins. Estimates for 24 proteins in the representative denovo-3 sublibrary are plotted against their mean estimates across all sublibraries (Pearson’s r = 0.99; RMSE = 0.21). Both axes show log₁₀ yield in mg/L. (see also Extended Fig. 2a) **h,** Comparison of MS-based pooled-assay estimates with monomeric yields measured by HPLC–SEC after individual expression and His-tag purification. (n = 48 proteins; Pearson’s r = 0.758; Spearman’s ρ = 0.777). Both axes show log₁₀ yield in mg/L. The dashed line denotes y = x. **i,** Composition of the 37,434 experimental entries: 19,250 inverse-folding designs, 15,584 mutant-collection entries and 2,600 nanobody designs. The mutant collection includes full scans of nine proteins, selected single mutants from 56 proteins and PROSS-designed variants from 112 proteins. Sublibrary composition is provided in Extended Fig. 1b.

We first selected barcode sequences with reliable detection. We attached 70,116 candidate barcodes from Feldman et al. (2025)^25^ to mGreenLantern^112^ and compared MS intensities of barcodes between two independently prepared samples. Filtering for detection, signal intensity and agreement between preparations retained 46,124 barcodes, with signals approximately between 10⁴ and 10⁷. Pearson’s r increased from 0.869 for all candidates to 0.981 for the selected set (Fig. 1b).

To increase the number of proteins that could be analyzed in a single experiment, we next determined how few barcodes were required per protein while maintaining accuracy. We prepared calibration nanobody sets at five known input levels spanning 0.012–120 ng per barcode, with each level represented by a distinct set of 60 peptide barcodes from the preselected peptide pool. We computationally subsampled these barcode sets and evaluated the fold error of the mean MS intensity as a function of barcode number (Fig. 1c). Because at the lowest input level (0.012 ng per barcode), fold errors remained large even when all 60 barcodes were used, we excluded this level when determining the required number of barcodes. We found that 15 barcodes per protein were sufficient to achieve quantification within twofold error for more than 90% of randomly sampled barcode sets at each of the remaining input levels (0.12, 1.2, 12, and 120 ng per barcode). Unless otherwise noted, we therefore assigned 15 peptide barcodes to each protein.

As an independent assessment of reproducibility across barcode assignments, we constructed two pilot libraries containing the same 1,038 proteins, each represented by 40 barcodes, but with different barcode assignments between libraries. Mean barcode intensities were highly correlated between the two libraries (Pearson’s r = 0.959; Extended Fig. 1a), consistent with previous work demonstrating that combining signals from multiple peptide barcodes enables quantitative estimation of protein abundance^24^. Computational subsampling further showed that agreement remained high with 15 barcodes per protein, with a representative comparison yielding Pearson’s r = 0.928 (Extended Fig. 1a). These results further supported the use of 15 barcodes per protein for reliable protein-level quantification.

To extend the assay to larger protein collections, we established a three-step calibration pipeline that converts peptide-barcode intensities into absolute protein amounts, enabling measurements from multiple sublibraries to be combined on a common quantitative scale (Fig. 1d–f). In Step 1, the calibration nanobody sets (also used in Fig. 1c) were mixed at known amounts with a pooled spike-in standard comprising 24 proteins, each assigned 15 barcodes. Calibration input and MS intensity were strongly related on a log–log scale (R² = 0.994; Fig. 1d). The fitted relationship was used to infer the amount of each protein in the spike-in standard (Fig. 1d). For subsequent steps, the quantified spike-in pool provided a common reference for absolute quantification across sublibraries. Using this pool simplified sample preparation compared with the calibration nanobody sets, facilitating extension to larger numbers of sublibraries. In Step 2, each sublibrary was cloned into a pET-derived plasmid and expressed in an *E. coli* BL21(DE3)-derived host using an autoinduction medium at 37 °C overnight. After His-tag purification, HPLC-SEC separation, and collection of the monomeric fraction, 1 µg of the quantified spike-in pool was added to each sublibrary sample. Following barcode isolation and MS analysis, the predetermined amounts of the spike-in proteins were used to fit a sublibrary-specific relationship between barcode intensity and protein amount (Fig. 1e; Extended Fig. 1b and 1c). In Step 3, barcode intensities were converted into protein amounts using the sublibrary-specific calibration equation (derived from Step 2). The amounts were then scaled by the sampled fraction, culture volume and number of barcoded constructs to estimate monomeric yields expressed as individual-culture equivalents (mg/L) (Fig. 1f; Methods).

We assessed the resulting yield estimates in two ways. First, shared control proteins included in each sublibrary showed strong agreement across sublibraries, with Pearson (r) values ranging from 0.92 to 0.99 when the estimates from each sublibrary were compared with the corresponding cross-sublibrary means. (Fig. 1g; Extended Fig. 2a). Second, we compared pooled-assay estimates with monomeric yields measured by HPLC–SEC after individual expression and His-tag purification of 48 proteins (Extended Fig. 3). Pooled-assay estimates generally agreed with monomeric yields measured after individual expression and purification, with most data points lying near the identity line (y = x; Pearson’s r = 0.758; Spearman’s ρ = 0.777; Fig. 1h). However, agreement declined at pooled-assay yields below approximately 0.1 mg/L, consistent with the reduced quantification reliability observed at the lowest calibration input (0.012 ng per barcode; Fig. 1c). Based on this result, we set the lower limit of reliable quantification at −1.5 (approximately 0.032 mg/L) and clipped lower values to this threshold for further analysis and model construction.

In total the experimental protein collections comprised 37,434 entries: 19,250 de novo designs for inverse-folding benchmarking, 15,584 mutants and 2,600 designed nanobodies, as summarized in Fig. 1i. In addition to these collections, we included a library of 2,500 synonymous coding-sequence variants, comprising 50 variants for each of 50 proteins, to assess the effects of codon usage on monomeric yield (Extended Fig. 1b). Of the 622,710 peptide barcodes, 614,152 (98.6%) were detected by DNA sequencing, and 596,881 (95.8%) were retained after excluding entries with counts per million below 2. This enabled monomeric yield estimation for 39,728 proteins (protein collections+coding-sequence variants), representing 99.5% of all proteins in the library. These collections supported comparisons of design methods, analyses of stability and yield, and training of yield predictors.

### Codon usage does not override sequence dependent differences in monomeric yield

We examined whether synonymous coding-sequence variation could account for differences in monomeric yield among proteins. We generated 50 synonymous coding sequences for each of 50 proteins, spanning broad ranges of codon adaptation index (CAI) and tRNA adaptation index (tAI) values,^113,114^ and evaluated them under the same expression and culture conditions. Monomeric yields of 85.7% of synonymous variants were within threefold of the median for the corresponding protein (Extended Fig. 4a). For 96% of proteins, the geometric standard deviation of monomeric yield across synonymous variants was less than threefold (Extended Fig. 4b). By comparison, representative proteins showed yield differences spanning several orders of magnitude over overlapping CAI and tAI ranges (Extended Fig. 4c). These results indicate that synonymous sequence variation modulated yield of individual protein but did not override the broad differences among proteins under the conditions tested.

To minimize the synonymous variation effects, in the main libraries, coding sequences were designed using DNAWorks and optimized for *E. coli* codon usage. Among coding sequences for de novo designs and DMS wild-type controls, the central 95% ranges (2.5th–97.5th percentiles) were 0.628–0.959 for CAI and 0.221–0.348 for tAI, with medians of 0.835 and 0.298, respectively (Extended Fig. 4d).The main libraries therefore used codon-optimized sequences spanning a much narrower range of codon usage than the synonymous-variant library, allowing us to focus on associations between amino acid sequence and monomeric yield.

### Backbone dependence and benchmarking of inverse folding methods

De novo protein design typically involves generating a backbone structure (e.g. RFdiffusion^6^, BoltzGen^9^) and designing amino acid sequences that fold into that structure, a task known as inverse folding. A key question is how backbone choice and sequence-design method influence monomeric yield, and whether structure-prediction-based filters commonly used to select promising designs also enrich proteins with high monomeric yield. To address this question, we systematically compared inverse-folding methods on a common set of backbones and evaluated the benefits and limitations of structure-based filtering. We benchmarked 11 inverse-folding methods or parameter settings^5,7,9,26–30,115^ on 70 backbone structures, generating 25 sequences per backbone–method combination for a total of 19,250 designs (Fig. 2a). The backbone set contained 58 de novo-designed structures and 12 natural domains and covered all-α, α/β and all-β architectures (Extended Fig. 5a). Applying all methods to the same backbone set enabled comparisons of yield across backbones and among sequence-design methods within each backbone.

**Figure 2.**
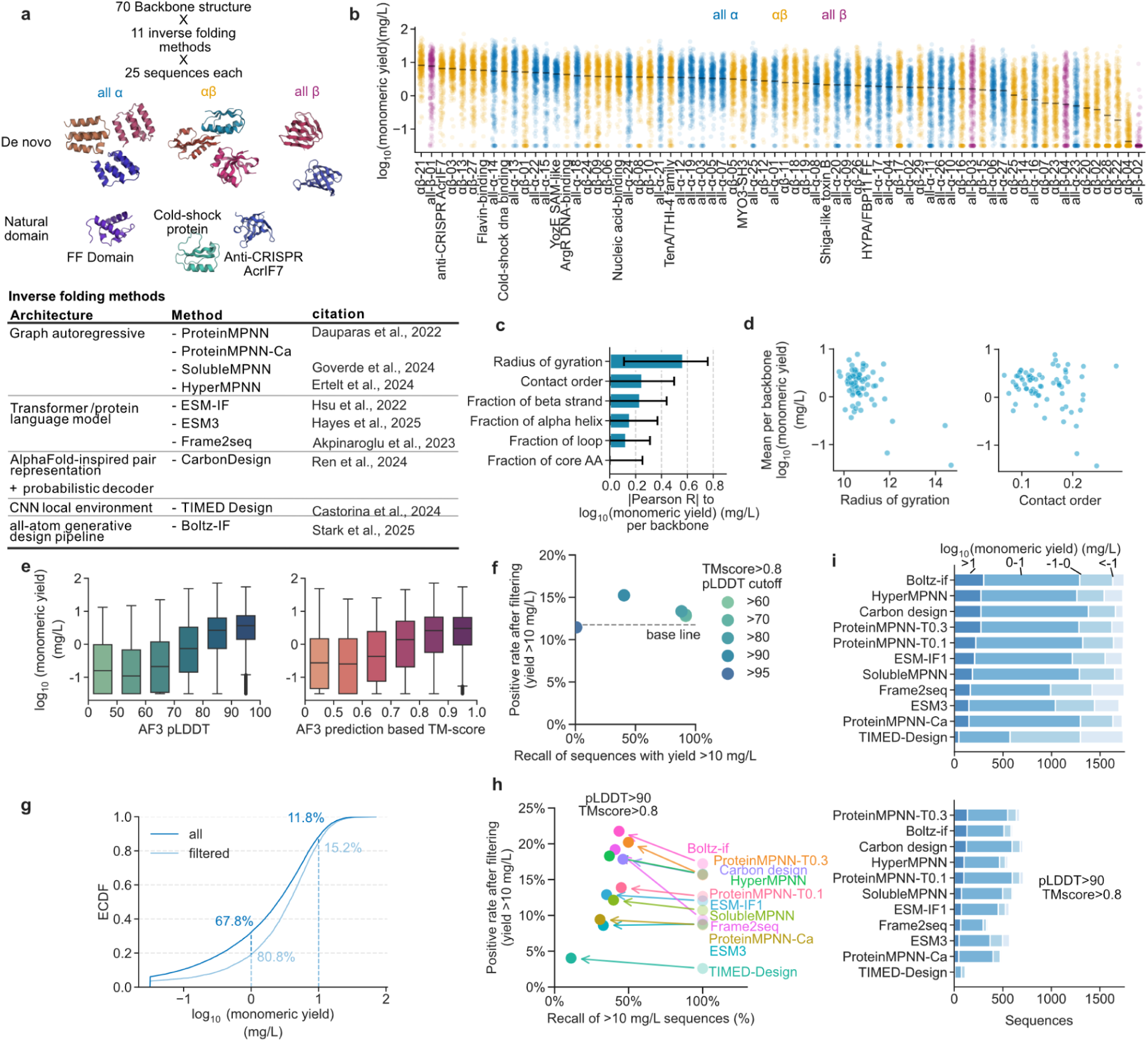
Large-scale benchmarking of inverse-folding methods and assessment of the benefits and limitations of AlphaFold-based filtering. **a,** Design of the inverse-folding benchmark. Eleven methods or parameter settings were applied to 70 protein backbones, with 25 sequences generated per backbone–method combination. Representative de novo and natural-domain backbones are shown, together with the model architectures and references for the evaluated inverse-folding methods. **b,** Monomeric yields of individual designed sequences grouped by backbone. Lines indicate the median yield for each backbone. Colours indicate backbone structural classes: all-α, α/β and all-β. **c,** Absolute Pearson correlations between backbone-level mean log₁₀ monomeric yield and structural descriptors. Bars show correlation magnitudes; error bars show 95% bootstrap confidence intervals (1,000 resamples). **d,** Relationships between backbone-level mean of log₁₀ monomeric yield and radius of gyration (left) or contact order (right). Each point represents one backbone. **e,** Distributions of monomeric yield grouped by AlphaFold3 (AF3) pLDDT (left) and TM-score between the AF3-predicted structure and the target backbone (right). **f,** Effect of structure-based filtering on the selection of high-yield proteins, defined as proteins with monomeric yield >10 mg/L. For a fixed TM-score cutoff of >0.8, different pLDDT cutoffs were evaluated. The positive rate is the fraction of retained sequences exceeding the yield threshold; recall is the fraction of all high-yield sequences retained after filtering. The dashed horizontal line indicates the unfiltered positive rate. **g,** Empirical cumulative distribution functions (ECDFs) of monomeric yield before and after filtering with AF3 pLDDT >90 and TM-score >0.8. **h,** Method-specific effects of the filter used in g. Arrows connect the unfiltered and filtered positive rates and recalls for each inverse-folding method. **i,** Numbers of sequences in the indicated monomeric-yield ranges for each inverse-folding method before (top) and after (bottom) filtering. Stacked bar segments correspond to log₁₀ yield ranges of >1, 0 to 1, −1 to 0 and <−1, with yield expressed in mg/L.

We first examined backbone-dependent variation in monomeric yield. Yield varied substantially among backbones, and several backbones produced low-yield sequences across methods (Fig. 2b). Secondary-structure composition alone did not account for this variation. We therefore compared backbone-level mean monomeric yield with backbone-dependent but sequence-independent features. Among such features, radius of gyration showed the largest absolute correlation among these descriptors (Fig. 2c,d; Extended Fig. 5b). Its negative association with yield indicated that more compact backbones tended to support higher recovery in this set.

We next examined whether structure-prediction confidence or agreement with the design backbone could enrich high-yield sequences. For both AlphaFold2 and AlphaFold3, the highest pLDDT and TM-score bins generally contained designs with higher median yields (Fig. 2e; Extended Fig. 5c,d). Filtering by TM-score alone produced only modest shifts in the yield distribution, whereas stringent pLDDT filtering produced a clearer shift (Extended Fig. 5e). Filtering with AlphaFold3 pLDDT >90 and TM-score >0.8 increased the fraction of high-yield designs (>10 mg/L) from approximately 12% to 15%, while retaining approximately 40% of all high-yield designs; raising the pLDDT threshold to >95 provided no further enrichment (Fig. 2f). AlphaFold2-based filtering showed less enrichment across the tested thresholds (Extended Fig. 5f). However, the AlphaFold3 filter increased the fraction of designs yielding >1 mg/L from 68% to 81% (Fig. 2g), indicating that it more effectively excluded low-yield designs than enriched high-yield designs. Thus, stringent structural-confidence filtering modestly increased the fraction of high-yield designs while excluding many experimentally useful sequences, consistent with the limited predictive power of confidence metrics for experimental design success reported previously^23^.

Using AlphaFold3 pLDDT >90 and TM-score >0.8, we compared inverse-folding methods before and after filtering (Fig. 2g–i). Boltz-IF, ProteinMPNN (Temperature = 0.3), CarbonDesign and HyperMPNN were among the higher-yielding methods before filtering. Filtering largely preserved this grouping, but improved the positive rate of Frame2seq from approximately 10% to approximately 20%. For the other methods, the filter preferentially removed low-yield sequences, yet also reduced the number of designs by ∼60%. Filtering also reduced backbone coverage: ProteinMPNN at T = 0.1 or 0.3 and CarbonDesign only approximately 30 retained backbones contained high-yield designs (Extended Fig. 5g). Before filtering, high-yield designs were distributed across approximately 50 backbones for several methods. Structural-confidence filtering therefore narrowed both the number of candidates and the range of backbones represented among useful designs.

### Hydrophobicity is associated with the coupling between folding stability and yield

Folding stability is often targeted to improve protein production^46,66^, but its relationship with monomeric yield remains unclear. We therefore asked whether increased stability consistently translates into higher monomeric yield and what determines the strength of this relationship. To address these questions, we measured folding free energies (ΔG) by cDNA-display proteolysis using the assay framework of Tsuboyama et al. (2023)^116^. Protein–cDNA fusions were exposed to a range of protease concentrations, and sequencing of surviving constructs was used to infer ΔG; more positive ΔG values indicate greater stability. The mutant collection included full scans of nine proteins, with single substitutions, deletions and alanine insertions, and selected single mutants spanning a broad range of previously measured stabilities, drawn from 56 additional proteins(Fig. 3a).

**Figure 3.**
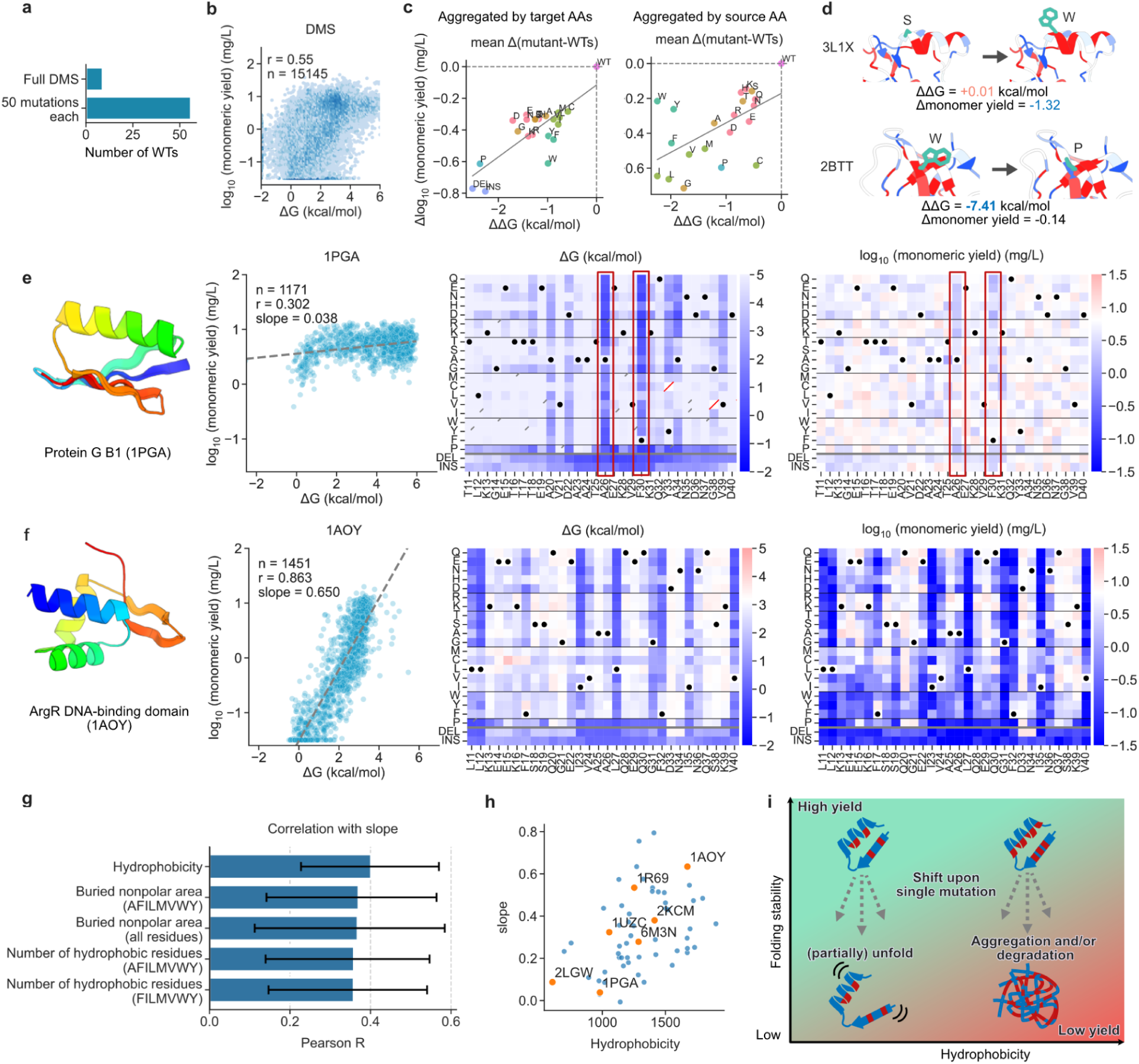
Protein hydrophobicity is associated with the slope of the relationship between folding stability and monomeric yield. **a,** Composition of the mutational analysis. Nine parental proteins were subjected to full scans containing all single substitutions, deletions and alanine insertions. Selected 50 single mutants of an additional 56 parental proteins were also analysed. **b,** Relationship between folding stability (ΔG, kcal/mol) and log₁₀-transformed monomeric yield (mg/L) across 15,145 variants (Pearson r = 0.55). **c,** Mean mutational effects on folding stability and monomeric yield, grouped by the target amino acid introduced by mutation (left) or the source amino acid in the wild-type sequence (right). ΔΔG and Δlog₁₀(monomeric yield) were calculated as mutant minus the corresponding wild-type value. Amino acids are indicated by one-letter codes; DEL and INS denote deletions and alanine insertions, respectively. The wild-type reference is located at the origin. Grey line represents linear regression line **d,** Representative mutations with contrasting effects on stability and monomeric yield. Top, 3L1X S30W has ΔΔG = +0.01 kcal/mol and Δlog₁₀(yield) = −1.32. Bottom, 2BTT W51P has ΔΔG = −7.41 kcal/mol and Δlog₁₀(yield) = −0.14. The affected side chains are highlighted in the structural models. **e,f,** Folding-stability and yield landscapes for protein G B1 (1PGA; e) and the ArgR DNA-binding domain (1AOY; f). Each panel shows, from left to right, a structural model, a scatter plot of log₁₀ monomeric yield versus ΔG, and heatmaps of ΔG and log₁₀ monomeric yield. Dashed lines in the scatter plots indicate linear fits. In the heatmaps, columns indicate sequence positions and rows indicate substitutions or deletion/insertion classes. Black dots indicate wild-type residues, red diagonal marks indicate unavailable values, and black diagonal marks indicate uncertain ΔG estimates (95% CI >0.5 kcal/mol). **g,** Pearson correlations between protein-specific slopes and hydrophobicity-related descriptors. Error bars show 95% percentile confidence intervals from 1,000 bootstrap resamples of parental proteins. **h,** Relationship between protein hydrophobicity and the slope of the log₁₀(yield)–ΔG relationship. Each point represents one protein; proteins with full DMS are highlighted in orange and labelled. **i,** Proposed model for hydrophobicity-dependent coupling between stability and recovery. Destabilizing mutations in proteins with relatively low hydrophobicity may lead to partially unfolded states without strongly reducing monomeric recovery. In more hydrophobic proteins, destabilization may instead promote aggregation and/or degradation, resulting in a larger decrease in monomeric yield. Arrows illustrate these hypothesized responses to single mutations.

Across the 15,145 variants shown in Fig. 3b, monomeric yield and ΔG were moderately correlated (Pearson’s r = 0.55), suggesting mutational effects differed between the two properties. Indeed, on average, substitutions to tryptophan markedly reduced monomeric yield with relatively small changes in folding stability (ΔG). Conversely, substitutions from tryptophan substantially reduced folding stability but had comparatively modest effects on monomeric yield (Fig. 3c; Extended Fig. 6). For example, 3L1X S30W changed ΔG by only +0.01 kcal/mol but reduced monomeric yield by ∼21-fold (1.32 in log10). Conversely, 2BTT W51P reduced ΔG by 7.41 kcal/mol but retained approximately 70% of the wild-type monomeric yield (a decrease of 0.14 in log₁₀ yield) (Fig. 3d). Similar but less pronounced trends were observed for substitutions involving tyrosine and phenylalanine. These asymmetric effects suggest that bulky hydrophobic residues contribute differently to folding stability and monomeric recovery depending on sequence context and the direction of substitution.

The relationship between ΔG and yield also differed among parental proteins (Fig. 3e,f; Extended Figs. 7–8). For example, protein G B1 (1PGA) yield changed relatively little across the measured stability range (Pearson’s r = 0.302; slope = 0.038), whereas the ArgR DNA-binding domain (1AOY) showed stronger coupling (r = 0.863; slope = 0.650). Even in 1PGA, substitutions at positions A26 and F30 that strongly reduced folding stability were accompanied by modest decreases in monomeric yield. Thus, a shallow ΔG–yield slope does not imply that yield is entirely independent of folding stability; rather, it indicates that changes in stability are associated with smaller changes in yield in some proteins than in others.

To investigate why the relationship between folding stability and monomeric yield differed among proteins, we examined protein features associated with the ΔG–yield slope and found positive associations with hydrophobicity-related descriptors (Fig. 3g; Extended Fig. 9). The strongest association among the displayed descriptors was with the sequence hydrophobicity score (Fig. 3h; Pearson’s r = 0.402). These associations indicate that destabilization tends to have a larger effect on monomeric recovery in more hydrophobic proteins within this collection. One possible explanation is that destabilization of more hydrophobic proteins may expose hydrophobic surfaces that promote aggregation or cellular degradation, thereby reducing monomeric recovery. In contrast, more hydrophilic proteins may remain soluble even when destabilized and partially unfolded. (Fig. 3i).

### Prediction of monomeric yield and prospective candidate selection

Our analyses showed that structure-prediction-based filtering provided only limited enrichment of high-yield designs and that the relationship between folding stability and monomeric yield varied with protein hydrophobicity (Figs. 2 and 3). These findings motivated us to test whether integrating multiple sources of sequence and structural information could improve yield prediction. Using the dataset generated by our workflow, we first established a baseline by examining individual associations of monomeric yield with folding stability, structure-prediction confidence and structural descriptors. Measured ΔG and ESM3ΔG-predicted stability^117^ showed the strongest correlations among the displayed features, each with Pearson’s r of approximately 0.45 (Fig. 4a; Extended Fig. 10a). Measured and ESM3ΔG-predicted folding stability showed similar correlations with monomeric yield, consistent with the high agreement between measured and predicted stability in both de novo designs and the mutant collection (Extended Fig. 10b). Structure-confidence and Rosetta-derived terms were also associated with yield, but no single feature fully explained the observed differences in monomeric yield.

**Figure 4.**
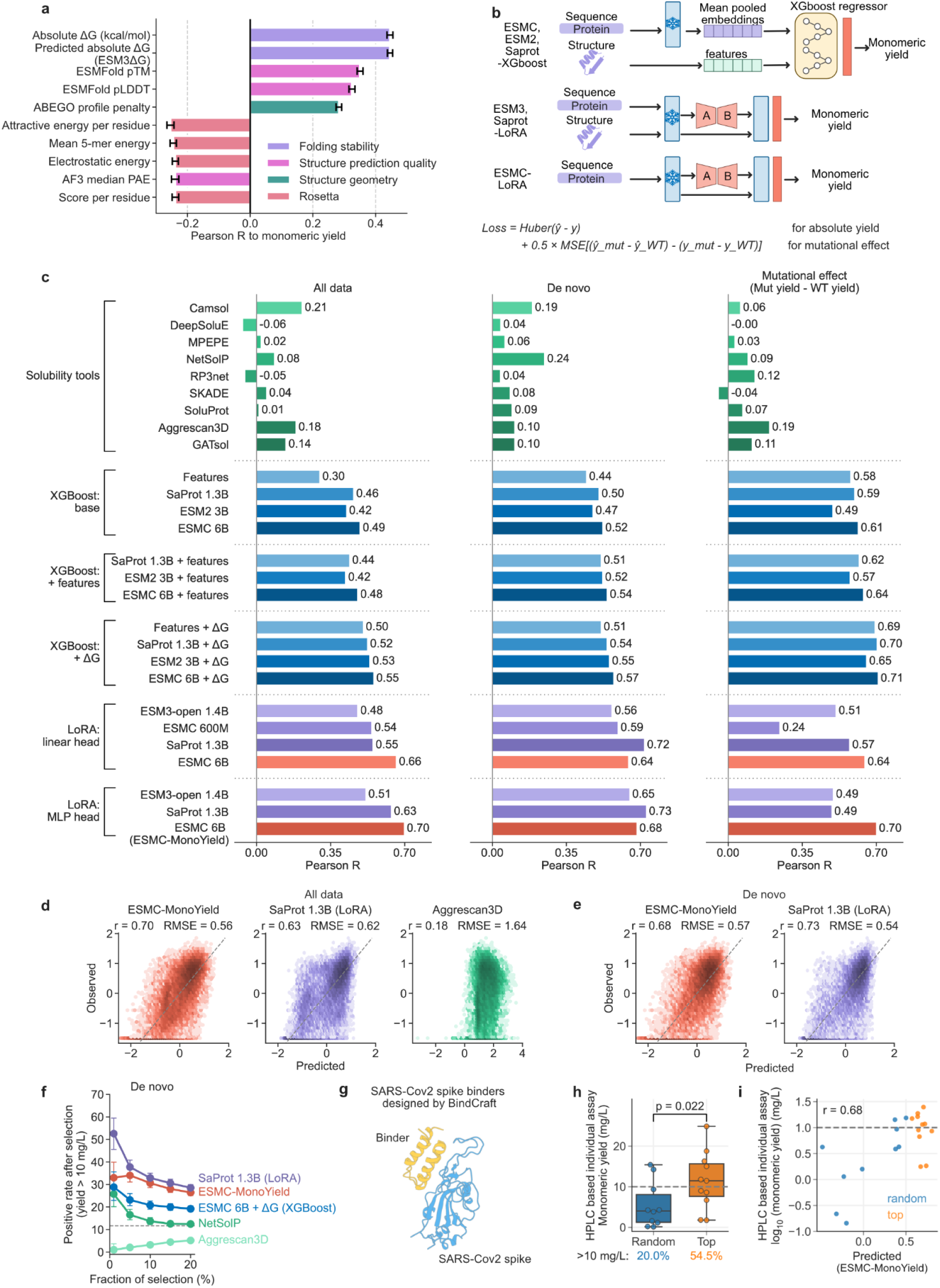
pLMs and structure-informed features enable prediction and prospective enrichment of high-yield proteins. **a,** Pearson correlations of measured or predicted folding stability, structure-confidence metrics, geometric descriptors and Rosetta-derived terms with log₁₀ monomeric yield. Correlations were calculated using unclipped log₁₀ monomeric yield values. Error bars show 95% bootstrap confidence intervals (1,000 resamples). Colours indicate feature categories. **b,** Modelling strategies. Top, frozen mean-pooled ESMC, ESM2 or SaProt embeddings and/or explicit descriptors were supplied to XGBoost; selected inputs also included predicted ΔG. Middle, ESM3 and SaProt were fine-tuned with LoRA using sequence and structural representations. Bottom, ESMC was fine-tuned using sequence alone. The displayed Huber-plus-mutational-effect loss describes the LoRA objective; the XGBoost objective is specified separately in Methods. **c,** Pearson correlations for trained models and existing predictors on the combined evaluation set (left), de novo designs (middle) and mutational effects (right). Mutational effects are mutant-minus-wild-type differences in log₁₀ monomeric yield. Sample sizes were n = 25,065, 19,327 and 5,735, respectively, for all methods except DeepSoluE, for which n = 19,592, 15,169 and 4,356. DeepSoluE did not return predictions for some sequences owing to tool-specific limitations and was evaluated on the subset with available predictions. Trained models were assessed using grouped cross-validation as described in Methods. **d,** Observed log₁₀ monomeric yield versus predictions from ESMC-MonoYield and SaProt 1.3B LoRA, or the AGGRESCAN3D score (left to right), for the combined evaluation set. The displayed Pearson correlations are 0.70, 0.63 and 0.18, respectively (n = 25,065 in each panel). RMSE values are 0.56, 0.62 and 1.64, respectively; the uncalibrated AGGRESCAN3D score has different units from yield, so its displayed RMSE is not comparable with the yield-model errors. Dashed diagonal lines denote y = x for calibrated yield predictions. **e,** Observed log₁₀ monomeric yield versus predictions from ESMC-MonoYield (left) and SaProt 1.3B LoRA (right) for de novo designs (n = 19,327 in each panel). Pearson correlations are 0.68 and 0.73, and RMSE values are 0.57 and 0.54, respectively. Dashed diagonal lines denote y = x. **f,** Enrichment of high-yield de novo proteins by predictor-based ranking. The percentage of proteins with monomeric yield >10 mg/L among the retained sequences is plotted against the fraction of sequences selected. Curves compare SaProt 1.3B LoRA, ESMC-MonoYield, ESMC 6B + ΔG XGBoost, NetSolP and AGGRESCAN3D. Error bars indicate pointwise 95% Wilson confidence intervals for the positive rates. The dashed horizontal line indicates the positive rate without selection. **g,** Representative predicted complex of a BindCraft-designed binder (gold) and SARS-CoV-2 spike (blue), illustrating the prospective validation set. **h,** Individually measured yields of BindCraft designs selected randomly (n = 10) or by ESMC-MonoYield (n = 11). Candidates had ipTM >0.8. Each protein was expressed and measured once. The fractions with yield >10 mg/L were 20.0% and 54.5%, respectively. P = 0.022 is from a one-sided Mann–Whitney U test of yield distributions, not a test of the two success proportions. Boxes show medians and interquartile ranges; whiskers extend to observations within 1.5 interquartile ranges. The dashed horizontal line marks a monomeric yield of 10 mg/L. **i,** Relationship between predicted yield and individually measured log₁₀ monomeric yield for the 21 prospectively tested binders included in the comparison (Pearson r = 0.68). Orange and blue points indicate top-ranked (n = 11) and randomly selected (n = 10) designs, respectively. The dashed horizontal line marks a monomeric yield of 10 mg/L.

We next developed models to predict monomeric yield from sequence and structural descriptors, pLM representations, or combinations of these inputs. We clustered sequences using MMseqs2^118^ at a 30% sequence identity threshold and assigned entire clusters to five cross-validation groups (Methods). We trained XGBoost^119^ regressors on mean-pooled pLM embeddings and sequence and structural descriptors, with or without predicted folding stability (ESMΔG; Cho et al., 2026^117^). We also fine-tuned ESMC, ESM3 and SaProt using low-rank adaptation (LoRA; Fig. 4b).

During fine-tuning, we jointly trained models to predict absolute log₁₀ monomeric yields of de novo designs and wild-type proteins, using Huber loss, and mutation-induced changes in log₁₀ yield, using mean-squared-error loss, with respective weights of 1.0 and 0.5 (Fig. 4b, Methods). Mutant absolute yields were not used as separate training targets. We assessed both yield predictions and predicted mutational effects on held-out data.

We first evaluated previously developed solubility and aggregation predictors^90,92,96,97,99,103,120–122^ as baselines, which achieved Pearson correlations of up to 0.21 with monomeric yield on the combined evaluation set (Spearman rho up to 0.17) (Fig. 4c, Extended fig. 10c, 11). XGBoost regressors trained on mean-pooled pLM embeddings from ESM-2, ESMC, and SaProt^33,123,124^ achieved Pearson correlations of 0.3-0.49 in the combined evaluation set. Adding explicit sequence and structural descriptors to the pLM embeddings provided little overall improvement in XGBoost performance (Fig. 4c, Extended fig. 11). Adding predicted folding stability (ΔG) improved predictions by up to 0.20 in Pearson correlations, consistent with the association between stability and yield (Fig. 4a,c).

LoRA fine-tuning further improved overall yield prediction, achieving a Pearson correlation of 0.70 on the combined evaluation set. Replacing linear regression heads with multilayer perceptron (MLP) heads improved performance on the combined evaluation set and de novo subset (Fig. 4c). SaProt 1.3B and ESMC 6B with LoRA and MLP heads were among the best-performing models: SaProt performed better than ESMC 6B on de novo yields (r = 0.73 versus 0.68), whereas ESMC 6B performed better on mutational effects (r = 0.70 versus 0.49). Given its balance of performance across these tasks, we selected ESMC 6B with LoRA and an MLP head as our primary predictor and named it ESMC-MonoYield (Fig. 4c-e).

To assess how effectively the models prioritize high-yield de novo proteins, we compared the fraction of designs with monomeric yields >10 mg/L among their top-ranked candidates (Fig. 4f, Extended Fig. 12a). ESMC-MonoYield and SaProt fine-tuned with LoRA and an MLP head were the best-performing models in this retrospective evaluation.

To prospectively evaluate ESMC-MonoYield in a practical protein-design setting, we generated SARS-CoV-2 spike binders using BindCraft^8^ (Fig. 4g). Among 226 candidates with interface predicted TM scores (ipTM) >0.8, we selected 11 candidates from among the top-ranked candidates based on ESMC-MonoYield predictions and 10 at random. All 21 proteins were individually expressed, purified and quantified by HPLC–SEC (Extended Fig. 12b). Six of 11 model-selected designs (54.5%) and two of 10 randomly selected designs (20.0%) achieved monomeric yields >10 mg/L (Fig. 4h). Across the 21 binders, ESMC-MonoYield predictions correlated more strongly with measured log₁₀ yields (Pearson’s r = 0.68) than those from SaProt 1.3B LoRA, AGGRESCAN3D and NetSolP (r = 0.59, −0.22 and 0.32 respectively; Fig. 4i; Extended Fig. 12c). These results support the use of ESMC-MonoYield to prioritize high-yield binders among candidates already filtered by BindCraft, with performance validated by individual expression and HPLC–SEC measurements independently of the pooled MS-based workflow.

For mutational effects, XGBoost models incorporating predicted ΔG achieved high correlations with measured changes (Fig. 4c). However, this did not translate into a clear advantage in selecting yield-improving mutations (Extended fig. 12d). Using the criterion of at least a fivefold yield increase relative to wild type, none of the learned models consistently outperformed the other methods across selection fractions; AGGRESCAN 3D and NetSolP gave comparable or greater enrichment at selection fractions of 5–20% (Extended Fig. 12d). Thus, although the models effectively prioritized high-yield de novo designs, they offered no consistent advantage in identifying yield-improving mutations.

We also examined PROSS^66^-designed variants independently of predictor ranking. Using a twofold-change threshold for yield, 27.4% of variants improved and 10.6% decreased monomeric yield relative to their wild types (Extended Fig. 12e). Across PROSS-designed variants, changes in folding stability (ΔΔG) showed no clear association with changes in monomeric yield, suggesting that factors beyond folding stability may contribute to yield improvements (Extended Fig. 12e).

### Sequence determinants and prediction of nanobody monomeric recovery

Recent advances in computational antibody design have enabled the generation of functional nanobodies targeting specified epitopes^43,125^. However, evaluating their production yields and recovery as monomers remains important for their practical use. We therefore extended our pooled assay to designed nanobodies, quantifying both monomeric and dimeric fractions to identify sequence features associated with monomeric yield and the monomer/dimer yield ratio.

To this end, we designed CDR2 and CDR3 of the humanized NbBCII10 (FGLA) scaffold^126^ using RFantibody and BoltzGen^9,43^ against PD-1 or PD-L1 (Fig. 5a) while keeping the original sequence of CDR1 due to length constraints in DNA oligonucleotide pool synthesis. CDR2 contained six residues and CDR3 ranged from five to thirteen residues; the framework and CDR1 were fixed. The library contained 650 designs for each target–method combination, giving 2,600 designs, each assigned 15 peptide barcodes. Monomeric and dimeric SEC fractions were collected separately and processed through the same barcode-quantification workflow (Fig. 5b).

**Figure 5.**
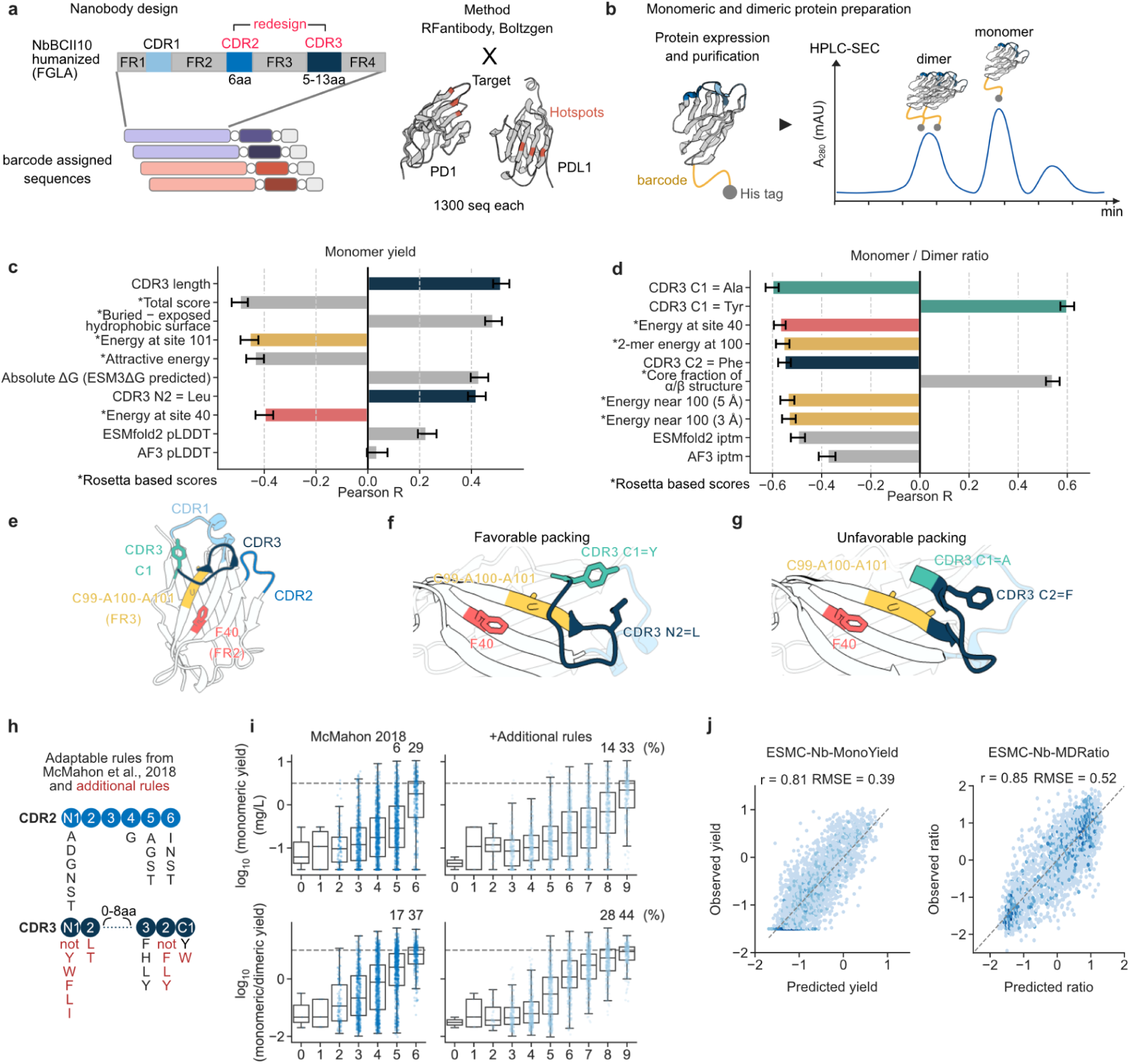
Sequence rules and nanobody-specific models guide selection for improved monomeric recovery. **a,** Design of the nanobody library. CDR2 and CDR3 of a humanized NbBCII10 (FGLA) scaffold were redesigned using RFantibody and BoltzGen against PD-1 or PD-L1, with target hotspot residues specified during design. CDR2 contained six residues, whereas CDR3 lengths ranged from five to thirteen residues. The experimental library contained 650 designs per target–method combination, for 2,600 designs in total, with 15 peptide barcodes assigned to each design. **b,** Measurement of nanobody monomeric and dimeric recovery. Following expression and affinity purification, monomeric and dimeric fractions were collected separately by HPLC–SEC and quantified using peptide barcodes. The monomer/dimer yield ratio was calculated as the recovered monomeric yield divided by the recovered dimeric yield. **c,d,** Pearson correlations between selected sequence and structural features and log₁₀ monomeric yield (c) or log₁₀ monomer/dimer yield ratio (d). CDR3 N1/N2 and C1/C2 denote the first two residues counted from the N and C termini of the loop, respectively. Asterisks denote Rosetta-derived scores; error bars indicate 95% bootstrap confidence intervals (1,000 resamples). **e–g,** CDR3–framework packing associated with nanobody monomeric yield and monomer/dimer yield ratio. **e,** Structure predicted by AlphaFold3 from a nanobody sequence in the library used in this study, showing CDR1 (light blue), CDR2 (blue) and CDR3 (dark blue). CDR3 C1 is highlighted in teal, F40 in FR2 in pink, and C99–A100–A101 in FR3 in yellow. Framework residues are numbered sequentially along the scaffold. **f,g,** Representative models showing favourable packing with Tyr at C1 and Leu at N2, associated with higher monomer/dimer yield ratio and monomeric yield, respectively (f), and unfavourable packing with Ala at C1 and Phe at C2, both associated with lower monomer/dimer yield ratios (g). The same colour scheme is used throughout. **h,** Sequence preferences adapted from McMahon et al. (2018)^111^, with additional or modified preferences in red. The original scoring scheme contains six rules. The expanded scheme contains nine rules, including a modified C1 rule and three added rules; the two scores are therefore not strictly nested. N1 and C1 are the first residues counted from the corresponding CDR terminus. **i,** Distributions of log₁₀ monomeric yield (top) and log₁₀ monomer/dimer yield ratio (bottom) by the number of satisfied rules. The six-rule and nine-rule schemes are shown at left and right, respectively. Points represent individual nanobodies. Boxes show medians and interquartile ranges, and whiskers extend to observations within 1.5 interquartile ranges. Percentages above the selected groups indicate the fractions meeting the corresponding positive criterion: monomeric yield ≥3.16 mg/L or monomer/dimer ratio ≥10. **j,** Out-of-fold predictions from independently fine-tuned ESMC 6B LoRA models for monomeric yield (left) and the monomer/dimer yield ratio (right). Dashed lines denote y = x,.

BoltzGen designs tended to have lower monomeric yields but higher monomer/dimer yield ratios than RFantibody designs (Extended Fig. 13a). We also assessed target-binding enrichment by cDNA display. Enrichment was modest and tended to correlate negatively with monomeric yield and the monomer/dimer ratio, particularly for PD-L1-targeting designs (Extended Fig. 14a). These results were treated as exploratory because monomeric and dimeric yields varied broadly across designs (Extended Fig. 13b) and enrichment in cDNA display was limited.

In the global feature analysis, stability-related metrics including predicted ΔG and Rosetta-derived energetic terms were associated with monomeric yield (Fig. 5c). Conversely, homodimer-confidence metrics such as ipTM negatively correlated with the ratio of monomeric to dimeric yield. Notably, ESMFold2 confidence metrics showed a stronger relationship with both monomeric yield and the yield ratio than those from AlphaFold3. Position-resolved analyses identified residue preferences near the CDR3 termini (Extended Fig. 13c–e), with N1/N2 and C1/C2 denoting positions counted from the N and C termini, respectively. Tyr at C1 was positively associated with the monomer/dimer ratio, whereas Ala at C1 and Phe at C2 were negatively associated; Leu at N2 was positively associated with monomeric yield (Fig. 5c,d). These CDR3 positions lie near invariant framework residues F40 in FR2 and C99–A100–A101 in FR3 (Fig. 5e). Indeed, a representative model with favourable yield behaviour, containing Tyr at C1 and Leu at N2, showed close packing against these framework residues (Fig. 5f), whereas a model with a lower monomer/dimer ratio, containing Ala at C1 and Phe at C2, showed a more extended CDR3 conformation with fewer contacts with these residues (Fig. 5g). This interpretation is consistent with prior work implicating framework–CDR3 interactions involving F40 (F42 in IMGT numbering) in VHH binding affinity and thermal stability^127^.

We next asked whether previously proposed sequence rules could help identify nanobodies with higher monomeric yield. Of the rules proposed by McMahon et al. (2018)^111^, we evaluated the six applicable to our nanobody dataset (Fig. 5h). Among designs satisfying all six rules, 29% achieved monomeric yields > ∼3 mg/L, compared with 6% and 3% of designs satisfying five and four rules, respectively (Fig. 5i). We then modified one of the previous rules and added three new rules on sequence preferences identified in our analysis, yielding a nine-rule scheme. Among designs satisfying all nine rules, 33% achieved yields > ∼3 mg/L and 44% achieved monomer/dimer yield ratios ≥10, compared with 29% and 37%, respectively, for the six-rule scheme. These results suggest modestly improved enrichment, although independent validation is needed because the additional preferences were derived from the same dataset.

When applied to nanobodies, the de novo/DMS-trained ESMC-MonoYield model showed only a weak correlation with measured monomeric yield (Pearson’s r = 0.23), and its predictions spanned a substantially narrower range than the observed yields (Extended Fig. 14b). These results indicate limited transfer of the model to the nanobody design space. This encouraged us to independently fine-tune the pretrained ESMC 6B model for monomeric yield and for the monomer/dimer yield ratio, naming the resulting models ESMC-Nb-MonoYield and ESMC-Nb-MDRatio. These models were initialized from pretrained ESMC, rather than from ESMC-MonoYield. In the existing nanobody-specific grouped cross-validation, observed and predicted values were correlated for both yield (Pearson’s r = 0.81; RMSE = 0.39) and ratio (r = 0.85; RMSE = 0.52; n = 2,589; Fig. 5j). Among the top-ranked 5%, more than 50% of sequences selected by ESMC-Nb-MonoYield had monomeric yields > ∼3 mg/L whereas more than 60% of sequences selected by ESMC-Nb-MDRatio had monomer/dimer ratios ≥10 (Extended Fig. 14c). We also evaluated the OmniLib models, which predict nanobody folding fitness from CDR sequences^128^, to test whether existing folding predictors could inform monomeric recovery under our production conditions. These models showed weaker associations with monomeric yield (r = 0.24–0.27) and the monomer/dimer yield ratio (r = 0.28–0.31) among 2,389 scorable designs (Extended Fig. 14d), suggesting that predicted folding fitness does not necessarily reflect monomeric yield.

## Discussion

Selecting among computationally designed sequences requires an estimate of how much protein can be recovered for experimental characterization. Our study makes purified monomeric yield a directly measured and predictable criterion for this decision. However, our measurements were obtained using a single production workflow comprising cytosolic expression in a BL21(DE3)-derived Escherichia coli host, overnight autoinduction at 37°C, His-tag affinity purification and size-exclusion chromatography (SEC). Other temperatures, hosts and production conditions were not tested, although host strain and cultivation conditions can substantially affect recombinant protein production^3,50^. Although comparison with 48 individually expressed proteins supported the pooled estimates (r > 0.75; Fig. 1h) and shared proteins showed high correlated monomeric yield across 15 sublibraries (r > 0.92; Fig. 1g, Extended Fig. 2a), differences in clone abundance, expression burden and purification recovery may contribute to discrepancies between MS-based estimates and individual production yields. The spike-in standard was added after collection of the monomeric SEC fraction and therefore did not directly correct variation introduced during cultivation or purification. Thus, even after calibration, pooled yield estimates should not be considered interchangeable with yields measured in individual cultures.

The scope of these measurements also defines the context in which our predictive models should be applied. ESMC-MonoYield was trained on small protein domains and is best supported for ranking candidate sequences in the same production workflow and for de novo designed proteins that function as monomers. In our deep mutational scanning analyses, however, ESMC-MonoYield did not show a clear advantage over the existing methods tested, including the structure-based aggregation predictor AGGRESCAN3D^103^ and language-model-based protein solubility predictor NetSolP^97^, for identifying single substitutions that improve monomeric yield. Because comprehensive mutational measurements covered only nine parental proteins and only ∼0.43% of them showed a >5-fold increase in monomeric yield, training data may contain too few examples of yield-increasing mutations for the model to reliably identify such mutants and broader coverage will be needed to assess and improve this application.

In addition, because the de novo designs in our dataset were generated without conditioning on function, functional proteins such as binders of proteins, RNA, DNA or small molecules, as well as enzymes, may occupy sequence and structural spaces that are underrepresented in the training data. Predictive performance across these broader sequence and structural spaces will require further evaluation and may benefit from additional measurements and training. Indeed, transfer of ESMC-MonoYield to nanobodies produced a compressed range of predictions (Extended Fig. 14b), motivating the development of ESMC-Nb-MonoYield and ESMC-Nb-MDRatio. The nanobody models were evaluated only within the humanized NbBCII10 (FGLA) framework and the CDR configurations represented in our libraries (Fig. 5a,j), and their performance on other frameworks or antibody formats remains untested.

Despite these limitations, this study quantified purified monomeric yield for 39,792 protein constructs under a common production workflow, enabling systematic analysis of monomeric yield and the development of predictive models. Building on peptide barcoding for pooled protein characterization^24,25^, our assay assigns a distinct peptide barcode to each variant, enabling discrimination of closely related variants that are difficult to resolve using target-derived peptides. We used this capability to measure the effects of single amino acid substitutions on monomeric yield in deep mutational scanning (DMS) libraries (Fig. 3; Extended Fig. 7). The resulting data allowed us to directly compare monomeric yield with folding stability, revealing that the association between these properties varied with parental protein hydrophobicity. Improvements in peptide-barcode detectability, MS sensitivity and throughput could extend coverage to more sequences and conditions, increasing both the scale and diversity of future datasets.

Our analysis of the de novo protein dataset showed that widely used strategies, including filtering by structural confidence metrics and using SolubleMPNN, provided limited advantages in monomeric yield under the conditions tested (Fig. 2). Differences among inverse-folding methods, including ProteinMPNN, CarbonDesign, HyperMPNN and Boltz-IF, were relatively modest, and SolubleMPNN showed no clear advantage over standard ProteinMPNN. These findings highlight the importance of directly benchmarking design methods through wet-lab experiments that directly measure the intended outcome, rather than relying solely on *in silico* assessments.

Beyond comparing design methods and filtering strategies, the dataset provides a basis for predicting and ranking monomeric yield at the level of individual candidate sequences. To this end, we developed ESMC-MonoYield by fine-tuning a pretrained pLM using low-rank adaptation^129^. ESMC-MonoYield supported prospective selection of higher-yielding candidates from a pool of BindCraft designs^8^ preselected using structural confidence metrics (Fig. 4g–i). Using only amino acid sequences as input, ESMC-MonoYield predicts monomeric yield in ∼0.3 s per sequence on a single consumer-grade GPU with reduced-precision inference, while maintaining close agreement with full-precision predictions (r = 0.9998, data not shown). Its low computational cost and minimal input requirements facilitate integration into de novo protein-design workflows as an additional selection criterion alongside structural confidence metrics such as ipTM and pLDDT.

Future work should expand both the protein space and the production workflows represented in the data. Measurements across more diverse backbones, protein families, hosts, temperatures and purification methods should enable more accurate prediction of monomeric yield across diverse protein families, better identification of yield-increasing mutations, and could also help optimize production conditions for individual proteins. A further goal is to improve de novo protein design by incorporating yield prediction into structural design, sequence generation, or integrated design and filtering workflows such as BindCraft^8^, Proteina-Complexa^44^ BoltzGen^9^ and RFD4-Proteina^130^. Extending our approach beyond yield, large-scale measurements of oligomeric state^13,25^, conformational fluctuations^131^, aggregation under defined stresses^132^ and enzyme kinetics in microfluidic assays^133^ could be used to fine-tune ESM-C^33^, ESM3^7^ or newer pretrained models to predict complementary properties. Combining these predictors with repeated cycles of design, experimental measurement and model updating, together with independent structural and functional evaluation, could enable more rational and efficient de novo protein design. This framework could ultimately support autonomous design agents that generate and prioritize design candidates in silico to achieve high experimental success rates within a single design cycle.

## Methods

### Protein library design

The experimental collections included 37,434 entries: 19,250 inverse-folding designs, 15,584 mutant-collection entries comprising full scans, selected single mutants and PROSS variants, and 2,600 designed nanobodies (Fig. 1i). Constructs were distributed among sublibraries for pooled expression and barcode quantification (Extended Fig. 1b). A separate sublibrary tested synonymous coding-sequence variation. Collection sizes, analysis-specific sample sizes and the number of distinct amino acid sequences are distinguished in the corresponding analyses.

For inverse-folding benchmarking, 25 sequences were generated for each of 70 backbone structures using 11 methods or parameter settings: ProteinMPNN at sampling temperatures of 0.1 and 0.3, ProteinMPNN-Cα, SolubleMPNN, HyperMPNN, ESM-IF1, ESM3, Frame2seq, CarbonDesign, TIMED-Design and Boltz-IF^5,7,9,26–30,115^. The backbone set comprised 58 de novo-designed structures and 12 natural domains, spanning all-α, α/β and all-β architectures. Sequences were padded to 71 aa with SAGG-repeat linkers at the N and C termini before synthesis.

ProteinMPNN used the v_48_020 checkpoint at T = 0.1 or 0.3. ProteinMPNN-Cα used Cα-only input, SolubleMPNN used soluble-protein weights at T = 0.1, and HyperMPNN used v48_020_epoch300_hyper at T = 0.1. These runs used seed 37 and generated 25 sequences per backbone, without fixed positions, amino-acid biases, PSSMs or tied positions. ESM-IF1 used esm_if1_gvp4_t16_142M_UR50 at T = 0.3; ESM3 used esm3_sm_open_v1 at T = 0.3; Frame2seq used a three-model ensemble (package 0.0.8) at T = 0.3; CarbonDesign used carbondesign_default at T = 0.4; and TIMED-Design used version 1.0.1 at T = 0.14 with seed 42. No explicit random seed was set for ESM-IF1, ESM3, Frame2seq or CarbonDesign. Boltz-IF was run through the BoltzGen inverse-folding mode with boltzgen1_ifold, 200 sampling steps, three recycling steps and backbone-only whole-chain design, without fixed residues. Sampling temperatures, where applicable, were chosen to obtain mean pairwise identities of approximately 0.65–0.75 among sequences generated for a backbone.

Full mutational scans were constructed for nine proteins: 1AOY, 1PGA, 1R69, 1UZC, 2BTT, 2KCM, 2LGW, 3L1X and 6M3N. The parental sequences were 56–69 residues long. Each scan included all 19 amino acid substitutions at each position, single-residue deletions and alanine insertions before each position and after the C terminus. The design manifest contained 10,944 substitution entries, 576 deletions and 585 insertions.

For 56 additional parents, selected single substitutions were drawn from Tsuboyama et al. (2023)^116^, Dataset 1 dated 16 April 2023. Candidate substitutions were restricted to unambiguous single mutants with ΔG between −1 and 5 kcal/mol. For each parent, the range was divided into 49 equal-width bins, one mutant was sampled from each occupied bin, and any shortfall was filled by additional random sampling to obtain 49 mutants (seed 42). The corresponding wild type was included separately.

The PROSS collection comprised 558 previously designed variants from 112 parental proteins, with one to nine designs per parent, obtained from Tsuboyama et al. (2023)^116^.

Wild-type sequences for the 177 parents were included together with four additional reverse translations per parent generated using human, mouse, yeast or E. coli codon-usage tables. These synonymous constructs provided repeated wild-type measurements. Repeated amino acid sequences can arise from synonymous coding constructs and from deletions at adjacent identical residues. Shared control proteins were encoded in the sublibraries indicated in Extended Fig. 1b; the nanobody sublibrary did not contain this shared control panel. Spike-in proteins were supplied as a separately prepared standard after SEC.

### Nanobody library design

Nanobodies targeting PD-1 or PD-L1 were designed with RFantibody and BoltzGen^9,43^ on the humanized NbBCII10 (FGLA) scaffold^126^, using h-NbBCII10.pdb distributed with RFantibody. Framework positions refer to sequential scaffold numbering rather than Kabat, Chothia or IMGT numbering. In the parental scaffold, CDR1 occupied positions 26–35, CDR2 positions 55–60 and CDR3 positions 102–117. The framework and CDR1 sequence GGSEYSYSTF were fixed. CDR2 contained six residues and CDR3 was varied from five to thirteen residues. Loop-relative labels N1, N2 and C1, C2 count from the N and C termini of each CDR, respectively.

The PD-1 target was the murine extracellular domain from PDB 3BIK, chain B, residues 33–148, extracted and renamed chain T. The PD-L1 target was a model of the human extracellular-domain expression construct, also supplied as chain T. Design hotspots were residues T64, T126, T129 and T133 for PD-1 and T40, T99 and T107 for PD-L1 in source-structure numbering. For BoltzGen, PD-1 hotspots were converted to chain-local positions 32, 94, 97 and 101; PD-L1 hotspot positions remained 40, 99 and 107.

RFantibody version 1.0.0 (commit e0b2188c) was run with the antibody configuration of RFdiffusion and the RFdiffusion_Ab checkpoint. Loops were specified as H2:6 and H3:5–13, with 50 diffusion steps, final_step = 48 and deterministic sampling. The design procedure generated 5,000 backbones per target. Each backbone was subjected to two ProteinMPNN sequence-design passes with one sequence per pass and redesign restricted to H2 and H3. Other settings were the interface-design script defaults: sampling temperature 0.000001, omit_AAs = CX, augment_eps = 0, full protein features, 48 connections and ProteinMPNN_v48_noise_0.2.pt. RoseTTAFold2 predictions were generated within the pipeline but were not used for library selection.

BoltzGen version 0.1.2 was run with the nanobody-anything protocol, num_designs = 10,000 and budget = 10,000 per target. The design stage used boltzgen1_diverse.ckpt and boltzgen1_adherence.ckpt, switching checkpoints halfway through sampling; inverse folding and folding used boltzgen1_ifold.ckpt and boltz2_conf_final.ckpt. CDR2 was redesigned at positions 55–60, and CDR3 was rebuilt as a variable-length insertion at position 102. Both loops were hidden from structural conditioning while the remaining framework was retained. Design used 500 sampling steps, three recycling steps and ten diffusion samples at multiplicity 1,000. Inverse folding used 200 sampling steps, with cysteine disallowed.

Candidate nanobody–target complexes were re-predicted with AlphaFold3 using the Kuhlman-Lab fork (commit 91552c32f9c7f5d1911a497ea84a3fb7a8484e9f; version 3.0.0) and official model parameters. Interface and structural-confidence metrics were extracted from the confidence outputs. Cα RMSD relative to the design model was evaluated for the nanobody and separately for CDR2 and CDR3. Identical CDR2–CDR3 sequence pairs were collapsed to one representative. Candidates were retained if all three RMSDs were <3 Å. For each target–method combination, the ipTM range was divided into 50 equal-width bins and 13 designs were sampled from each bin (seed 42), giving 650 designs per combination and 2,600 in total. No additional threshold was applied to ipTM, pTM, pLDDT, PAE or the clash flag. Each selected design received 15 peptide barcodes.

### Peptide barcode selection and assignment

Candidate peptide barcodes from Feldman et al. (2025)^25^ were screened as C-terminal fusions to mGreenLantern. The screen included 70,116 sequences from the supplied barcode collection. Two samples contained the same nominal 1:1 mixture of two subpools: one was mixed after separate expression and purification, and the other was mixed at the plasmid stage before pooled expression. Signals were compared as log₁₀(intensity + 1). Barcodes were retained if detected in both preparations, if the first-preparation intensity lay between 10⁴ and 10⁷ with the corresponding regression-mapped window applied to the second preparation, and if the perpendicular distance from the fitted line was <0.4 log₁₀ units. Of 46,835 barcodes detected in both samples, 46,124 met all criteria (Fig. 1b).

Two pilot libraries containing the same 1,038 proteins, each represented by 40 independently assigned barcodes, were cloned, expressed and purified in parallel. Missing intensities were set to zero, 10⁴ was added to each barcode intensity, and protein-level signals were computed as the log₁₀ of the arithmetic mean. Subsets of 5–40 barcodes in increments of five were sampled without replacement within each protein and sublibrary. Pearson correlations between the two libraries were computed for each subsample, including the monomeric-fraction comparison in Extended Fig. 1a.

For subsequent libraries, each construct was normally assigned 15 barcodes. The retained pool was divided into 15 equal-count strata based on mean signal intensity, and each construct received one barcode from each stratum. Assignment used independent sublibrary-specific shuffles. Barcodes were unique within a sublibrary but could be reused across sublibraries. Spike-in barcodes were reserved and excluded from the experimental-construct assignments.

Barcodes were expressed as C-terminal fusions. The mGreenLantern construct contained a GGSGGSL linker, a 15-residue cassette consisting of lysine, a 9–14-residue barcode ending in arginine and serine-rich padding, followed by GGSGGSG and His₆. Protein-library constructs contained the padded protein, the lysine–barcode–padding cassette, the vector junction and His₆. The restriction-cloned constructs carried the XhoI-encoded Leu-Glu junction, whereas the nanobody construct used a GSGGS junction. Lys-C cleaved before the barcode cassette, and subsequent trypsin digestion released the arginine-terminated barcode peptide from the captured His-tagged fragment.

Sequence-to-barcode mapping tables specify the construct identifier, amino acid sequence, coding sequence, padding, sublibrary, barcode peptide and coding sequence, intensity stratum and synthesized oligonucleotide.

### DNA synthesis and library cloning

Amino acid sequences were reverse-translated with DNAWorks version 3.2.4 using E. coli codon usage^134^; no explicit CAI target range was imposed. Oligonucleotide pools were synthesized by Twist Bioscience. Each sublibrary was flanked by a unique pair of 15-nucleotide adapters for selective amplification. Restriction-cloned oligonucleotides contained, in order, the 5′ adapter, NdeI site, padded protein coding sequence, barcode cassette, XhoI site and 3′ adapter. The nanobody pool encoded variable loop regions and barcodes for assembly into the scaffold vector.

Adapter-specific PCR was used to amplify each sublibrary from the synthesized pool. Cycle numbers were chosen by pilot quantitative PCR with SsoAdvanced Universal SYBR Green Supermix (Bio-Rad) to prevent overamplification. Preparative PCR used KOD One PCR Master Mix (TOYOBO). Mutant, synonymous-codon and inverse-folding sublibraries were amplified in 400-µL reactions divided into 50-µL aliquots, pooled, column-purified and eluted in 30 µL water. Products were checked by MultiNA microchip electrophoresis (Shimadzu). Nanobody inserts underwent two PCR rounds; the nested 200-µL second reaction used a 100-fold dilution of the unpurified first-round product.

Mutant, synonymous-codon and inverse-folding inserts were cloned into pTF3-MDH by NdeI/XhoI (NEB) restriction ligation. Inserts and vector were digested in CutSmart buffer overnight at 37 °C; inserts were column-purified and vector was gel-purified. Ligation used 250 ng of each DNA, T4 DNA ligase and 5% PEG (Thermo Fisher Scientific) in 50 µL for 1–2 h at 22 °C, followed by column purification and ethanol precipitation with glycogen. Nanobody inserts were assembled into PCR-linearized, DpnI (TaKaRa)-treated pTF3-nanobody-vector_v2 with NEBuilder HiFi DNA Assembly Master Mix (New England Biolabs), using 250 ng insert and 600 ng vector for 1 h at 50 °C, followed by ethanol precipitation.

Library DNA was introduced into 50 µL of BL21-derived E. cloni electrocompetent cells (Lucigen) by electroporation at 1.8 kV with a 4.5-ms pulse and recovered in 1 mL SOC for 1 h at 37 °C. Serial dilutions were plated on LB–kanamycin agar to assess transformant coverage. The remaining recovery culture was expanded overnight in 100 mL LB–kanamycin at 37 °C. Plasmid DNA was isolated from 5 mL culture with the Plasmid Mini Kit (FastGene), and glycerol stocks were stored at −80 °C. The library-containing cultures were used for expression without another transformation step.

Plasmid-library representation was measured by Illumina NextSeq 1000/2000 sequencing with a P1 flow cell and 150-bp paired-end reads. Miniprepped plasmids were amplified with sublibrary-specific primers, treated with DpnI and purified with Sera-Mag carboxylate-modified beads (Cytiva) at a 0.9:1 bead-to-DNA volume ratio. A second PCR added Illumina P5 and indexed P7 adapters. PCR cycles were selected by pilot qPCR. Indexed products were checked by MultiNA, pooled, ethanol-precipitated and gel-purified before sequencing.

Demultiplexed FASTQ files (one sample per sublibrary; NextSeq run 019, January 2026, for the lib77 sublibraries and run 021, June 2026, for the lib82 sublibraries; 2.2–5.6 million reads per sample) were analysed using read 2 only, which reads each construct from the His6/XhoI end of the insert. Read 2 was trimmed at its 5′ end with cutadapt 5.1 (-g, maximum error rate 0.10), removing the constant adapter GTGGTGGTGGTGCTCGAG (the His6–XhoI junction) together with any upstream bases; 98.0–98.9% of reads carried this adapter. For the lib82 nanobody sublibrary, the adapter ACTTCCACCGCTTCC (the GSGGS-linker junction) was used instead and was detected in 65.5% of reads. Trimmed reads were reverse-complemented to the sense orientation and converted to FASTA with seqkit.

For each design–barcode entry, the reference window comprised the last 95 nt (lib82 muts-1, codon-1 and denovo-8 sublibraries), 133 nt (lib77 sublibraries) or 130 nt (lib82 nanobody sublibrary) of the ordered insert upstream of the XhoI site, or upstream of the GSGGS linker for the nanobody sublibrary. This window spans the 15-residue barcode cassette and the 3′ end of the coding sequence and therefore distinguishes the individual barcodes of a design. A read was counted only if its full sequence was identical to the reference window, with no mismatches allowed; counts were summed per window. Counts per million (CPM) were calculated within each sublibrary as the count for each entry divided by the total matched count for that sublibrary, multiplied by 10⁶. CPM values were used for the barcode-level representation filter described under “Calibration and calculation of monomeric yield”.

Individual reference constructs were obtained as clonal synthetic genes from Twist Bioscience. PCR-amplified inserts were gel-purified, NdeI/XhoI-digested and ligated into pTF3 before transformation into chemically competent E. coli DH5α. Colonies were checked by PCR with KT45_pET_seq_fw (5′-CCCGCGAAATTAATACGACTCACTATAG-3′) and KT46_pET_seq_rv_T7ter_rv (5′-GCCCCAAGGGGTTATGCTAG-3′), followed by Sanger sequencing. Verified plasmids were transformed into E. cloni BL21(DE3) for expression.

### Synonymous coding sequence library

A synonymous-coding-sequence library was designed for 50 proteins. Candidate parents were drawn from previously measured de novo designs after filtering for structural agreement and confidence with AlphaFold3. The measured log₁₀ yield range was divided into 50 equal-width bins, and four designs were sampled per bin to obtain 200 candidate parents spanning the yield distribution.

Each parent was padded to 71 residues with SAGG-repeat linkers, giving a 213-nucleotide protein coding region. DNAWorks version 3.2.4 generated 50 alternative coding solutions per parent for each of 11 codon-usage tables, with repeat = 8. BamHI, NdeI, XhoI, NheI and BsaI sites and homopolymeric A, C, G or T runs were forbidden. The tables represented E. coli Class II, Saccharomyces cerevisiae, Pichia pastoris, human, mouse, rat, Drosophila melanogaster, Xenopus laevis, Caenorhabditis elegans, Arabidopsis thaliana and Sulfolobus tokodaii. The last two tables were obtained from the Kazusa codon-usage database and converted to GCG format. This generated 110,000 candidate coding sequences.

CAI and tAI were calculated on the padded coding region using codonbias version 0.3.5, following Sharp and Li (1987)^114^ and dos Reis et al. (2004)^113^. CAI weights used the distributed reference set of 274 highly expressed E. coli UTI89 coding sequences (NC_007946.1), with genetic_code = 1, k_mer = 1, ignore_stop = True and pseudocount = 1. For tAI, tRNA gene counts were obtained from the GtRNAdb annotation of E. coli K-12 MG1655. Undetermined isotypes, pseudogenes and truncated genes were excluded, leaving 86 genes and 39 anticodons. Weights were computed with the dos Reis wobble coefficients and prokaryotic parameters. Sequence scores were geometric means of per-codon weights.

For each parent, the 550 candidate CAI values were divided into 49 equal-width bins and one sequence was sampled from each occupied bin (seed 42). Parents were eligible if at least 35 bins were occupied and candidate CAI values spanned ≤0.4 to ≥0.75. One eligible parent was selected from each of the 50 yield bins. The final library contained 49 new synonymous variants and the original coding sequence for each parent, giving 2,500 constructs. Among analysed variants, CAI ranged from 0.202 to 0.998 and tAI from 0.185 to 0.368.

Each synonymous construct received 15 unique barcodes, one from each intensity stratum, so synonymous variants could be distinguished despite encoding the same protein. Including 24 shared controls, the sublibrary comprised 2,524 constructs and 37,860 barcoded oligonucleotides. Expression, SEC fractionation and barcode quantification followed the pooled workflow. Missing barcode signals were set to zero and 10⁴ was added before arithmetic averaging and log₁₀ transformation. Protein amounts and yields were calculated using the calibration procedure below. The protein-level summary across synonymous variants was the median yield.

The original coding sequence for each parent was excluded from the variability analysis because CAI and tAI had not been assigned to these entries. Three additional variants were excluded because none of their barcodes met the representation threshold of 2 counts per million, leaving 2,447 variants across 50 proteins. CAI, tAI and log₁₀ yield were separately converted to percentile ranks within the pooled distribution of analysed variants. The interquartile range of each percentile-ranked quantity was then calculated within each parent. One-sided paired Wilcoxon signed-rank tests compared the CAI or tAI interquartile ranges with yield interquartile ranges (CAI: W = 1,275, P = 8.9 × 10⁻¹⁶; tAI: W = 1,258, P = 1.8 × 10⁻¹³). Median percentile-scale interquartile ranges were 0.480, 0.344 and 0.111 for CAI, tAI and yield, respectively. CAI and tAI distributions of coding sequences for de novo designs and DMS wild-type controls were summarized separately in Extended Fig. 4d.

### Pooled protein expression and affinity purification

Pooled libraries were expressed in E. cloni BL21(DE3). Each sublibrary glycerol stock was inoculated into 1 mL LB and incubated for 1 h at 37 °C, then transferred to 100 mL autoinduction medium and cultured overnight at 37 °C. Expression relied on autoinduction without IPTG addition or a temperature shift. Two 1-mL aliquots were pelleted at 6,000 × g for 5 min and stored at −30 °C for library-representation measurements. The remaining culture was harvested at 4,000 × g for 15 min in two approximately equal portions. One portion was frozen at −80 °C as a backup; the other, treated as approximately 50 mL culture equivalent, was processed for protein recovery.

Cell pellets were resuspended in 2.0 mL lysis buffer containing 50 mM Tris-HCl, pH 7.4, 200 mM NaCl, 20 mM imidazole, 1× cOmplete™, EDTA-free Protease Inhibitor Cocktail (Roche), 1× BugBuster (Millipore) and 125 U/mL Benzonase (Millipore). Samples were incubated with shaking for 15 min at room temperature and clarified at 6,000 × g for 20 min. Approximately 1.5 mL clarified lysate was incubated with 500 µL His Mag Sepharose Ni beads (Cytiva), prewashed three times with buffer A (50 mM Tris-HCl, pH 7.4, 200 mM NaCl and 20 mM imidazole), for 30 min at room temperature with rotation. Beads were washed three times with 1 mL buffer A. Bound proteins were eluted in 500 µL buffer B (buffer A with 400 mM imidazole) for 15 min at room temperature with rotation. Following elution, all protein samples were handled in low protein-binding tubes (SARSTEDT). An eluate aliquot was retained for SDS–PAGE, and the remaining sample was buffer-exchanged into 50 mM Tris-HCl, pH 8.0, and 150 mM NaCl using a Zeba spin desalting column (Thermo Fisher Scientific) and stored at 4 °C until SEC.

### Size exclusion chromatography and fraction collection

A 200-µL aliquot of buffer-exchanged sample was subjected to HPLC–SEC in 50 mM Tris-HCl, pH 8.0, and 150 mM NaCl at 0.4 mL/min. Eluate was collected in 0.5-mL fractions. Fractions around the expected monomer elution region were examined by SDS–PAGE and Coomassie staining alongside the affinity eluate or Agilent ProteoAnalyzer system (Agilent Technologies). The fraction assigned to the monomeric elution region was selected for barcode preparation. Monomeric status was assigned operationally from SEC elution behaviour; SDS–PAGE assessed the polypeptides present in each fraction. For nanobodies, the dimeric elution fraction was collected and processed separately by the same downstream workflow.

### Preparation of calibration and spike-in protein pools

Absolute quantification used a GFP-nanobody calibration pool and a separate 24-protein spike-in pool. Shared controls encoded in experimental sublibraries served a different purpose: they monitored agreement through expression and purification, whereas the externally added spike-in calibrated downstream barcode quantification.

The calibration pool consisted of a GFP-binding nanobody represented by 300 barcodes divided into five sets of 60. The sets were expressed separately in BL21(DE3), purified by His-tag affinity capture and quantified with a Qubit protein assay (Thermo Fisher Scientific). They were mixed at tenfold-spaced levels corresponding to 0.012–120 ng per barcode. Four calibration-input conditions (S1–S4) were each prepared twice. An ordinary least-squares fit to the five sets in S2 and S3 from both preparations (20 points) gave log₁₀(amount in ng per barcode) = 0.795 × log₁₀(MS intensity) − 5.342 (R² = 0.994; P = 1.6 × 10⁻²¹). Barcode-number performance was examined by repeated sampling without replacement, using 5–60 barcodes in increments of five and 50 subsamples per size.

The spike-in pool comprised 24 proteins, each represented by 15 barcodes, expressed and affinity-purified as a pool. Its total protein concentration was measured with a Qubit protein assay. Individual reference amounts were inferred from the calibration experiment. Barcode signals were aggregated, and outlying measurements among the eight S1–S4 preparation combinations were excluded using a modified median-absolute-deviation z-score threshold of 3.5. The remaining log₁₀ signals were averaged and converted using the calibration line. The resulting values specified ng per barcode for each protein in a 1-µg aliquot of the shared spike-in master mix.

A 1-µg aliquot of the spike-in pool was added to each monomeric or dimeric SEC sample immediately before proteolysis. The spike-in therefore calibrated peptide release, affinity capture, desalting and MS analysis, but did not directly correct losses during culture, lysis, affinity purification or SEC. The shared-control comparison was limited to sublibraries containing the control panel (Extended Fig. 1b).

### Peptide barcode preparation and desalting

For each monomeric or dimeric fraction, 400 µL sample containing approximately 40 µg protein and 1 µg spike-in standard was combined with an equal volume of Lys-C buffer (8 M urea and 100 mM Tris-HCl, pH 8.0). Endoproteinase Lys-C (New England Biolabs; 1 µg) was added and the mixture was incubated overnight at 37 °C. Protein concentrations were measured using a Qubit protein assay.

After Lys-C digestion, 25 µL Dynabeads HisTag Isolation and Pulldown beads (Thermo Fisher Scientific), prewashed twice with barcode buffer (50 mM Tris-HCl, pH 8.0, 150 mM NaCl and 0.1% Tween-20), was added without further dilution. Samples were incubated for 10 min at 25 °C with rotation. Beads were collected magnetically, washed twice with 200 µL barcode buffer and twice with 300 µL trypsin buffer (50 mM Tris-HCl, pH 8.0). Beads were resuspended in 50 µL trypsin buffer containing 0.25 µg Trypsin-ultra (New England Biolabs) and incubated overnight at 37 °C..

Beads were removed magnetically and the recovered supernatant was acidified with 2.5 µL 20% trifluoroacetic acid (TFA). Acidification was checked with pH indicator paper. GL-Tip SDB (GL Science) were conditioned with 20 µL acetonitrile and equilibrated with 20 µL 0.1% TFA/5% acetonitrile, centrifuging at 3,000 × g for 2 min after each step. Acidified samples were loaded at 3,000 × g for 3 min and washed with 200 µL equilibration solution. Peptides were eluted with 50 µL 0.1% TFA/50% acetonitrile at 3,000 × g for 2 min, dried in a centrifugal evaporator and stored at 4 °C until MS analysis.

### Mass spectrometry acquisition and barcode quantification

Peptides were analysed by nanoflow LC–MS/MS on an Orbitrap Astral mass spectrometer (Thermo Fisher Scientific; instrument software 2.1.132.17) at Nagoya University, except for the data shown in Fig. 1b, which were acquired using a Q Exactive mass spectrometer (Thermo Fisher Scientific). Dried peptides were reconstituted in 30 or 80 µL of 0.1% trifluoroacetic acid containing 2% acetonitrile, and a 3-µL aliquot was injected for LC–MS/MS analysis. Peptides were analyzed using an Orbitrap Astral mass spectrometer coupled to a Vanquish Neo nanoLC system (Thermo Fisher Scientific) equipped with a nano-electrospray ionization source. A trap-and-elute configuration was used. Most samples were separated on an NTCC-360/75-1.9-123 C18 reversed-phase column (75 µm i.d. × 120 mm, 1.9 µm particle size; Nikkyo Technos Co., Ltd.); a 250-mm C18 column was used for a limited subset of measurements. The column temperature was maintained at 50 °C using an external column heater. Mobile phases A and B consisted of 0.1% formic acid in water and 0.1% formic acid in acetonitrile, respectively.

For the principal 120-mm-column method, the gradient was programmed as follows: 4–12% B over 0–3.5 min, 12–13% B over 3.5–4 min, 13–30% B over 4–18 min, and 30–36% B over 18–20 min. Solvent B was then increased to 76% at 20.5 min and maintained until 25 min, followed by column re-equilibration. The flow rate was 600 nL/min through 3.5 min, decreased to 300 nL/min at 4 min, and maintained at 300 nL/min thereafter.

Data were acquired in positive-ion data-independent acquisition (DIA) mode, with a spray voltage of 2.0 kV and an ion transfer tube temperature of 290 °C. Full MS spectra were acquired in the Orbitrap analyzer over m/z 380–980 at a resolution setting of 240,000, with an automatic gain control (AGC) target of 5 × 10⁶, and recorded in profile mode. DIA MS/MS spectra were acquired in the Astral analyzer over m/z 150–2,000 using 300 nominally 2-Th isolation windows spanning the precursor range of m/z 380–980, with window placement optimization enabled and no window overlap. Higher-energy collisional dissociation was performed at a normalized collision energy of 25. The MS2 AGC target was 5 × 10⁴, and MS/MS spectra were recorded in centroid mode. The MS1 and MS2 injection-time settings were 5 and 3 ms, respectively. The total MS acquisition time was 27 min.

Thermo raw files were converted to mzML using ThermoRawFileParser 1.4.5 and analysed with DIA-NN 2.2.0^135^. A predicted spectral library was generated from a FASTA containing 45,952 barcode peptides. Parameters included cleavage after K/R, zero missed cleavages, peptide lengths of 7–20 residues, precursor m/z 375–985, fragment m/z 150–2,000 and charge 2+. Methionine oxidation was allowed, with at most two variable modifications per precursor; no fixed modification was applied. Searches used 10-ppm MS1 and MS2 mass tolerances and a 1% precursor false-discovery-rate threshold. Quantification used the precursor-level matrix, keyed by stripped peptide sequence. Barcode assignments were unique within each experimental sublibrary, and results were joined to the corresponding sublibrary-specific map.

### Calibration and calculation of monomeric yield

Barcode intensities were processed separately for each sublibrary and SEC fraction. Missing signals were set to zero and a pseudocount of 10⁴ was added. Before aggregation, individual barcodes with plasmid-library representation <2 counts per million (CPM), including barcodes with no matching read, were excluded (see “DNA synthesis and library cloning” for read processing and CPM calculation). For each construct, the arithmetic mean across the remaining barcodes was calculated and then log₁₀-transformed. Constructs were excluded entirely if none of their 15 barcodes passed the filter. The log₁₀(intensity + 1) transform used in the mGreenLantern barcode screen was specific to barcode selection and was not used for yield estimation.

For sublibraries containing shared controls, a between-sublibrary intensity correction was applied before construct-level aggregation. The ratio of median spike-in intensity to median control intensity was divided by a common monomer-channel target ratio of 0.6 to obtain a scale factor. Experimental-construct intensities were multiplied by this factor; spike-in and control intensities were not rescaled. The target ratio was selected empirically from the observed sublibrary ratios.

For each sublibrary and fraction, the aggregated log₁₀ signals of the 24 spike-in proteins were related to their reference log₁₀ amounts in ng per barcode by ordinary least squares. No weighting or spike-in exclusion was applied at this step. The fitted relationship, log₁₀(amount per barcode) = slope × log₁₀(MS signal) + intercept, was then applied to the experimental constructs. Monomeric and dimeric samples each received 1 µg of the spike-in pool.

The inferred amount per barcode was converted to an individual-culture-equivalent yield using the following scaling:

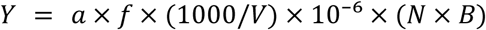

Here, Y is yield in mg/L, a is the inferred amount in ng per barcode, f is the scale-up factor from the analysed sample to the nominal preparation, V is the processed culture volume in mL, N is the number of constructs represented in the pooled culture and B is the number of barcodes per construct. The monomeric and dimeric analyses used f = 2.5, V = 50 and B = 15. The factor 2.5 corresponds to scaling a 200-µL SEC injection to a nominal 500-µL affinity eluate. Multiplication by N × B assumes equal representation of barcoded constructs when expressing a pooled measurement as an individual-culture equivalent. Yields were reported as log₁₀(mg/L).

For analyses of the main de novo and mutant collections, log₁₀ yield values below −1.5 were set to −1.5, corresponding to approximately 0.032 mg/L. Values at this floor represent measurements below the practical quantification range. Unclipped yield values were used for visualization in Extended Fig. 4c. The nanobody sublibrary did not include the shared control panel used for the other sublibraries, so the same lower quantification threshold could not be assumed to apply. Nanobody monomeric and dimeric yields were therefore not clipped at −1.5, and the log₁₀ monomer/dimer yield ratio was calculated as the difference between the unclipped log₁₀ yields.

### Individual expression and HPLC-based yield measurement

Forty-eight proteins spanning the pooled-assay yield range were selected for individual expression. Each construct retained a peptide barcode assigned in the pooled library. Constructs were cloned into pTF3 with a C-terminal His₆ tag using NdeI and XhoI, propagated in DH5α, and verified by colony PCR and Sanger sequencing.

Verified plasmids were transformed into E. cloni BL21(DE3). Single colonies were expanded in LB–kanamycin and used to inoculate autoinduction cultures, which were grown overnight at 37 °C. For the first nine-protein batch, 50 mL of a 100-mL culture was processed. Later batches and the prospective binder set were expressed and harvested at 40 mL per construct. Yields were normalized to the processed culture volume.

Cells were harvested at 4,000 × g, stored at −80 °C, lysed and purified with His Mag Sepharose Ni beads, then buffer-exchanged into SEC buffer with Zeba spin desalting columns. The nominal final eluate volume was 500 µL. Each construct was evaluated by one culture, one purification and one HPLC–SEC injection.

Individual proteins were analysed by HPLC–SEC with a 200-µL injection and 0.4-mL/min flow in 50 mM Tris-HCl, pH 8.0, and 150 mM NaCl, without fraction collection. Absorbance was recorded at 215 and 280 nm using Shimadzu LabSolutions; the 215-nm trace was used for quantification.

Chromatograms were baseline-corrected with a rolling third-percentile baseline over 10,001 points. Monomer peak area was integrated by the trapezoidal rule over a batch-specific window of approximately 30–40 min. An external cytochrome c calibration related peak area to injected mass on a log–log scale, with slope 0.9730 and intercept −7.292. The standard was measured as a twofold dilution series with 100-µL injections in duplicate. Inferred injected mass was multiplied by 2.5 to scale to the nominal 500-µL eluate and divided by processed culture volume to obtain mg/L.

The 21 prospective BindCraft designs were expressed and processed with the later individual-expression batch using the same SEC method, calibration and 40-mL culture normalization. These constructs carried a C-terminal His₆ tag but no peptide barcode. Candidate selection and group definitions are described under prospective validation.

### Folding stability and mutational effect analysis

Folding stability was measured in a dedicated cDNA-display proteolysis experiment using the assay framework of Tsuboyama et al. (2023)^116,136^. The lib82_6 collection contained 97,671 clone entries: 38,830 main entries, 5,984 entries overlapping the previously published dataset and 52,857 scrambled controls. Stability estimates were obtained for 97,347 entries. Protein–cDNA fusions were challenged with trypsin or chymotrypsin across serially diluted protease concentrations, and surviving constructs were quantified by NGS. The published Bayesian K₅₀/ΔG inference pipeline was used to infer protease resistance and ΔG in kcal/mol; more positive values indicate greater stability. Inferred values outside the fitted range were bounded at ±15 kcal/mol. Overlapping sequences were compared with published measurements as a quality check; the stability–yield comparison used measurements from the new experiment.

Stability and yield were matched by clone identifier within the main collection. Mutational effects were computed as mutant minus the corresponding wild-type reference for both ΔG and log₁₀ yield. Wild-type reference values were averaged across repeated wild-type constructs for each parent. Yield values were floored at −1.5 before differencing. Substitutions were assigned source and target amino acids from construct annotations; deletions and alanine insertions were analysed as separate classes. Mean substitution effects by source or target residue were unweighted arithmetic means across eligible variants.

For the PROSS analysis (Extended Fig. 12e), increases and decreases relative to the corresponding wild type were classified using thresholds of a twofold change in monomeric yield and a 1 kcal/mol change in folding stability (ΔG).

Protein-specific relationships were summarized by ordinary least-squares regression of log₁₀ yield on ΔG. Protein-level slopes were compared with sequence hydrophobicity and structural descriptors of the relaxed wild-type model. The hydrophobicity scale was the score_monomeric_designs scale: F 100, I 99, W 97, L 97, V 76, M 74, Y 63, C 49, A 41, T 13, H 8, G 0, S −5, Q −10, R −14, K −23, N −28, E −31, P −46 and D −55. Structural descriptors included buried nonpolar area, buried nonpolar residue counts and hydrophobic residue counts. Relative solvent accessibility used the normalization of Tien et al. (2013)^137^, with core, boundary and surface positions defined by values <0.2, 0.2–<0.4 and ≥0.4, respectively.

### Structure prediction and structural descriptors

AlphaFold2 predictions were generated with local ColabFold^138^ in single-sequence mode, without templates, using one model and three recycles. The sole rank-1 model was used. The principal AlphaFold3 pLDDT/TM-score benchmark used the Kuhlman-Lab single-sequence-capable fork (commit 87bd9e678d9acacc4aa9baa05e820f32b80e1b49), without the data pipeline or template search, with seed 1, three recycles and one diffusion sample. Separate AlphaFold3 runs supplied structures for ESM3ΔG, using ten recycles and five diffusion samples; the top AlphaFold3-ranked model was used for these stability predictions.

Nanobody monomers were predicted as one chain, target complexes as nanobody and antigen chains, and homodimers as two identical nanobody chains. AlphaFold3 nanobody predictions used one seed and empty MSA/template inputs. ESMFold2 predictions used biohub/ESMFold2. The DMS runs used revision 1ebf0e3481a5184eb6171d40615c79e384b48796 and nanobody runs used 8fc3ff471022fdce52c77030685eb775de0c00a3. DMS monomers used one recycling loop and 16 sampling steps; DMS homodimers and nanobody monomer/target-complex runs used three loops and 50 steps. ESMFold2 predictions used one diffusion sample and seed 0. Designed chains were supplied without an MSA; the natural antigen chain in nanobody–target ESMFold2 runs used a precomputed MSA.

Structural agreement with the design backbone was quantified by TM-score^139^ using tmtools on Cα coordinates, normalized by the reference-backbone length. AlphaFold3 pLDDT was summarized as the mean per-atom confidence; AlphaFold2 confidence was obtained from ColabFold score files. The main filter required AlphaFold3 pLDDT >90 and TM-score >0.8. Other pLDDT and TM-score thresholds were evaluated for the filtering analyses in Fig. 2 and Extended Fig. 5.

Backbone descriptors were calculated from design-backbone PDB files. Radius of gyration was the root-mean-square Cα distance from the centroid. Relative contact order was the sum of sequence separations over Cα pairs separated by <8 Å, divided by chain length and the number of contacts. For the core-residue fraction, Cα neighbours within 14 Å were counted after excluding pairs with sequence separation ≤10; a residue was classified as core if it had >15 neighbours. Helix, sheet and loop fractions were obtained from the Rosetta feature pipeline’s secondary-structure assignment.

Structures were relaxed using Rosetta with beta_nov16, constraints to the starting coordinates and constraint ramping disabled. Rosetta version 2020.50.post.dev+7.master.5a82c7c was used with the score_monomeric_designs feature pipeline (commit 134d3597c73ed83a6c6430bf786215d6350b0a6a), following the framework of Rocklin et al. (2017)^42^. Features included energetic terms, buried and exposed surface areas, hydrophobicity, packing, core/boundary/surface composition, secondary structure, contacts, fragment quality and sequence composition. The same feature-extraction framework was applied to the main protein and nanobody collections.

Predicted folding stability was obtained with ESM3ΔG^117^, using the mean prediction of three LoRA-augmented checkpoints. Predictions were reported in kcal/mol. The main de novo and DMS inputs were the separately generated AlphaFold3 structures described above, rather than the structures from the principal pLDDT/TM-score filtering run.

### Dataset partitioning for the general prediction models

Data were split for the general-model evaluation using MMseqs2 sequence clusters together with backbone identity. Sequence clustering was performed using MMseqs2 easy-cluster (build 23503f3bbd4ea19bfe03beaba3b8def27d5f8187) with a minimum sequence identity of 0.3 (30%), coverage 0.8, bidirectional coverage mode 0 and set-cover clustering mode 0; other parameters were left at their defaults. The sequence input and cluster assignments are recorded in the final split manifest.

De novo designs are grouped by backbone identity. Backbones connected through shared MMseqs2 clusters are linked into indivisible groups before fold assignment. Groups are distributed across five folds with a deterministic size-balancing procedure: groups are shuffled with seed 42, ordered by descending sequence count and assigned to the currently smallest fold.

For both LoRA and XGBoost cross-validation, the same fold assignments were used: one fold was held out for testing, the next fold in cyclic order was used for validation and the remaining three folds were used for training. Test and validation roles rotate across the five runs. Duplicate amino acid sequences and synonymous constructs must retain a consistent group assignment, and measurement aggregation is performed before evaluation.

Mutant–wild-type pairs are assigned with reference to the mutant cluster, wild-type cluster and parental backbone. Pairs related to a test or validation group are excluded from that fold’s training set. General-model performance was assessed using held-out predictions from this partitioning.

The split manifest records sequence and backbone assignments, mutant–wild-type pair assignments, per-fold counts and checks for overlap among training, validation and test sets. This partitioning was used for the XGBoost and LoRA model evaluations.

### pLM embeddings and XGBoost regression

Frozen mean-pooled embeddings were extracted from ESMC 6B, ESM2 3B and SaProt 1.3B. ESMC used biohub/ESMC-6B in bfloat16 with batch size 1; the 2,560-dimensional last hidden state was averaged over attention-mask positions, excluding padding but including tokenizer special tokens. ESM2 used esm2_t36_3B_UR50D with residue-mean layer-36 embeddings (2,560 dimensions), excluding BOS/EOS. SaProt used westlake-repl/SaProt_1.3B_AFDB_OMG_NCBI in float32, with 1,280-dimensional residue-mean embeddings excluding special tokens. For frozen SaProt embeddings, structural tokens were set to the unknown symbol #. Structure-aware SaProt inputs were used separately for LoRA fine-tuning, with amino acid and Foldseek 3Di symbols interleaved per residue. Mutants and wild types used their own predicted structures; insertions and deletions were represented at their respective sequence lengths.

XGBoost regressors used explicit numeric sequence and structural descriptors, frozen embeddings, or combinations of these with predicted ΔG. The feature table contained 1,025 numeric descriptors, including Rosetta-derived features and six AlphaFold3 confidence metrics. Eleven input combinations were considered: descriptors alone; each of three embedding sets alone; each embedding set with descriptors; and descriptors or each embedding set with predicted ΔG. Missing features were handled by XGBoost, and records without yield labels were excluded.

The XGBoost target was log₁₀ monomeric yield, bounded to −1.5 to 3.5 for model training. A custom objective combined squared error on absolute yields of de novo and wild-type proteins with squared error on mutant-minus-wild-type changes, using weights 1.0 and 0.5. Mutant absolute yields were not separate targets. The settings were max_depth = 3, n_estimators = 500, learning_rate = 0.03, subsample = 0.8, colsample_bytree = 0.8 and random_state = 42, without validation-based early stopping. Predicted mutational effects were computed as predicted mutant yield minus predicted wild-type yield.

### Fine-tuning with low-rank adaptation

LoRA fine-tuning was applied to ESMC 6B, ESMC 600M, ESM3-open 1.4B and SaProt 1.3B. Checkpoints were biohub/ESMC-6B, biohub/ESMC-600M, esm3_sm_open_v1 and westlake-repl/SaProt_1.3B_AFDB_OMG_NCBI. ESMC used amino acid sequences. ESM3 encoded predicted structures with ESMProtein.from_pdb, and SaProt used interleaved amino acid/3Di tokens. Mutant and wild-type structural inputs were generated separately.

LoRA adapters were added with PEFT at rank 4 and alpha 8, with frozen pretrained weights. ESMC used dropout 0.05 and out_proj adapters. ESM3 used dropout 0.15 with attn.layernorm_qkv.1 and attn.out_proj adapters. SaProt used dropout 0.15 with query, key, value and attention.output.dense adapters. For ESMC 600M, only a linear regression head was evaluated. For ESMC 6B, ESM3-open 1.4B and SaProt 1.3B, both linear and multilayer perceptron (MLP) regression heads were evaluated. These heads were applied to the residue representations. Per-residue outputs were averaged over non-special-token positions. The tokenizer maximum length was 512.

The LoRA objective combined a Huber loss (δ = 1.0) for absolute log₁₀ yields of de novo designs and unique wild types with a mean-squared-error loss for mutant-minus-wild-type differences, weighted 1.0 and 0.5, respectively. Yield labels were bounded to −1.5 to 3.5 before computing targets. Independent loaders supplied absolute-yield examples and mutant–wild-type pairs at a 1:1 batch ratio; the shorter loader was cycled and wild-type self-pairs were excluded.

Training used AdamW with learning rates of 5 × 10⁻⁵ for LoRA parameters and 10⁻⁴ for the regression head, weight decay 0.01 and gradient-norm clipping at 1.0, without a learning-rate schedule. ESMC and SaProt used micro-batches of two with eight accumulation steps; ESM3 used micro-batches of one with 16 accumulation steps. The epoch limits were ten for ESMC/SaProt and twenty for ESM3/SaProt. Early stopping used validation-set de novo Spearman correlation with patience five, retaining the best-validation checkpoint. Python, NumPy and PyTorch seeds were set to 42. Training used a single RTX PRO 6000 Blackwell GPU per model run. ESMC 6B LoRA was designated ESMC-MonoYield.

### Comparison with existing predictors and prospective validation

Comparators included CamSol, AGGRESCAN3D, GATSol, NetSolP, SKADE, MPEPE, SoluProt, RP3Net and DeepSoluE. CamSol used the intrinsic-solubility score; NetSolP-1.0 used the distilled solubility model; SoluProt used version 1.0.1.0; RP3Net used rp3net_v0.1_d.ckpt; and MPEPE used MODELs-1027 with linker-trimmed coding sequences. AGGRESCAN3D was run through de-STRESS and GATSol through its ESM-1b/contact-map pipeline. Remaining sequence predictors used their reported sequence inputs. Scores were oriented so higher values indicated favourable predictions. Uncalibrated comparator scores were assessed by correlation and ranking; their raw-score RMSE or R² cannot be interpreted as calibrated yield error.

Ranking performance was measured as the percentage of true positives among the highest-scoring fraction of candidates. High-yield de novo designs were defined as having monomeric yield >10 mg/L. For the mutational-selection analysis, positives were defined as at least a fivefold increase in monomeric yield over wild type, equivalent to Δlog₁₀ yield ≥ log₁₀(5) ≈ 0.699. Selection fractions and baseline positive rates were calculated for the evaluation population shown in each panel. Error bars for the positive rates in Fig. 4f and Extended Fig. 12d (left) represent pointwise 95% Wilson confidence intervals, calculated separately for each predictor and selection fraction using the number of positives and the total number of selected candidates. For the empirical cumulative distribution function (ECDF) analysis (Extended Fig. 12a,d), candidates were ranked separately within each evaluation population (all data, de novo designs and mutational effects) using each predictor, and the top 1%, 5%, 10%, 15% and 20% were selected. ECDFs were calculated from observed log₁₀ monomeric yields for the all-data and de novo sets and from observed mutant-minus-wild-type changes in log₁₀ yield for the mutational-effect set.

Binders against the SARS-CoV-2 spike receptor-binding domain (RBD) were designed de novo using BindCraft v1.2.0 (commit 1163713; ColabDesign 1.1.2, JAX 0.4.27 and PyRosetta 2024.42). The target comprised residues 334–527 of chain B from PDB 7JZM, with the bound LCB3 minibinder removed. Residues 337, 340, 343, 345 and 356 were specified as hotspots, and binder lengths ranged from 70 to 90 residues. Design used the four-stage AlphaFold2 multimer protocol with 75 soft, 45 temporary, 5 hard and 15 greedy iterations, a helicity weight of−0.3, and one recycle during design and three during validation. Each hallucinated backbone was redesigned using ProteinMPNN with 20 sequences per trajectory at a sampling temperature of 0.1, with interface residues fixed and cysteine excluded. The two highest-ranked sequences per trajectory were re-predicted using AlphaFold2 and evaluated using the default BindCraft filters.

For prospective validation, a candidate pool of 550 designs passing the default filters was assembled. Applying an additional Average_i_pTM threshold of >0.8 yielded 226 eligible candidates. These candidates were ranked by the mean predicted log10 monomeric yield from the five ESMC-MonoYield cross-validation models. Eleven high-scoring candidates and ten candidates randomly sampled from the same eligible pool were selected for experimental validation. All constructs were expressed and purified in a single batch and analysed once each using the individual HPLC–SEC procedure described above. Peaks were integrated over 33–40 min, and yields were normalized to a 40-mL culture volume. Group yields were compared using a one-sided Mann–Whitney U test. The fractions exceeding 10 mg/L were reported descriptively.

### Nanobody sequence rules and task-specific prediction

Nanobody analyses used the fixed-framework library with invariant CDR1, six-residue CDR2 and five-to thirteen-residue CDR3. Monomeric and dimeric log₁₀ yields were not clipped, and their difference was used to calculate the log₁₀ monomer/dimer ratio. CDR2 positions and N- or C-terminally aligned CDR3 positions were analysed by residue identity. Heatmap cells with ten or fewer designs were masked.

Local structural analyses focused on invariant F40 in FR2 and C99–A100–A101 in FR3. Rosetta descriptors included energies at sites 40 and 101, mean energies within 3 or 5 Å of site 100, and energies of local two- and four-residue segments around site 100. Representative relaxed models were inspected for CDR3 packing relative to these framework sites (Fig. 5e–g).

Associations were screened across 2,496 features, comprising 2,239 structural/energetic features and 257 CDR sequence features, against eight yield-derived response variables. Pearson correlations were calculated in the pooled dataset and by design method. Benjamini–Hochberg correction was applied separately within each response and method subset; features were selected at q <0.01 and |r| ≥0.3. Features correlated with one another at |r| ≥0.9 were represented by one member for display.

The six-rule scheme adapted from McMahon et al. (2018)^111^ required CDR2 position 1 to be A, D, G, N, S or T; position 4 to be G; position 5 to be A, G, S or T; and position 6 to be I, N, S or T. CDR3 C1 was required to be Y and C3 to be F, H, L or Y. Absence of cysteine and methionine was recorded but did not contribute to the score. The expanded scheme retained the four CDR2 rules and the C3 rule, broadened C1 to Y or W, and added CDR3 N1 not in Y/W/V/F/L/I, N2 in L/T and C2 not in F/L/Y. Scores ranged from zero to six or zero to nine. The expanded preferences were derived from the current dataset and were evaluated descriptively in 2,590 designs with complete monomeric and dimeric measurements.

ESMC-Nb-MonoYield and ESMC-Nb-MDRatio were independently initialized from pretrained biohub/ESMC-6B, with newly initialized LoRA adapters and regression heads. They were not initialized from ESMC-MonoYield. Each used rank-4 LoRA, alpha 8, dropout 0.05 and out_proj adapters, with a residue-wise linear head and maximum token length 512. AdamW learning rates were 5 × 10⁻⁵ for LoRA and 10⁻⁴ for the head, with weight decay 0.01 and gradient clipping at 1.0. Training used micro-batches of two with eight accumulation steps, Huber loss with δ = 1.0, up to ten epochs and early stopping based on validation Spearman correlation with patience five. The seed was 42. Nanobody monomeric-yield and monomer/dimer-ratio labels were used without clipping.

Nanobody cross-validation was defined independently of the general-model split. CDR2 and CDR3 sequences were concatenated, and pairwise Levenshtein distances were divided by the longer sequence length. Complete-linkage clustering at normalized distance 0.70 generated 105 clusters across the 2,600 designs. Clusters were assigned to five groups by size-balancing greedy assignment with seed 42. One group was used for testing, the next for validation and the other three for training, rotating cyclically. Each cluster was assigned wholly to one group, and performance was calculated from one out-of-fold prediction per evaluable design.

OmniLib CNN and linear-regression models used cnn_24_fc_8.pt and 20231223_LR.pt from omnilib-ml (commit 2f0765af2342e06c7b7bd7023975b37bda04a4f0; Wan et al., 2026)^128^. CDRs were annotated with AntPack 0.3.8.6 in IMGT numbering. Of 2,600 ordered designs, 2,590 had complete pooled yield measurements; one lacked Rosetta features, leaving 2,589 in the feature/model table. Two hundred designs were outside the OmniLib length ranges, leaving 2,389 scorable designs. Ranking was evaluated by top-fraction precision. Positive thresholds were monomeric yield ≥3.16 mg/L (log₁₀ yield ≥0.5) and monomer/dimer ratio ≥10 (log₁₀ ratio ≥1).

### Nanobody binding enrichment assay

Nanobody binding enrichment was assessed with a separate cDNA-display selection library containing the 2,600 nanobody designs. This library was distinct from the proteolysis library and was analysed by changes in sequence abundance after target capture, without protease challenge.

Templates were PCR-amplified from the nanobody binding-library plasmids using KK32 and Setsu714, with qPCR-selected cycle numbers, and gel-purified. RNA was transcribed overnight at 37 °C using T7-Scribe, DNase I-treated and column-purified. Approximately 2 pmol RNA was annealed to 2 pmol cnvK–puromycin linker in 10 µL containing 200 mM NaCl and 40 mM Tris-HCl, pH 7.4, by cooling from 95 to 45 °C at 0.1 °C/s and then to 25 °C. Complexes were crosslinked at 365 nm for approximately 15 min, translated using PUREfrex 2.0 (Gene Frontier) in 40 µL for 2 h at 37 °C, and treated with EDTA. Complexes were captured on Dynabeads MyOne Streptavidin C1 beads (Thermo Fisher Scientific), reverse-transcribed on beads for 30 min at 37 °C, released with RNase T1 for 15 min at 37 °C, purified on His-tag magnetic beads, eluted with 400 mM imidazole, buffer-exchanged into PBS and stored at −80 °C.

Selections used biotinylated PD-1 or PD-L1 in blocking buffer (PBS, 0.5% BSA, 0.05% Tween-20 and 100 µg/mL tRNA). PD-L1 was Acro Biosystems PDL-H82E4, residues 19–134. The protocol included target concentrations of 500 and 100 nM in round 1 and 100 and 20 nM in round 2, with no-target controls. Binding reactions were captured with 20 µL blocked streptavidin M-280 Dynabeads in a final volume of 70 µL for 5 min. Beads were washed twice with blocking buffer and twice with PBS/0.05% Tween-20. Captured material was released with proteinase K at 55 °C for 15 min, followed by 95 °C for 10 min. Input samples were processed with the same proteinase K treatment. Recovered cDNA was re-amplified for the next round. One selection was performed per target per round.

Input and first- and second-round samples for each target were converted to Illumina libraries by two-step PCR, purified with Sera-Mag beads and sequenced on a NextSeq. Read 2 was trimmed with cutadapt 5.1, reverse-complemented and matched to a 109-nucleotide reference window using seqkit 2.10.0 and custom analysis. Counts were normalized by sample total, and enrichment was calculated as selected normalized abundance divided by input normalized abundance, displayed as log₂(enrichment + 0.1). Designs undetected in the input were excluded from enrichment plots and correlations. The analysis was joined to the yield table by construct identifier.

### Statistical analysis

Associations were summarized with Pearson correlation and, where indicated, Spearman rank correlation using SciPy. Linear fits used ordinary least squares. Yield-prediction RMSE was calculated on the log₁₀ scale, and R² was calculated with scikit-learn. Sample sizes refer to evaluable constructs, variants or parental backbones as specified for each analysis; pooled-assay constructs are not biological replicates of one another.

Correlation uncertainty was estimated by nonparametric bootstrap. After excluding non-finite pairs, 1,000 resamples of n observations were drawn with replacement from the n evaluable rows (seed 0). Resamples with zero variance in either variable were discarded. Error bars show the 2.5th and 97.5th percentiles of bootstrap correlations. Resampling units followed the analysed table: constructs for sequence-level analyses and parental proteins or backbones for protein-level analyses.

Barcode-number analyses sampled without replacement within the corresponding protein or calibration set. These procedures are subsampling analyses rather than nonparametric bootstraps. The calibration-barcode analysis used 50 subsamples per barcode count; the independent-assignment analysis used repeated subsamples of the two 40-barcode libraries (Extended Fig. 1a).

Model metrics were computed on pooled held-out predictions, rather than as the mean of per-fold metrics. Nanobody-specific evaluation used one out-of-fold prediction per evaluable design. The final general-model evaluation uses the revised split manifest and its specified prediction-aggregation procedure. Ranking performance was summarized as top-fraction precision, with baseline positive rates calculated from the same evaluation population.

For the nanobody feature screen, correlation P values were adjusted by the Benjamini–Hochberg procedure within each response variable and design-method subset, using q <0.01 and |r| ≥0.3 as selection criteria. Other displayed correlations were descriptive and were reported with unadjusted P values or bootstrap confidence intervals, as indicated.

Within-protein percentile-scale variability in CAI or tAI was compared with yield variability using one-sided paired Wilcoxon signed-rank tests. The prospective binder groups were compared using a one-sided Mann–Whitney U test with the model-selected group specified as the greater-yield alternative. Nanobody two-group comparisons, where reported, used two-sided Mann–Whitney U tests. The test, direction, sample size and response variable are specified in the corresponding analyses.

Missing values were handled by pairwise deletion in correlations and regressions. Designs lacking a required yield or structural descriptor were excluded from the corresponding analysis. Input-undetected clones were excluded from binding-enrichment plots. The separate treatment of missing MS signals, barcode-level NGS filtering and yield flooring is described above.

Box plots show medians and interquartile ranges. Unless otherwise indicated, whiskers extend to the most extreme values within 1.5 interquartile ranges. The per-amino-acid mutational-effect panels show quartile boxes without extended whiskers or outlier markers. When all observations are shown as overlaid points, separate outlier markers are omitted. Regression lines are shown without confidence bands unless stated otherwise.

Analyses used Python with NumPy, pandas, SciPy, scikit-learn, statsmodels, Matplotlib, seaborn and Biopython. XGBoost and pLM training used separate environments, including PyTorch, Hugging Face Transformers and PEFT. Software versions and model revisions are specified with the corresponding analysis procedures.

### Use of generative AI

OpenAI’s ChatGPT and Anthropic’s Claude were used to assist with manuscript preparation, including language editing, improving clarity and organization, and refining scientific writing. All AI-assisted content was critically reviewed and verified by the authors, who take full responsibility for the accuracy, integrity, and conclusions of the manuscript.

## Data Availability

### Data availability

Source data supporting all figures are provided in Supplementary Tables S1–S25 and the associated Zenodo repository [DOI: 10.5281/zenodo.23874456]. The Index sheet of the supplementary Excel workbook maps each figure panel to its source data. Raw and intermediate data, including sequencing read counts, mass-spectrometry data, HPLC chromatograms, structural features and predicted structures, are available in the Zenodo repository under a CC BY 4.0 license.

### Code availability

Code for data analysis, model training and monomeric yield prediction is available at https://github.com/tsubo-lab/monoyield-predictor/ (Apache-2.0. Trained model weights are available at https://huggingface.co/Tsuboyama-Lab/monoyield-predictor.

### Supplementary Data

Supplementary_Tables.xlsx. Supplementary Tables S01–S25 provide the source data underlying the figures in this study. The workbook contains one worksheet per table and an Index sheet linking each table to the corresponding figure panels. Tables S05, S23 and S24 exceed spreadsheet size limits and are available as CSV files in the associated Zenodo dataset. Together, these tables include barcode- and protein-level yield measurements from pooled peptide-barcode mass spectrometry for the main and nanobody libraries; sublibrary calibration constants; data from spike-in, control and HPLC validation experiments; model predictions presented in Figs. 4 and 5; and design sequences and structural metrics. Each measurement is recorded once within its library. Rows are identified by the combination of sublibrary (sublib) and construct name (name), with peptide barcode sequences providing an additional identifier in barcode-level tables.

## Acknowledgments

We thank members of the Tomari and Tsuboyama laboratories for critical comments on the manuscript and Gabriel J. Rocklin at Northwestern University for valuable advice and discussions. We also thank Mamoru Ishii in the Tsuboyama laboratory for sharing SARS-CoV-2 binder designs, and Keiko Kano and Shinya Sato at Nagoya University for performing mass spectrometry analysis.

This study was supported by PRESTO (JPMJPR21E9 to K.T.), FOREST (JPMJFR230Z to K.T.), CREST (JPMJCR23B6 to K.T.), A-STEP (JPMJTR24U7 to K.T.), BOOST (JPMJBY24E9 to K.T.) and GteX (JPMJGX23B9 and JPMJGX23B4 to K.T.) from the Japan Science and Technology Agency (JST); the UTOPIA Young Researcher Development Program, UTOPIA AI Research Discovery Program and UTOPIA Next-Generation Diagnostic Tool Discovery Program (JP223fa627001 to K.T.) from the Japan Agency for Medical Research and Development (AMED); Grants-in-Aid for Transformative Research Areas (A) (24H01117, 24H01356, 26H01795 and 26H00913 to K.T.), Scientific Research (B) (25K00103 to K.T.), Challenging Research (Exploratory) (26K23134 to K.T.) and JSPS Research Fellows (RPD) (26KJ0105 to S.S.) from the Japan Society for the Promotion of Science (JSPS); and grants from the Uehara Memorial Foundation (to K.T.), the Nakatani Foundation (to K.T.), Leading Pioneers Science Foundation (to K.T.), SECOM Science and Technology Foundation (to K.T.), the Takeda Science Foundation through its Visionary Research Grants (Start) and Visionary Research Continuous Grants (Hop) (to K.T.), the Inoue Foundation for Science (to K.T.), the Nakajima Foundation (to K.T.), the Daiichi Sankyo Foundation of Life Science (to K.T.), the Mitsubishi Foundation (to K.T.) and the Canon Foundation (to K.T.).

## Contributions

S.S. and K.K. contributed equally to this work. K.T. conceived the study. S.S. analysed the data, and K.K. performed most of the experiments. A.Y.W.L. performed the cDNA display proteolysis experiments. E.M.-S. performed and oversaw the mass spectrometry analysis at Nagoya University. Y.T. and K.T. supervised the research. S.S. and K.T. wrote the manuscript. All authors have read and approved the final manuscript.

## Extended figure legends

**Extended Figure 1.**
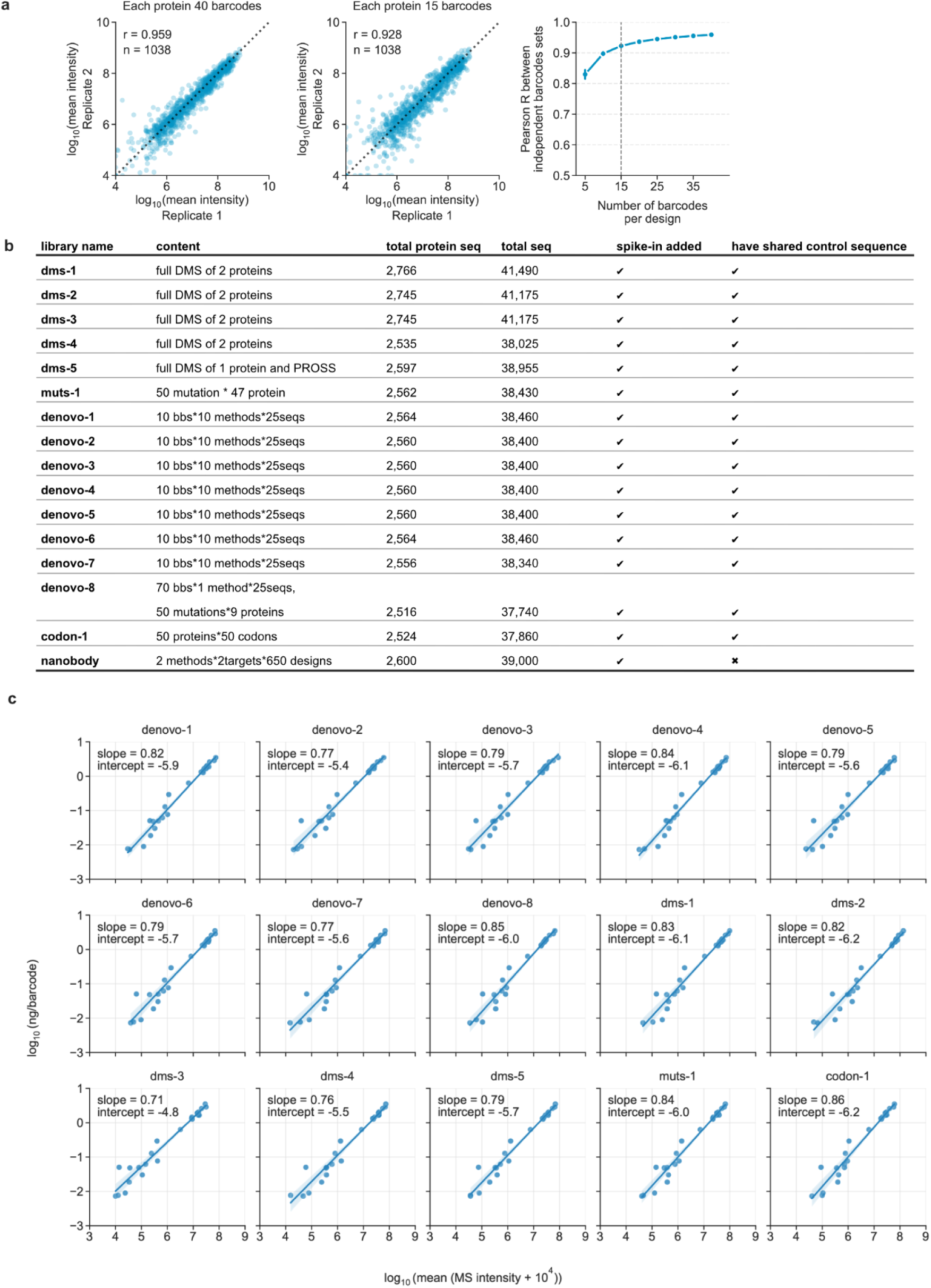
Barcode-assignment reproducibility, sublibrary composition and spike-in calibration. **a**, Comparison of independently assigned barcode sets for the same 1,038 proteins. Left, protein-level log₁₀ mean intensities calculated using all 40 barcodes per protein (Pearson’s r = 0.959). Middle, a representative comparison using 15 barcodes per protein (r = 0.928). Right, Pearson correlations between barcode assignments after subsampling 5–40 barcodes per protein, with 25 subsampling iterations at each barcode count. Error bars represent SD, and the vertical dashed line marks 15 barcodes per protein. Missing signal intensities were set to zero, and an offset of 10⁴ was added to all values before averaging. **b,** Composition of each sublibrary, including the numbers of protein constructs and barcoded entries. Separate columns indicate whether an external spike-in standard was added and whether shared control proteins were encoded within the sublibrary. The nanobody library received the spike-in standard but did not contain the shared control proteins. **c,** Sublibrary-specific calibration relationships between log₁₀ MS intensity (horizontal axis) and the reference log₁₀ protein amount per barcode (ng/barcode; vertical axis) for the spike-in proteins. Each point represents one spike-in protein. Fitted regression lines, slopes and intercepts are shown. These calibration equations were used to convert barcode intensities into absolute protein amounts for each sublibrary.

**Extended Figure 2.**
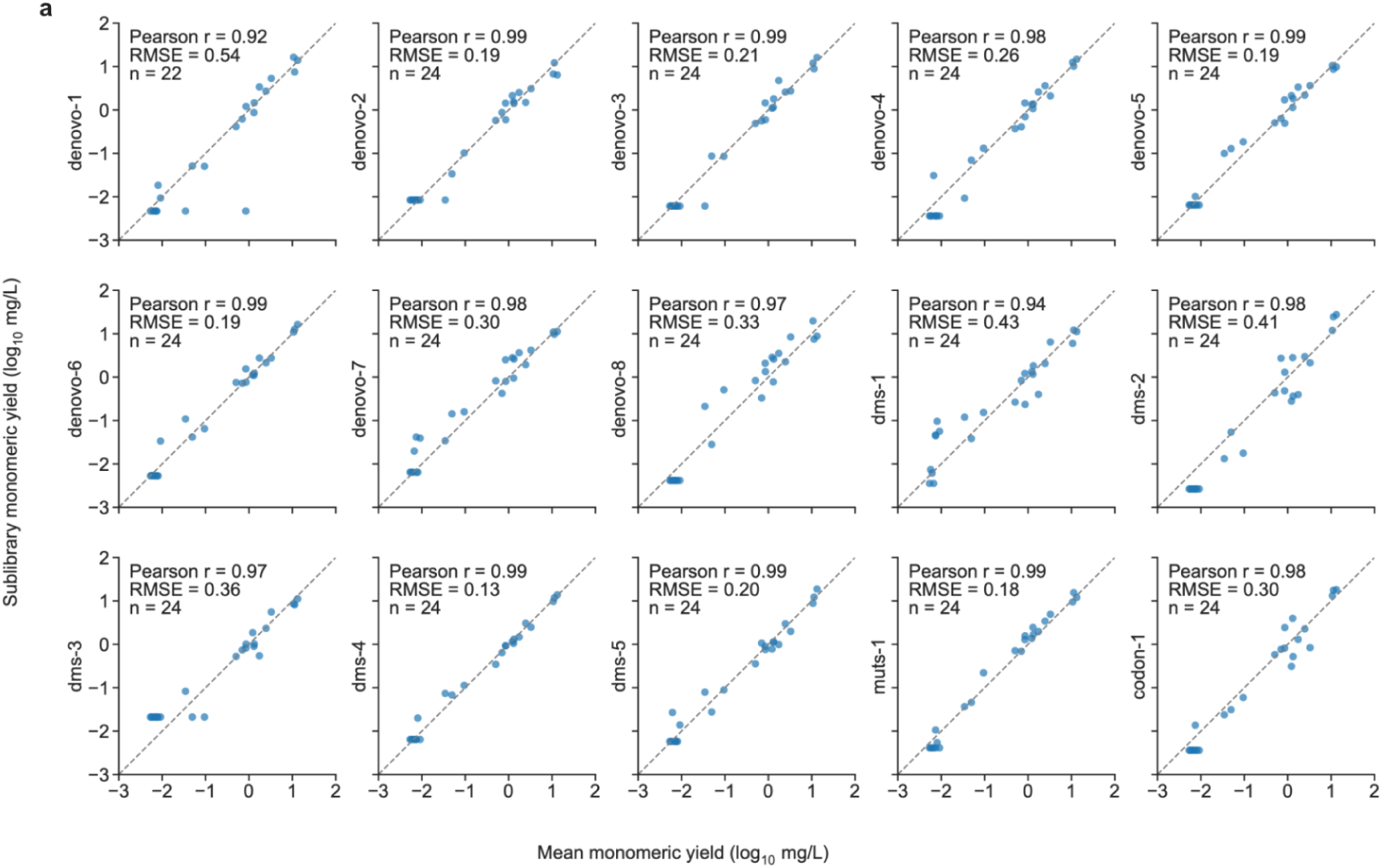
Shared-control agreement across sublibraries after our calibration. **a**, Comparison of calibrated log₁₀ monomeric yield estimates for shared control proteins in each sublibrary (vertical axis) with their cross-sublibrary mean log₁₀ yields (horizontal axis). Yields are expressed in mg/L before log₁₀ transformation. Each point represents one shared control protein, and dashed diagonal lines indicate y = x. Pearson correlation coefficients, root mean square errors (RMSEs) and sample sizes are shown. Each sublibrary contains 24 evaluable controls, except denovo-1, which contains 22 because all barcodes for two control proteins failed the NGS read-count threshold and were excluded.

**Extended Figure 3.**
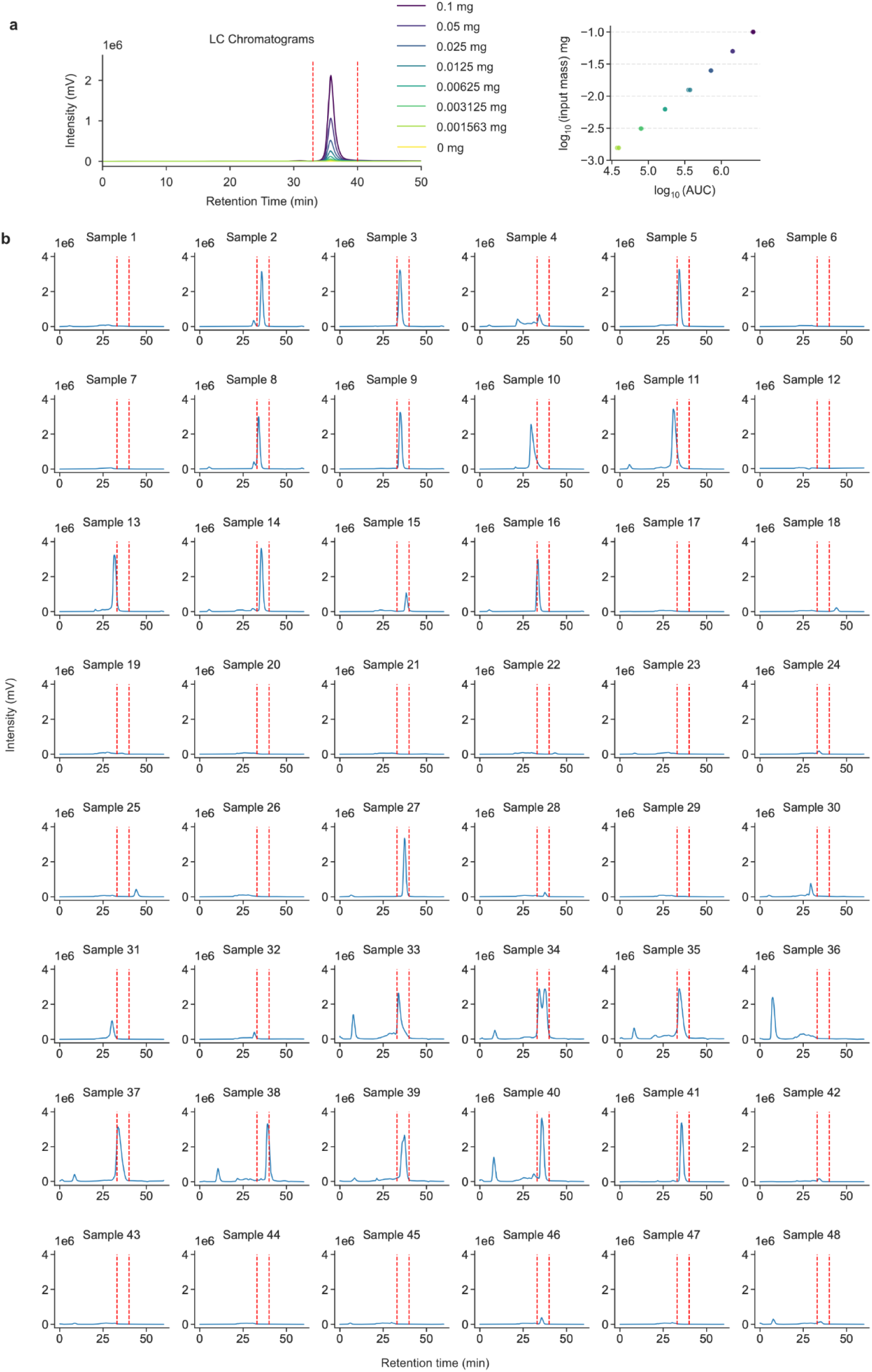
Quantification of individual proteins by HPLC–SEC. **a**, HPLC–SEC traces of a cytochrome c dilution series and the log–log relationship between integrated peak area and injected protein mass used for external calibration. **b,** Chromatograms for the 48 individually expressed and purified proteins compared with pooled-assay estimates in Fig. 1h. The region between the red dashed lines was used to integrate the monomer peak. Each construct was expressed, purified and measured once.

**Extended Figure 4.**
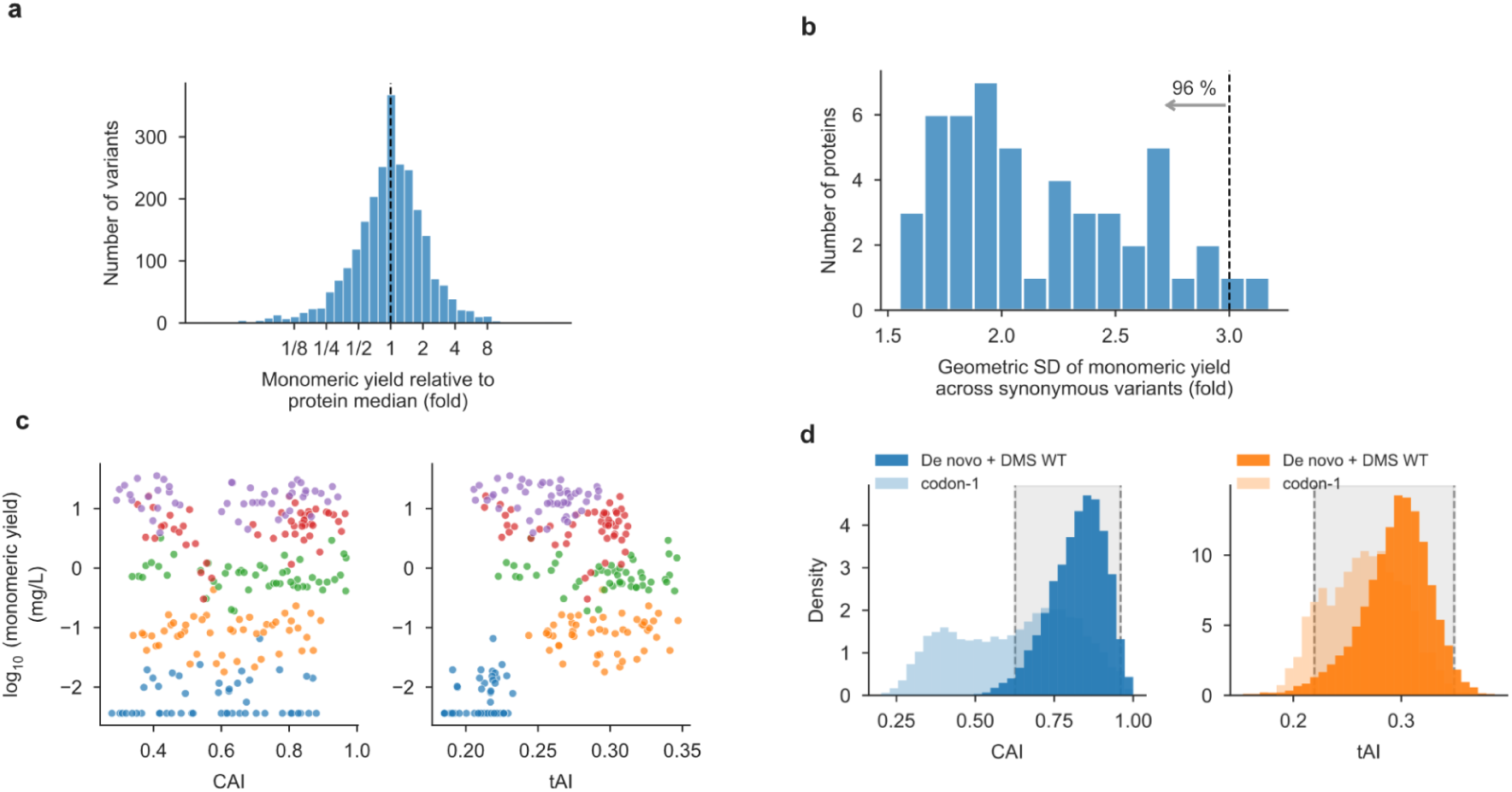
Synonymous codon usage shows little association with monomeric yield. **a,** Distribution of monomeric yields across synonymous coding variants, normalized to the median yield of variants encoding the same protein. The dashed line marks a relative yield of 1 (= median). **b,** Distribution of the geometric standard deviation of monomeric yield across synonymous coding variants for each of 50 parent proteins. For 96% of proteins, the geometric standard deviation was below 3-fold (dashed line). **c,** Log₁₀ monomeric yield (mg/L) versus CAI (left) and tAI (right) for representative five parent proteins. Each point is a synonymous coding variant; colours identify parents. Yield values are shown before clipping at the lower quantification threshold of −1.5. **d,** Distributions of CAI (left) and tAI (right) for coding sequences for de novo designs and DMS wild-type controls (dark colours) and the synonymous-variant library, codon-1 (light colours). Dashed lines and grey shading indicate the central 95% ranges for the de novo designs and DMS wild-type controls: 0.63–0.96 for CAI and 0.22–0.35 for tAI.

**Extended Figure 5.**
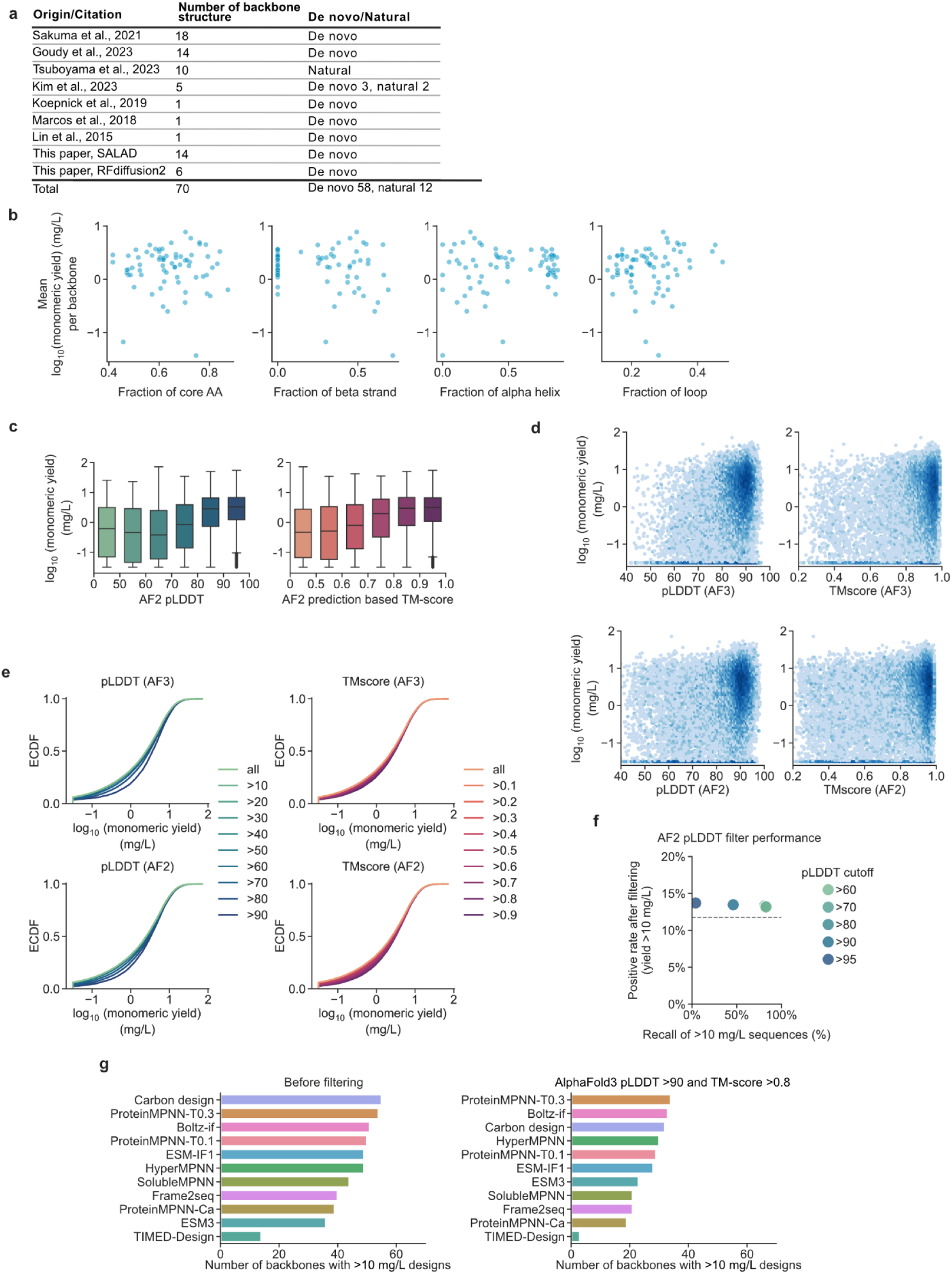
Relationship between monomeric yield and backbone descriptors or structure-confidence filtering. **a,** Sources of the 70 backbone structures used for inverse-folding benchmarking, divided into de novo designs and natural domains. **b,** Backbone-level mean log₁₀ monomeric yield versus the core-residue fraction and β-strand, α-helix and loop fractions. Each point represents one backbone. **c,**Yield distributions binned by AlphaFold2 pLDDT and TM-score. Boxes show medians and interquartile ranges, with 1.5-interquartile-range whiskers. **d,** Yield versus pLDDT or TM-score for AlphaFold3 (top) and AlphaFold2 (bottom). Each point is a design. **e,** Empirical cumulative distributions of yield after filtering at the indicated pLDDT or TM-score thresholds, for AlphaFold3 (top) and AlphaFold2 (bottom). **f,** Positive rate versus recall for AlphaFold2 pLDDT cutoffs at TM-score >0.8. Positives have yield >10 mg/L; the dashed line marks the unfiltered positive rate. **g,** For each inverse-folding method, numbers of distinct backbone structures represented among the sequences retained high-yield (>10 mg/L) sequences before filtering (left), after filtering (right) by AlphaFold pLDDT >90 and TM-score >0.8. A backbone is counted once if at least one sequence in the corresponding set was designed from it.

**Extended Figure 6.**
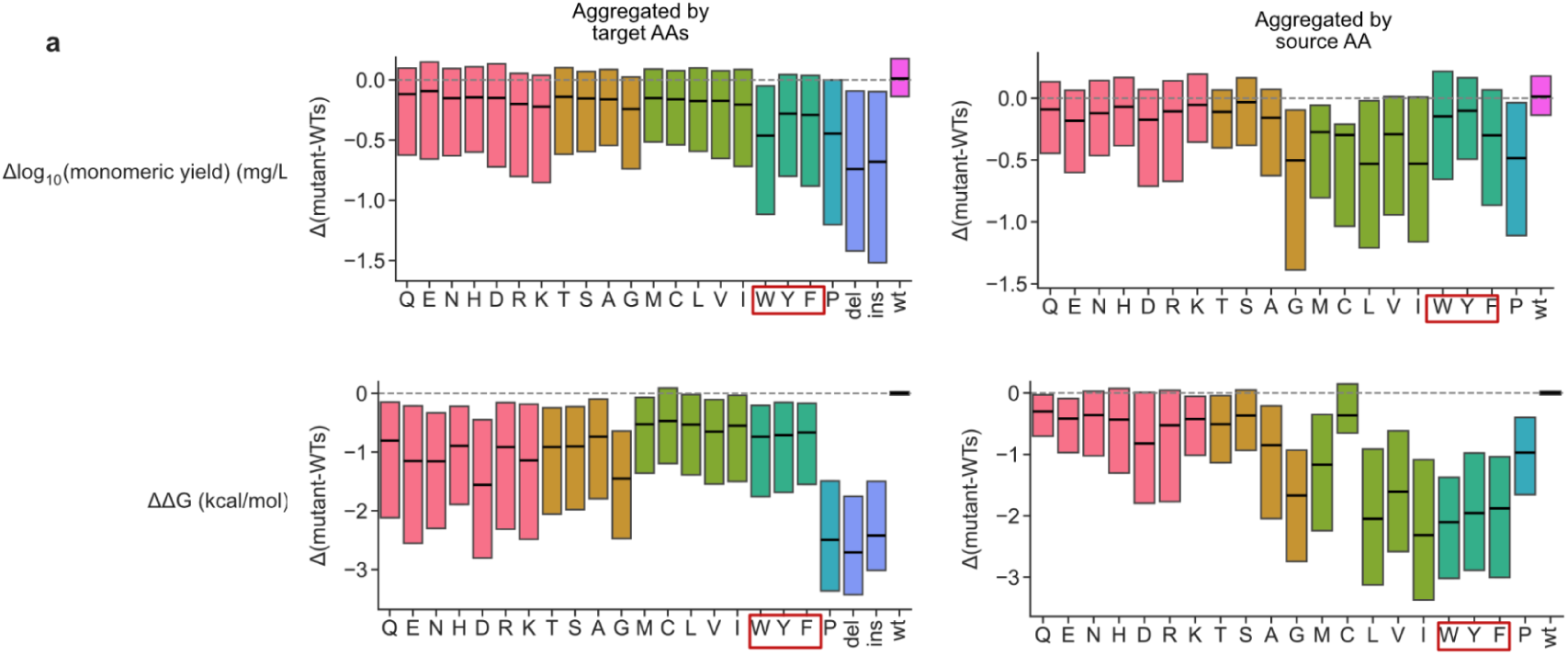
Introducing or replacing aromatic amino acids has distinct effects on folding stability and monomeric yield. **a,** Box plots showing the distributions of mutational effects underlying the summaries in Fig. 3c. Mutational effects are mutant-minus-wild-type changes in log₁₀ monomeric yield (top) and ΔG (bottom), grouped by introduced amino acid (left) or wild-type source amino acid (right). Boxes show medians and interquartile ranges without extended whiskers or outlier markers. Deletion and insertion classes are shown only in the introduced-amino-acid panels. Red outlines highlight the amino acid groups (W, Y and F) that showed pronounced divergence between mutational effects on monomeric yield and ΔG in Fig. 3c.

**Extended Figure 7.**
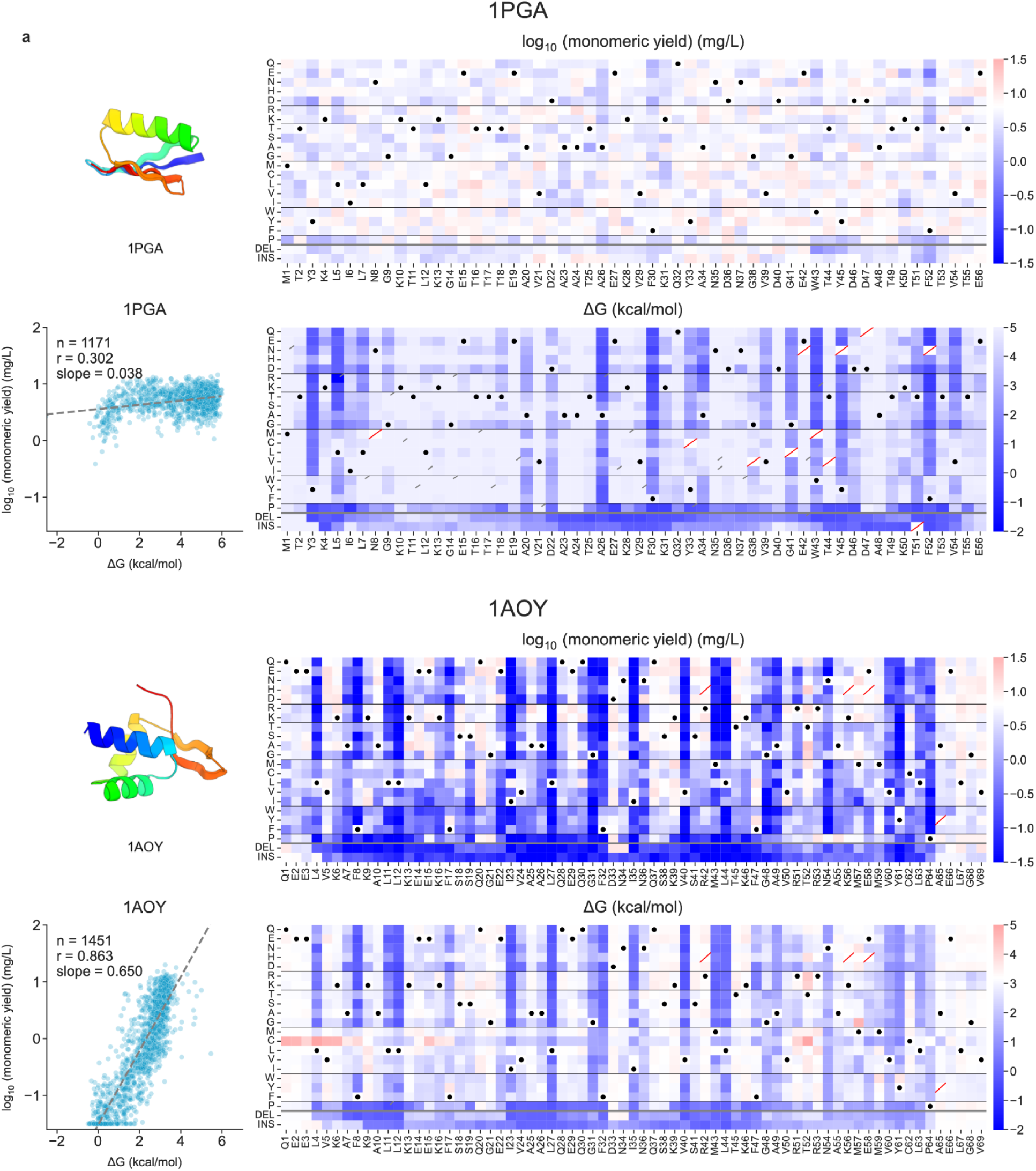
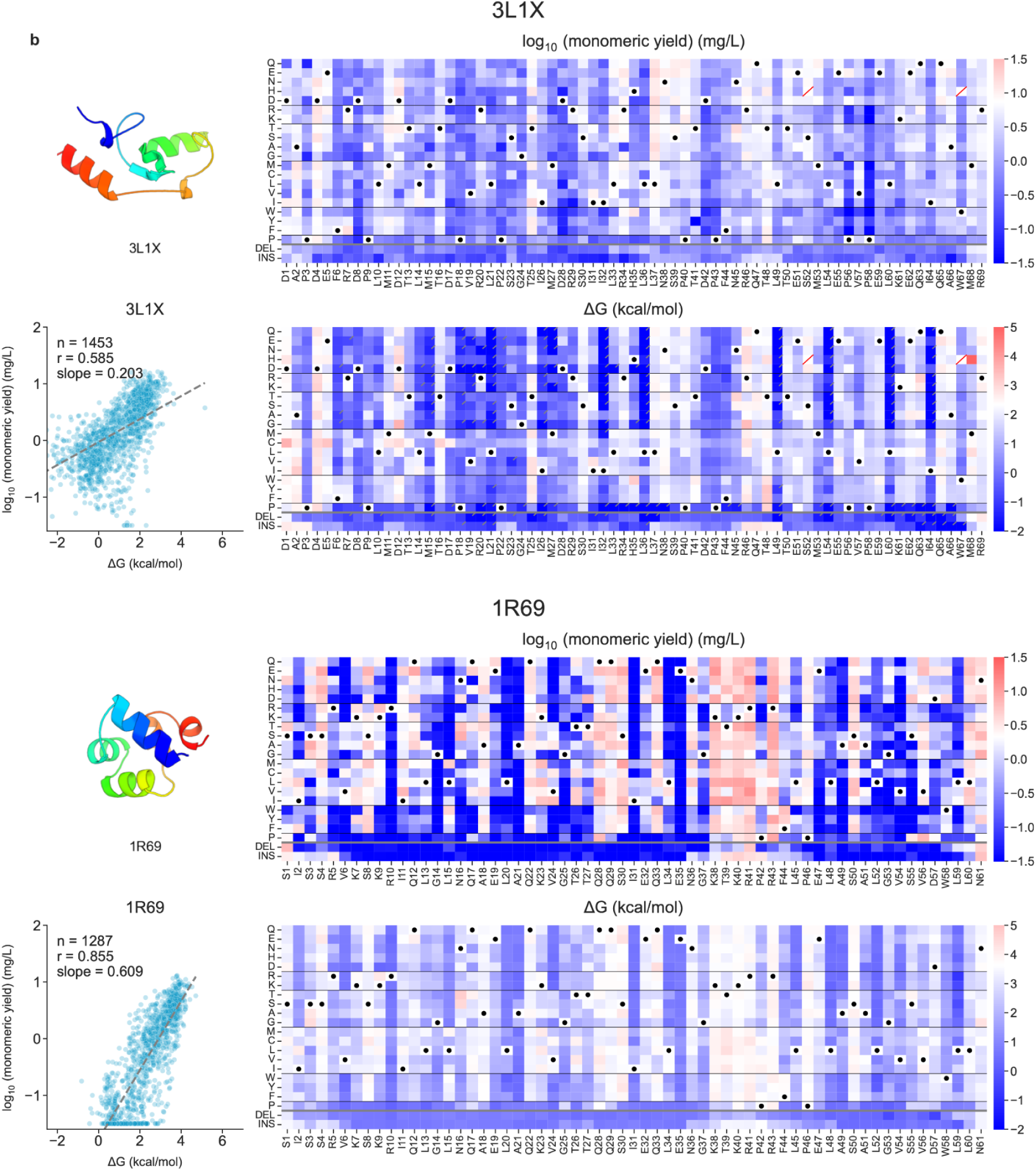
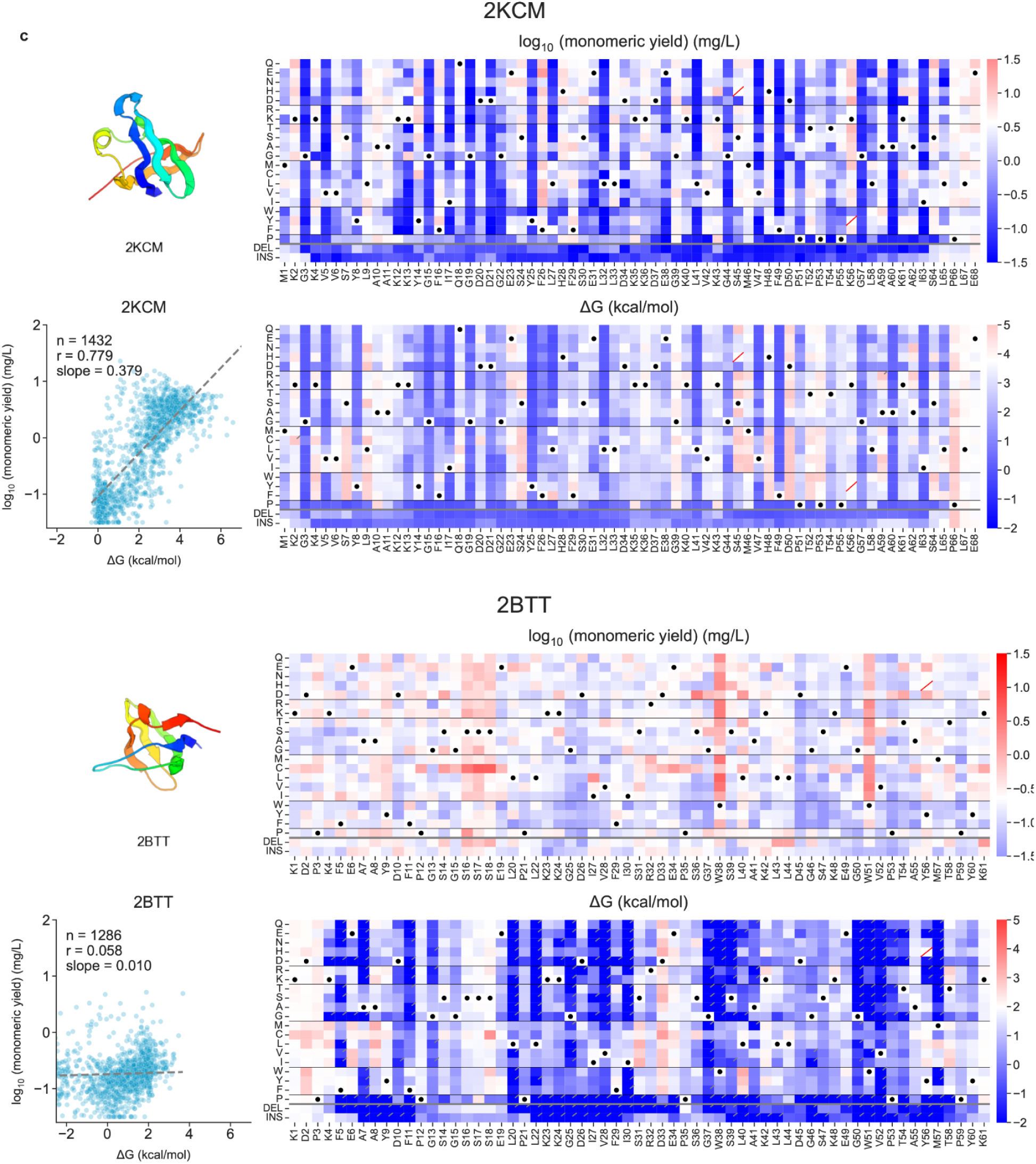
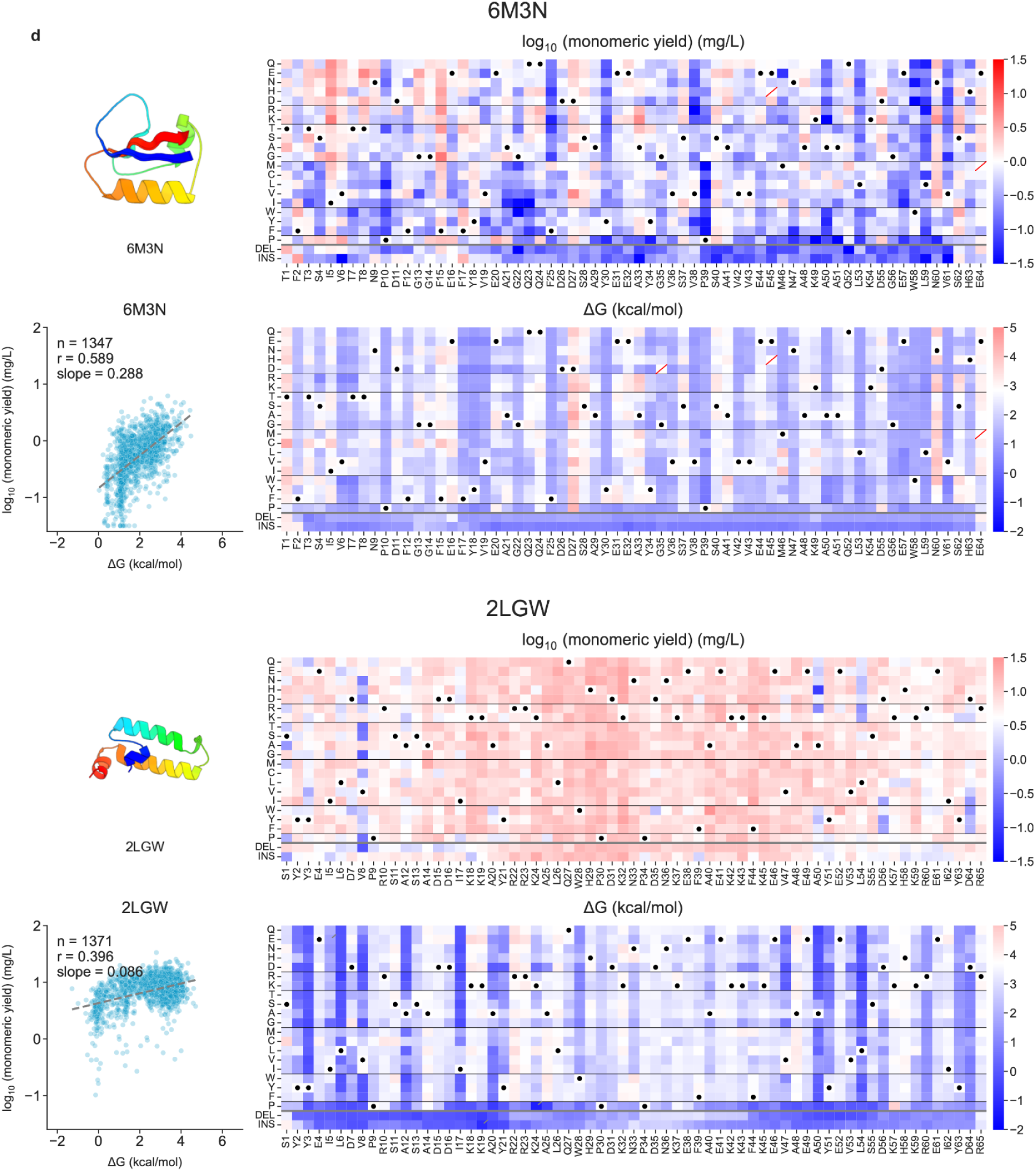
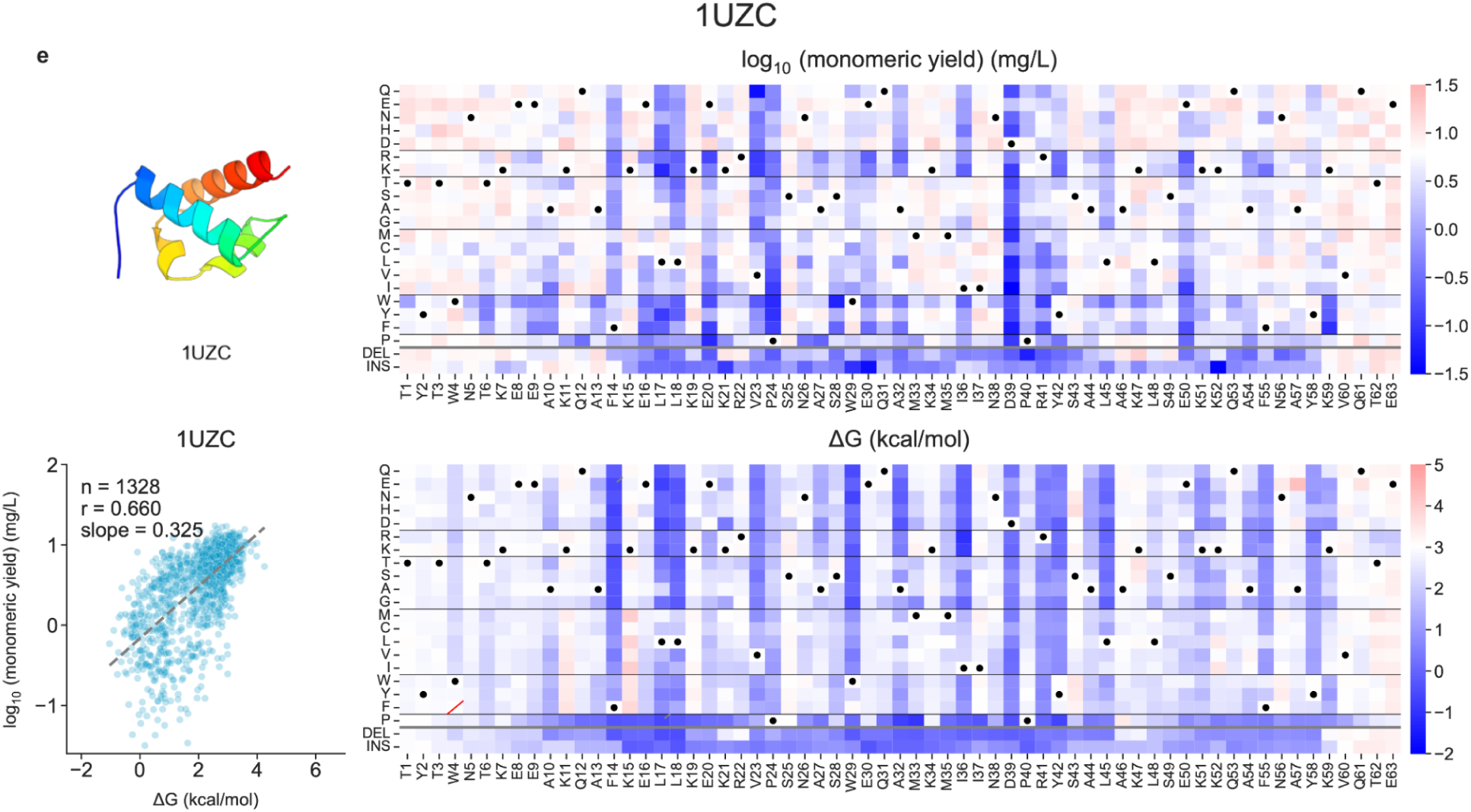
Mutational landscapes of monomeric yield and folding stability across nine protein domains. **a–e,** Structural models, yield–stability scatter plots and position-resolved heatmaps for variants of 1PGA and 1AOY (**a**), 3L1X and 1R69 (**b**), 2KCM and 2BTT (**c**), 6M3N and 2LGW (**d**), and 1UZC (**e**). Scatter plots show log₁₀ monomeric yield versus folding stability (ΔG), with sample sizes, Pearson correlation coefficients and regression slopes indicated. Heatmap columns correspond to sequence positions, and rows represent amino acid substitutions, deletions (DEL) and alanine insertions (INS). Black dots indicate wild-type residues, red diagonal marks indicate unavailable values, and black diagonal marks indicate uncertain ΔG estimates (95% CI >0.5 kcal/mol). Monomeric yield is shown as log₁₀[yield/(mg/L)], and ΔG is expressed in kcal/mol.

**Extended Figure 8.**
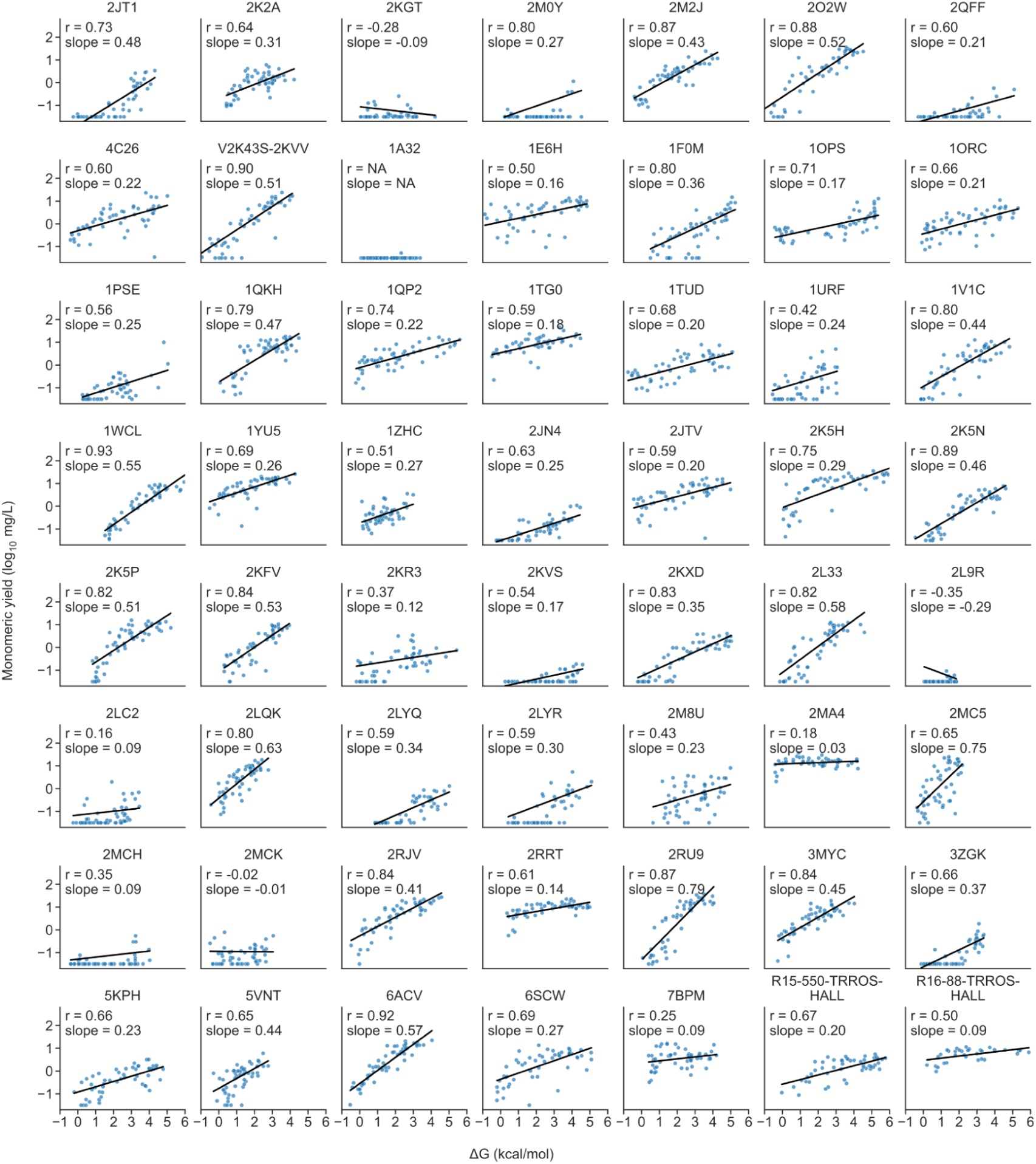
Stability–yield relationships for selected-mutant sets. Each panel shows log₁₀ monomeric yield (mg/L) versus ΔG for a parental protein and its analysed variants from the selected-mutant collection. Parent identifiers, Pearson correlations and fitted slopes are shown. Black lines indicate linear regression fits using only samples with log₁₀ monomeric yield > −1.5, excluding those at the lower quantification limit. The slope for 1A32 is reported as NA because nearly all samples were below the quantifiable range and were assigned the floor value of −1.5.

**Extended Figure 9.**
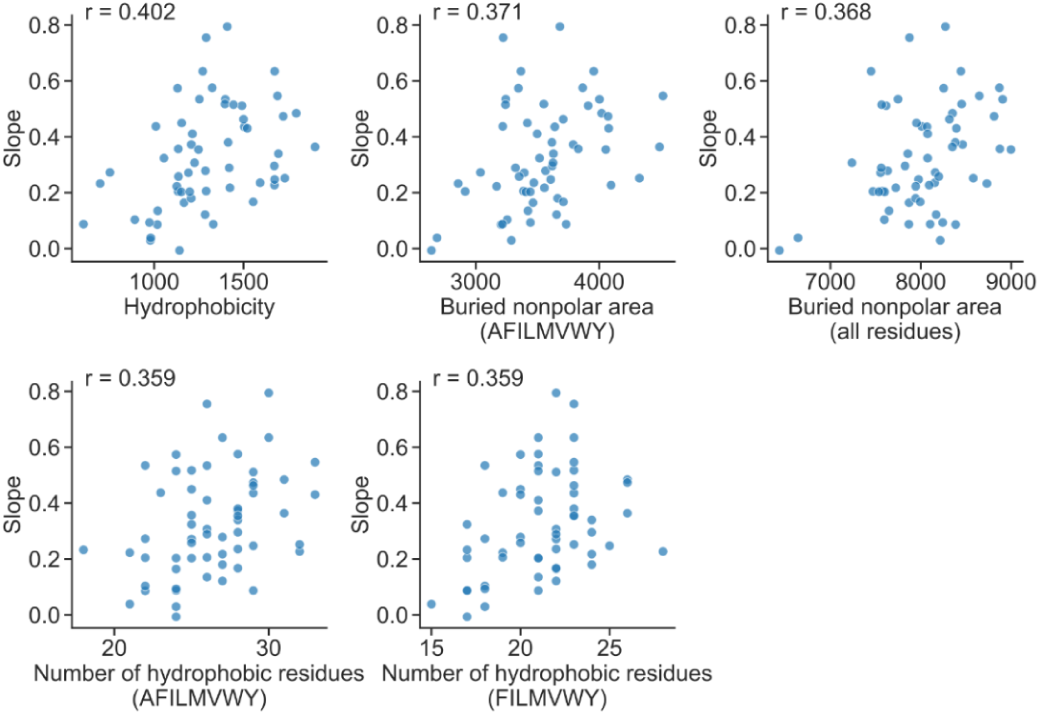
Relationships between hydrophobicity descriptors and stability–yield slopes. Protein-specific slopes from regressions of log₁₀ monomeric yield on ΔG are plotted against the displayed sequence or structural descriptors. Each point represents a parental protein. Pearson correlations are shown in each panel. The descriptors correspond to those summarized in Fig. 3g.

**Extended Figure 10.**
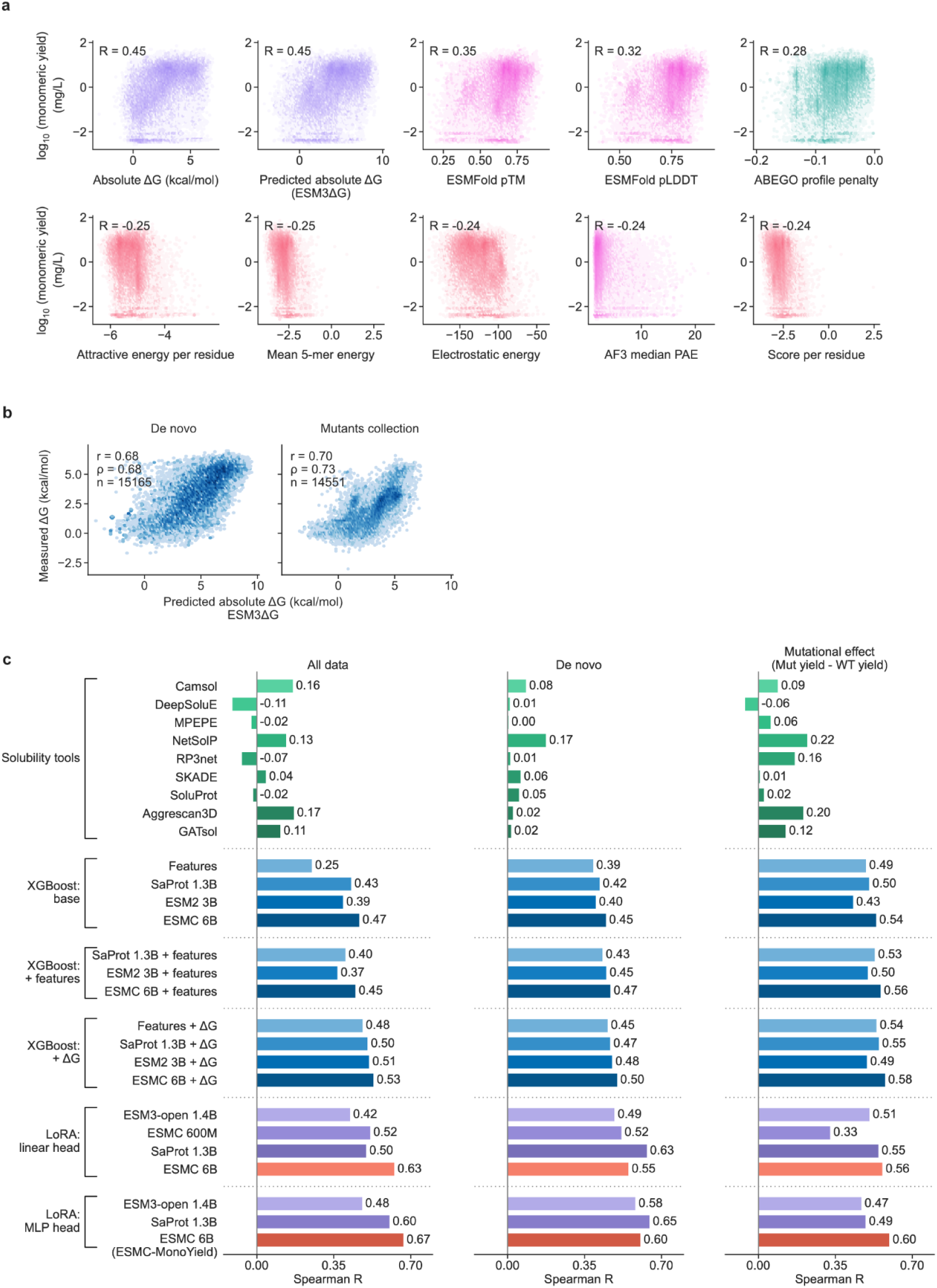
Associations between individual features and monomeric yield. **a,** monomeric yield versus measured ΔG, ESM3ΔG-predicted ΔG, structure-confidence metrics and selected structural or energetic features. Unclipped log₁₀ monomeric yield values were used for plotting and correlation analyses. Pearson correlations are displayed. Colours match the feature categories in Fig. 4a. **b,** measured versus ESM3ΔG-predicted folding stability for the de novo and mutant collections. Pearson and Spearman coefficients and sample sizes are displayed; the units are kcal/mol. **c,** Spearman correlations for trained models and existing predictors on the combined evaluation set (left), de novo designs (middle) and mutational effects (right). Mutational effects are mutant-minus-wild-type differences in log₁₀ monomeric yield. Trained models were assessed using grouped cross-validation as described in Methods.

**Extended Figure 11.**
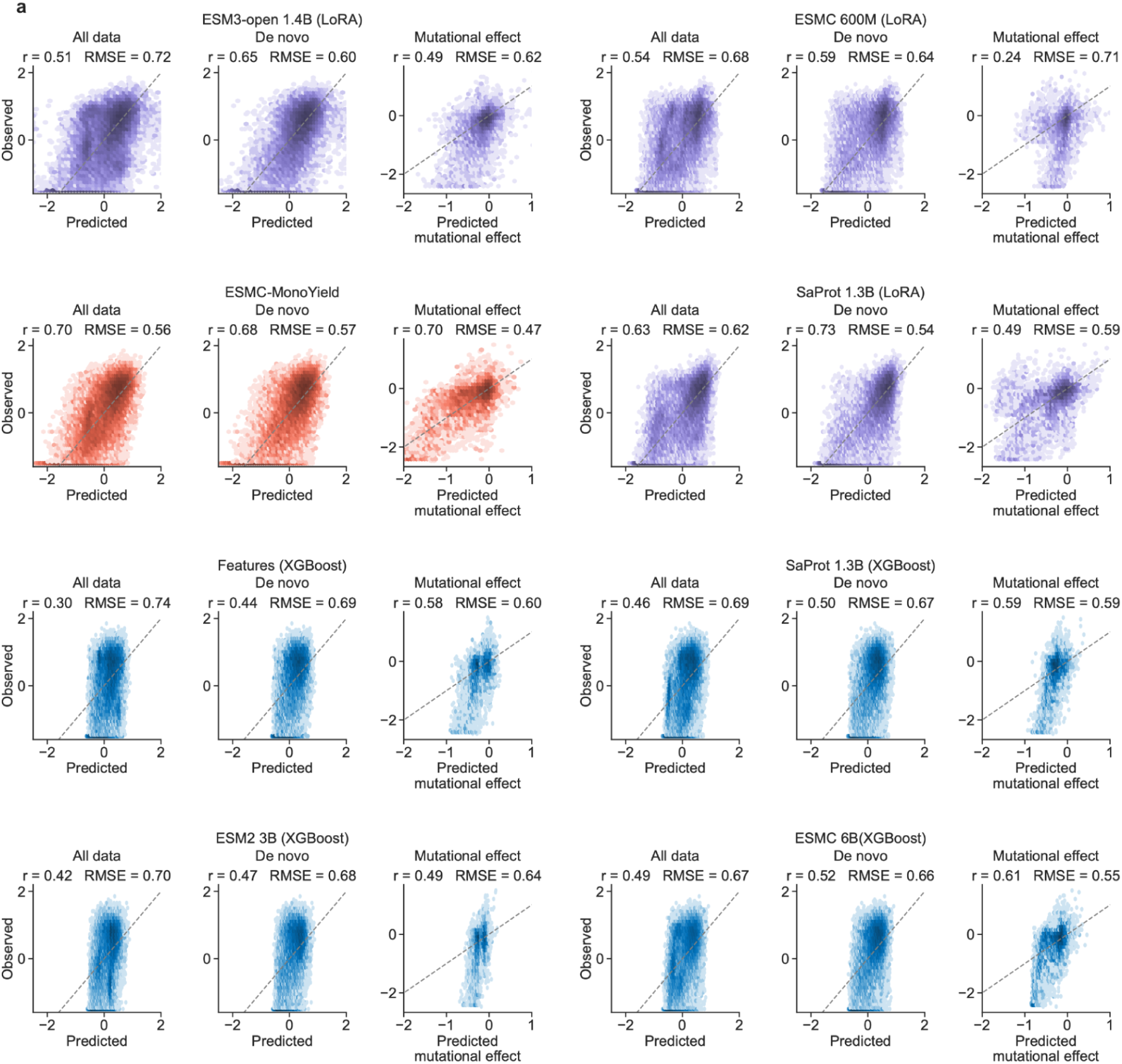
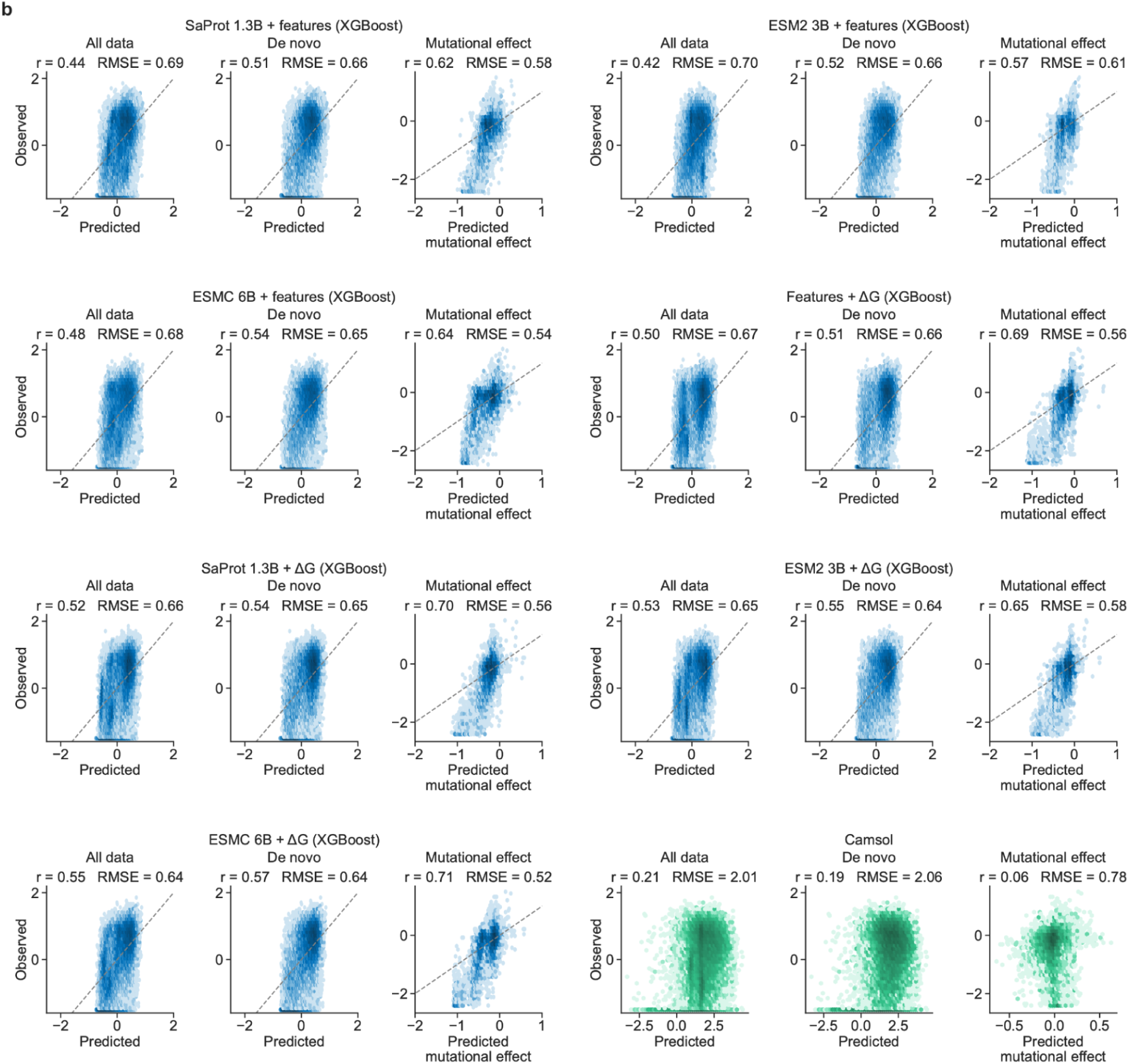
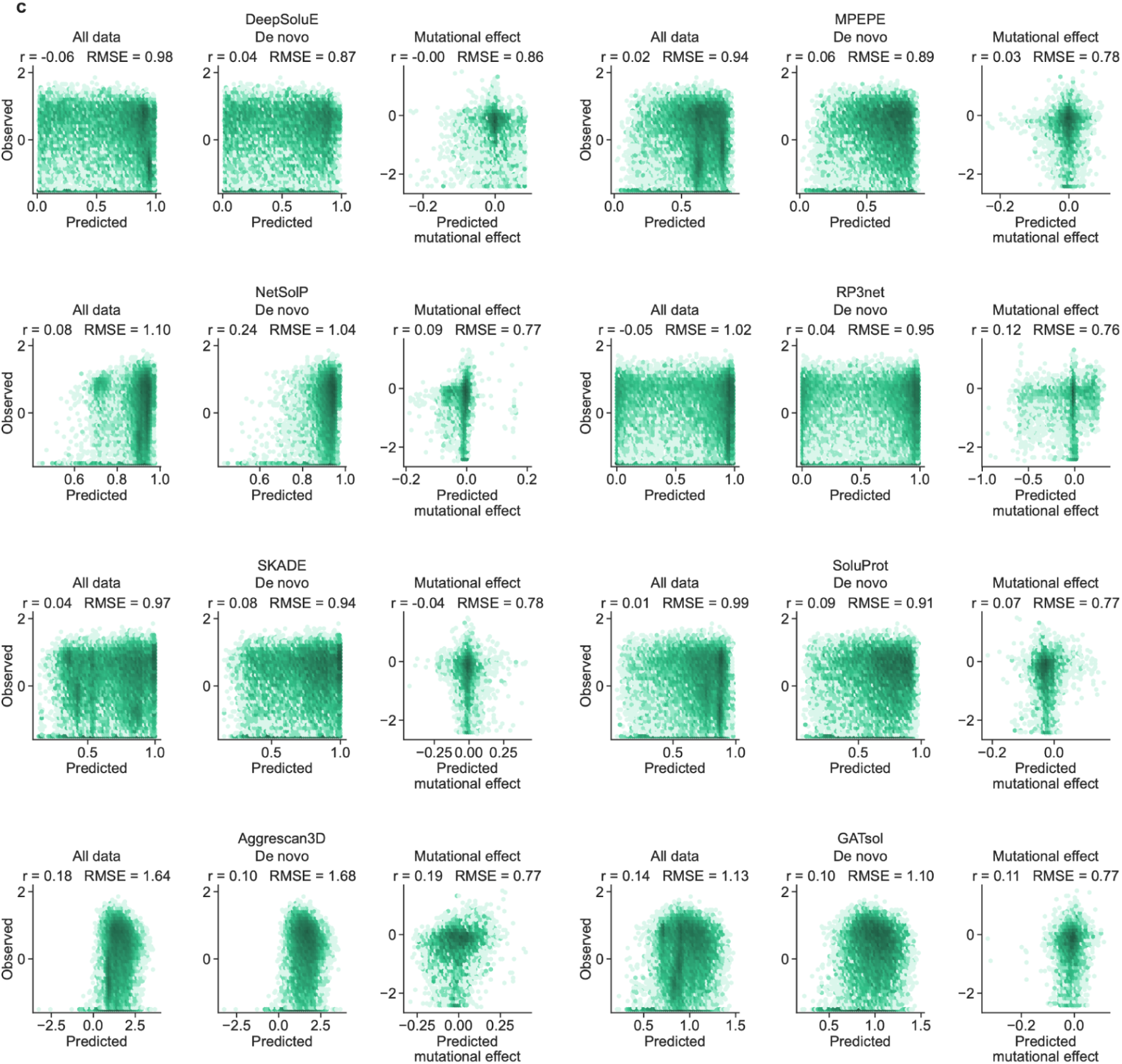
Comparison of predictive models for monomeric yield and mutational effects. **a,** Predictions from pLMs fine-tuned using low-rank adaptation (LoRA) with regression heads, including ESMC-MonoYield, and XGBoost models using either sequence and structural descriptors or pLM embeddings. ESMC 600M used a linear regression head. **b,** Predictions from XGBoost models combining embeddings with sequence and structural descriptors or predicted folding stability (ΔG), and descriptors with predicted ΔG. CamSol scores are shown in the bottom-right group. **c,** Scores from existing solubility and aggregation predictors: DeepSoluE, MPEPE, NetSolP, RP3Net, SKADE, SoluProt, AGGRESCAN3D and GATSol. For each method, the three plots show results for the combined evaluation set (“All data”), de novo designs and mutational effects, from left to right. Sample sizes were n = 25,065, 19,327 and 5,735, respectively, for all methods except DeepSoluE, for which the corresponding sample sizes were n = 19,592, 15,169 and 4,356. DeepSoluE did not return predictions for some sequences owing to tool-specific limitations and was evaluated on the subset with available predictions. Observed values are plotted on the vertical axes and predictions or predictor scores on the horizontal axes. Monomeric yield is expressed as log₁₀[yield/(mg/L)]. Observed and predicted mutational effects are calculated as mutant-minus-wild-type differences in log₁₀ yield or predictor score, as appropriate. Pearson correlation coefficients (r) and root mean square errors (RMSEs) are indicated. Dashed diagonal lines denote y = x for yield-prediction models. Colours distinguish LoRA models (purple), ESMC-MonoYield (red), XGBoost models (blue) and existing predictors (green). Existing predictor scores were oriented so that higher values indicate more favourable predictions. These scores are not calibrated to yield, and their displayed RMSEs are therefore not directly comparable with those of yield-prediction models.

**Extended Figure 12.**
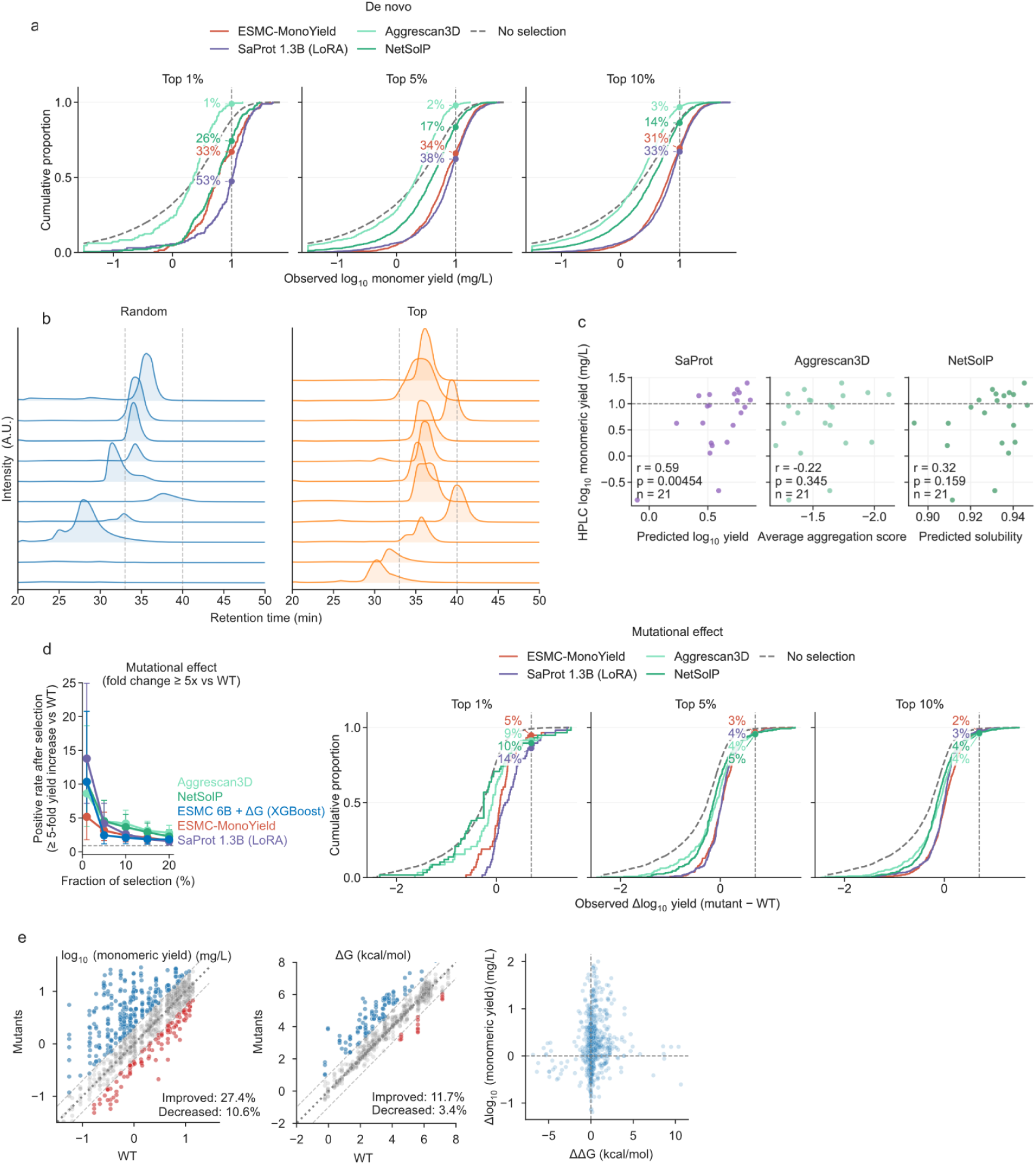
Predictive models support high-yield protein selection, while improving yield through sequence modification remains challenging. **a,** Empirical cumulative distribution functions (ECDFs) of observed log₁₀ monomeric yield for de novo designs among the top 1%, 5% and 10% of candidates ranked by ESMC-MonoYield, SaProt 1.3B LoRA, AGGRESCAN3D or NetSolP. Grey dashed curves show the distribution without selection. Vertical dashed lines mark a monomeric yield of 10 mg/L. Coloured percentages indicate the fractions of selected candidates with monomeric yield >10 mg/L. Yields are expressed in mg/L before log₁₀ transformation. **b,** HPLC–SEC chromatograms of individually expressed BindCraft designs selected randomly (“Random”, blue; n = 10) or prioritized by ESMC-MonoYield (“Top”, orange; n = 11). Traces are vertically offset for display. Dashed vertical lines indicate the interval used to integrate the monomer peak. **c,** Evaluation of SaProt 1.3B LoRA, AGGRESCAN3D and NetSolP on the 21 prospectively tested BindCraft designs. Monomeric yields measured by HPLC–SEC are plotted on a log₁₀ scale against SaProt-predicted log₁₀ yield (left), the AGGRESCAN3D average aggregation score (middle) and NetSolP-predicted solubility (right). The AGGRESCAN3D x-axis is reversed so that predictions become more favorable from left to right across all three panels. Each point represents one design. Pearson correlation coefficients are r = 0.59, −0.22 and 0.32, respectively; P values and sample sizes are shown in each panel. Horizontal dashed lines mark a monomeric yield of 10 mg/L. Yields are expressed in mg/L before log₁₀ transformation. **d,** Selection of mutations predicted to improve monomeric yield. Left, positive rates among the highest-ranked fractions of mutations, comparing SaProt 1.3B LoRA, ESMC-MonoYield, ESMC 6B + ΔG XGBoost, NetSolP and AGGRESCAN3D. Error bars indicate pointwise 95% Wilson confidence intervals for the positive rates. Positives are defined as mutations that increase monomeric yield by at least fivefold relative to wild type; the dashed horizontal line indicates the positive rate without selection. Right, ECDFs of observed mutant-minus-wild-type changes in log₁₀ monomeric yield among the top 1%, 5% and 10% of mutations ranked by ESMC-MonoYield, SaProt 1.3B LoRA, AGGRESCAN3D or NetSolP. Grey dashed curves show the distribution without selection. Vertical dashed lines mark a fivefold increase relative to wild type, and coloured percentages indicate the fractions meeting this criterion. **e,** Effects of PROSS redesign on monomeric yield and folding stability. Measured mutant values are plotted against wild-type values for log₁₀ monomeric yield (left) and ΔG (middle). Blue and red points indicate increases and decreases, respectively, using thresholds of a twofold change in monomeric yield and a 1 kcal/mol change in ΔG. The corresponding fractions are 27.4% increased and 10.6% decreased for yield, and 11.7% increased and 3.4% decreased for folding stability. The right panel compares mutant-minus-wild-type changes in log₁₀ monomeric yield and ΔG (ΔΔG); dashed lines mark zero change. Yield is expressed in mg/L before log₁₀ transformation, and ΔG in kcal/mol.

**Extended Figure 13.**
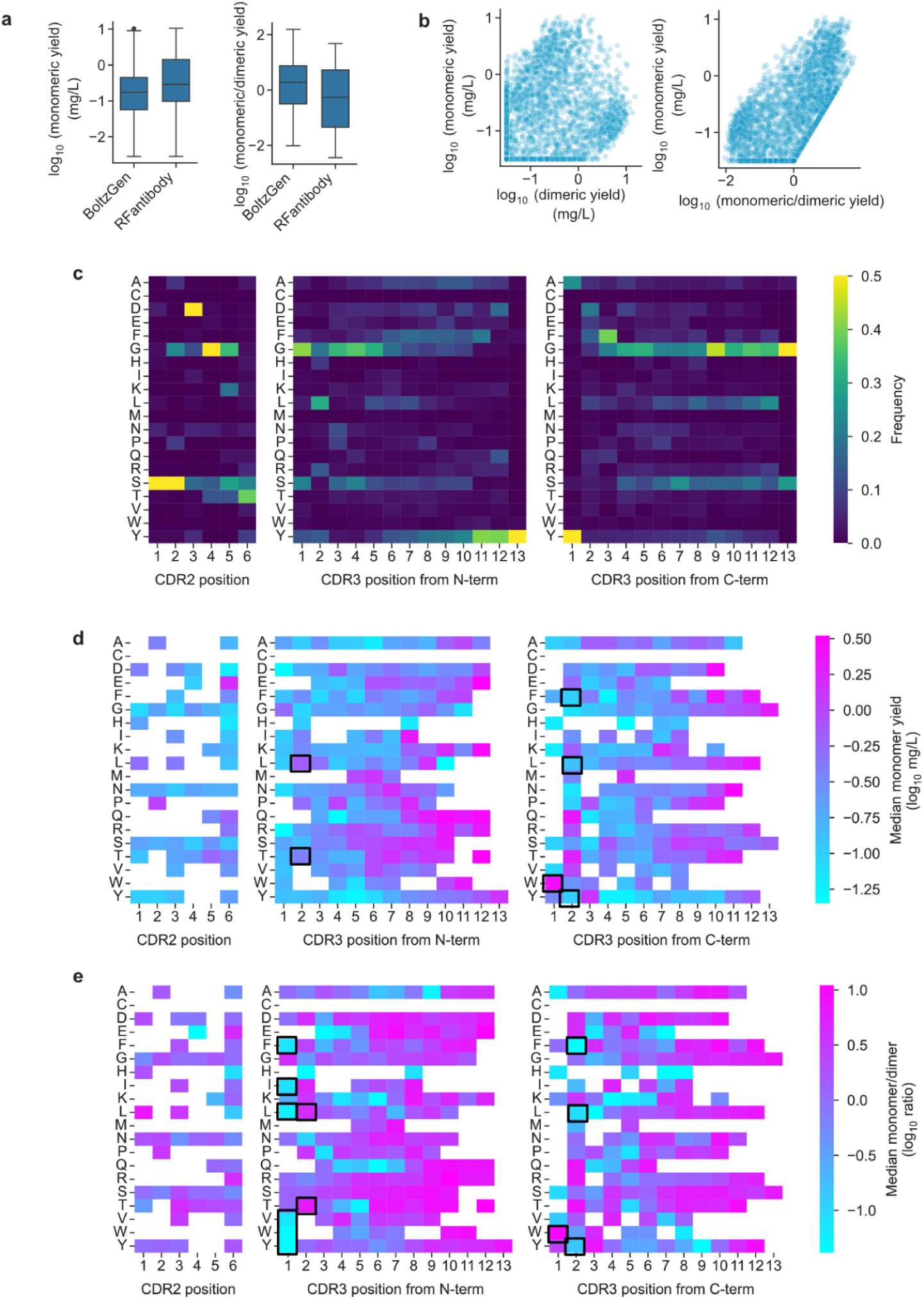
Nanobody yield distributions and associations between CDR sequences and yield. **a,** Monomeric yield and monomer/dimer yield ratio for BoltzGen and RFantibody designs. Both metrics are plotted on log₁₀ scales. Boxes show medians and interquartile ranges, with 1.5-interquartile-range whiskers. Nanobody yields were not clipped in any panel, and monomer/dimer ratios were calculated from the unclipped yields. **b,** Monomeric yield versus dimeric yield (left) and monomer/dimer ratio (right). Each point is a nanobody. The ratio is calculated from the same monomeric and dimeric yields, so the right-hand association is not statistically independent of monomeric yield. **c,** Residue frequencies by CDR2 position and by CDR3 position aligned from the N or C terminus. **d,e,** Median log₁₀ monomeric yield (d) and log₁₀ monomer/dimer ratio (e) for each residue at the corresponding loop position. Cells represented by ten or fewer designs are masked in white. Black outlines highlight selected residue preferences discussed in the text.

**Extended Figure 14.**
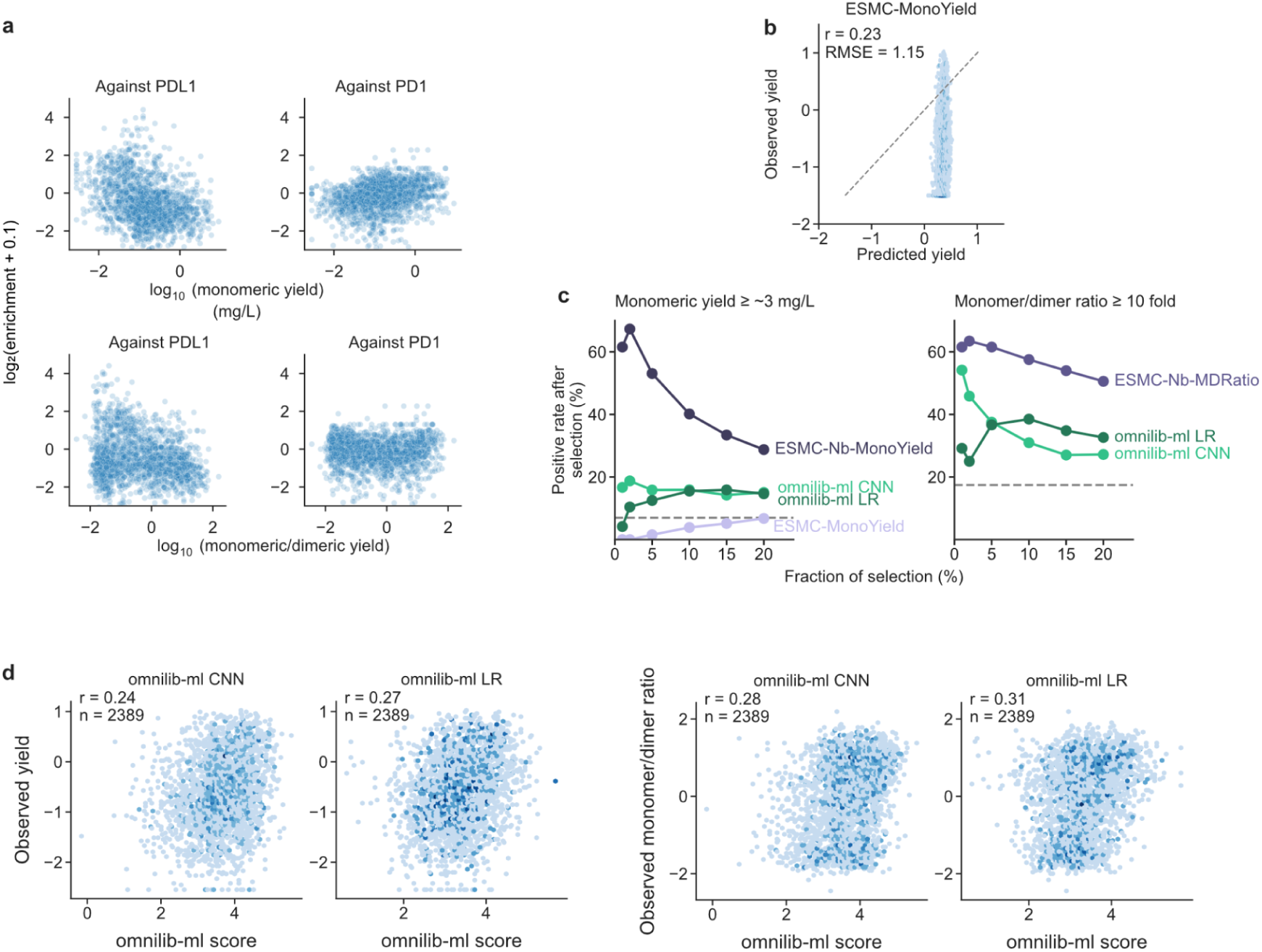
Nanobody target-binding enrichment and yield prediction using previous models. **a,** Target-binding enrichment versus monomeric yield and monomer/dimer ratio for PD-L1- and PD-1-directed designs. Enrichment is displayed as log₂(enrichment + 0.1); yield metrics use log₁₀ scales. Associations are exploratory because target enrichment was modest. In all panels, nanobody yields were used without clipping, and monomer/dimer ratios were calculated from the unclipped yields. **b,** Predictions from the general de novo/DMS-trained ESMC-MonoYield model versus nanobody yield (n = 2,589; Pearson’s r = 0.23; RMSE = 1.15). The predicted range is compressed relative to the observations. **c,** Positive rates after ranking by nanobody-specific ESMC or OmniLib models; the yield panel also includes the general ESMC-MonoYield predictor. Positives have log₁₀[yield/(mg/L)] ≥0.5 (approximately 3.16 mg/L; left) or log₁₀(monomer/dimer ratio) ≥1.0 (a ratio of 10; right). Dashed lines indicate baseline positive rates. **d,** OmniLib CNN and linear-regression scores versus monomeric yield (left two panels) and monomer/dimer ratio (right two panels) for 2,389 designs within the models’ supported CDR length ranges. Pearson correlations are 0.24 and 0.27 for yield and 0.28 and 0.31 for ratio, respectively.

## References

1. Baranowski, C. et al. Can protein expression be “solved”? Trends Biotechnol. 43, 2724–2742 (2025).

2. Williams, D. C., Van Frank, R. M., Muth, W. L. & Burnett, J. P. Cytoplasmic inclusion bodies in Escherichia coli producing biosynthetic human insulin proteins. Science 215, 687–689 (1982).

3. Studier, F. W. Protein production by auto-induction in high density shaking cultures. Protein Expr. Purif. 41, 207–234 (2005).

4. Rosano, G. L. & Ceccarelli, E. A. Recombinant protein expression in Escherichia coli: advances and challenges. Front. Microbiol. 5, 172 (2014).

5. Dauparas, J. et al. Robust deep learning-based protein sequence design using ProteinMPNN. Science 378, 49–56 (2022).

6. Watson, J. L. et al. De novo design of protein structure and function with RFdiffusion. Nature 620, 1089–1100 (2023).

7. Hayes, T. et al. Simulating 500 million years of evolution with a language model. Science 387, 850–858 (2025).

8. Pacesa, M. et al. One-shot design of functional protein binders with BindCraft. Nature 646, 483–492 (2025).

9. Stark, H. et al. BoltzGen: Toward universal binder design. bioRxivorg (2025) doi:10.1101/2025.11.20.689494.

10. Lin, Y., et al. Fast and ultra-capable protein design: Advancing the frontier through atomistic SE(3)-equivariance with Genie 3. bioRxivorg (2026) doi:10.64898/2026.05.01.722168.

11. Cao, L. et al. De novo design of picomolar SARS-CoV-2 miniprotein inhibitors. Science 370, 426–431 (2020).

12. Cao, L. et al. Design of protein-binding proteins from the target structure alone. Nature 605, 551–560 (2022).

13. Kim, D. E. et al. De novo design of small beta barrel proteins. Proc. Natl. Acad. Sci. U. S. A. 120, e2207974120 (2023).

14. Vázquez Torres, S., et al. De novo design of high-affinity binders of bioactive helical peptides. Nature 626, 435–442 (2024).

15. Vázquez Torres, S., et al. De novo designed proteins neutralize lethal snake venom toxins. Nature 639, 225–231 (2025).

16. Klima, J. C. et al. Incorporation of sensing modalities into de novo designed fluorescence-activating proteins. Nat. Commun. 12, 856 (2021).

17. Chen, Y. et al. De novo pan-rhodamine binders for fluorescence microscopy from mammalian cells to extremophiles. Cell 10.1016/j.cell.2026.08.007 (2026) doi:10.1016/j.cell.2026.08.007.

18. Tran, L. et al. De novo design of orthogonal far-red, orange, and green fluorophore-binding proteins for multiplexed imaging. Science 393, 813–819 (2026).

19. Dou, J. et al. De novo design of a fluorescence-activating β-barrel. Nature 561, 485–491 (2018).

20. Yeh, A. H.-W. et al. De novo design of luciferases using deep learning. Nature 614, 774–780 (2023).

21. Lauko, A. et al. Computational design of serine hydrolases. Science 388, eadu2454 (2025).

22. Kim, D. et al. Computational design of metallohydrolases. Nature 649, 246–253 (2026).

23. Garcia, M., Dixit, S. M. & Rocklin, G. J. Evaluating zero-shot prediction of monomeric protein design success by AlphaFold, ESMFold, and ProteinMPNN. Protein Sci. 35, e70453 (2026).

24. Egloff, P. et al. Engineered peptide barcodes for in-depth analyses of binding protein libraries. Nat. Methods 16, 421–428 (2019).

25. Feldman, D., et al. Massively parallel assessment of designed protein solution properties using mass spectrometry and peptide barcoding. bioRxivorg (2025) doi:10.1101/2025.02.24.639402.

26. Akpinaroglu, D. et al. Structure-conditioned masked language models for protein sequence design generalize beyond the native sequence space. bioRxiv 2023.12.15.571823 (2023) doi:10.1101/2023.12.15.571823.

27. Goverde, C. A. et al. Computational design of soluble and functional membrane protein analogues. Nature 631, 449–458 (2024).

28. Ertelt, M. et al. HyperMPNN-A general strategy to design thermostable proteins learned from hyperthermophiles. bioRxivorg 2024.11.26.625397 (2024) doi:10.1101/2024.11.26.625397.

29. Ren, M., Yu, C., Bu, D. & Zhang, H. Accurate and robust protein sequence design with CarbonDesign. *Nat*. Mach. Intell. 6, 536–547 (2024).

30. Castorina, L. V., Ünal, S. M., Subr, K. & Wood, C. W. TIMED-Design: flexible and accessible protein sequence design with convolutional neural networks. Protein Eng. Des. Sel. 37, (2024).

31. Jumper, J. et al. Highly accurate protein structure prediction with AlphaFold. Nature 596, 583–589 (2021).

32. Abramson, J. et al. Accurate structure prediction of biomolecular interactions with AlphaFold 3. Nature 630, 493–500 (2024).

33. Candido, S. et al. Language modeling materializes a world model of protein biology. bioRxiv 2026.06.03.729735 (2026) doi:10.64898/2026.06.03.729735.

34. Itakura, K. et al. Expression in Escherichia coli of a chemically synthesized gene for the hormone somatostatin. Science 198, 1056–1063 (1977).

35. Structural Genomics Consortium et al. Protein production and purification. Nat. Methods 5, 135–146 (2008).

36. Savile, C. K. et al. Biocatalytic asymmetric synthesis of chiral amines from ketones applied to sitagliptin manufacture. Science 329, 305–309 (2010).

37. Rosano, G. L., Morales, E. S. & Ceccarelli, E. A. New tools for recombinant protein production in Escherichia coli: A 5-year update: Recombinant protein production in Escherichia coli. Protein Sci. 28, 1412–1422 (2019).

38. Ojima-Kato, T. Advances in recombinant protein production in microorganisms and functional peptide tags. Biosci. Biotechnol. Biochem. 89, 1–10 (2025).

39. Wrenbeck, E. E. et al. An automated data-driven pipeline for improving heterologous enzyme expression. ACS Synth. Biol. 8, 474–481 (2019).

40. Kuhlman, B. et al. Design of a novel globular protein fold with atomic-level accuracy. Science 302, 1364–1368 (2003).

41. Koga, N. et al. Principles for designing ideal protein structures. Nature 491, 222–227 (2012).

42. Rocklin, G. J. et al. Global analysis of protein folding using massively parallel design, synthesis, and testing. Science 357, 168–175 (2017).

43. Bennett, N. R. et al. Atomically accurate de novo design of antibodies with RFdiffusion. Nature 649, 183–193 (2026).

44. Didi, K., et al. Scaling atomistic protein binder design with generative pretraining and test-time compute. arXiv [cs.LG] (2026) doi:10.48550/arXiv.2603.27950.

45. Huettemann, P. & Kortemme, T. Programming biological behavior with AI-guided de novo protein design. Mol. Cell 86, 4167–4182 (2026).

46. Sumida, K. H. et al. Improving protein expression, stability, and function with ProteinMPNN. J. Am. Chem. Soc. 146, 2054–2061 (2024).

47. Qian, J. et al. Accelerating protein design by scaling experimental characterization. Nat. Commun. 10.1038/s41467-026-76740-9 (2026) doi:10.1038/s41467-026-76740-9.

48. Nikolados, E.-M. & Oyarzún, D. A. Deep learning for optimization of protein expression. Curr. Opin. Biotechnol. 81, 102941 (2023).

49. de Marco, A. Recent advances in recombinant production of soluble proteins in E. coli. Microb. Cell Fact. 24, 21 (2025).

50. Miroux, B. & Walker, J. E. Over-production of proteins in Escherichia coli: mutant hosts that allow synthesis of some membrane proteins and globular proteins at high levels. J. Mol. Biol. 260, 289–298 (1996).

51. Lobstein, J. et al. SHuffle, a novel Escherichia coli protein expression strain capable of correctly folding disulfide bonded proteins in its cytoplasm. Microb. Cell Fact. 11, 56 (2012).

52. Heyde, S. A. H. & Nørholm, M. H. H. Tailoring the evolution of BL21(DE3) uncovers a key role for RNA stability in gene expression toxicity. *Commun*. Biol. 4, 963 (2021).

53. Shilling, P. J. et al. Improved designs for pET expression plasmids increase protein production yield in Escherichia coli. *Commun*. Biol. 3, 214 (2020).

54. Schein, C. H. & Noteborn, M. H. M. Formation of soluble recombinant proteins in Escherichia coli is favored by lower growth temperature. Nat. Biotechnol. 6, 291–294 (1988).

55. Hatahet, F., Nguyen, V. D., Salo, K. E. H. & Ruddock, L. W. Disruption of reducing pathways is not essential for efficient disulfide bond formation in the cytoplasm of E. coli. Microb. Cell Fact. 9, 67 (2010).

56. Calloni, G. et al. DnaK functions as a central hub in the E. coli chaperone network. Cell Rep. 1, 251–264 (2012).

57. Niwa, T., Kanamori, T., Ueda, T. & Taguchi, H. Global analysis of chaperone effects using a reconstituted cell-free translation system. Proc. Natl. Acad. Sci. U. S. A. 109, 8937–8942 (2012).

58. Schramm, F. D., Schroeder, K. & Jonas, K. Protein aggregation in bacteria. FEMS Microbiol. Rev. 44, 54–72 (2020).

59. Wilkinson, D. L. & Harrison, R. G. Predicting the solubility of recombinant proteins in Escherichia coli. Biotechnology (N. Y.) 9, 443–448 (1991).

60. Kiefhaber, T., Rudolph, R., Kohler, H. H. & Buchner, J. Protein aggregation in vitro and in vivo: a quantitative model of the kinetic competition between folding and aggregation. Biotechnology (N. Y.) 9, 825–829 (1991).

61. Kapust, R. B. & Waugh, D. S. Escherichia coli maltose-binding protein is uncommonly effective at promoting the solubility of polypeptides to which it is fused. Protein Sci. 8, 1668–1674 (1999).

62. Marblestone, J. G. et al. Comparison of SUMO fusion technology with traditional gene fusion systems: enhanced expression and solubility with SUMO. Protein Sci. 15, 182–189 (2006).

63. Bivona, L., Zou, Z., Stutzman, N. & Sun, P. D. Influence of the second amino acid on recombinant protein expression. Protein Expr. Purif. 74, 248–256 (2010).

64. Price, W. N., 2nd et al. Large-scale experimental studies show unexpected amino acid effects on protein expression and solubility in vivo in E. coli. Microb. Inform. Exp. 1, 6 (2011).

65. Kramer, R. M., Shende, V. R., Motl, N., Pace, C. N. & Scholtz, J. M. Toward a molecular understanding of protein solubility: increased negative surface charge correlates with increased solubility. Biophys. J. 102, 1907–1915 (2012).

66. Goldenzweig, A. et al. Automated structure- and sequence-based design of proteins for high bacterial expression and stability. Mol. Cell 63, 337–346 (2016).

67. Ojima-Kato, T., Nagai, S. & Nakano, H. N-terminal SKIK peptide tag markedly improves expression of difficult-to-express proteins in Escherichia coli and Saccharomyces cerevisiae. J. Biosci. Bioeng. 123, 540–546 (2017).

68. Verma, M. et al. A short translational ramp determines the efficiency of protein synthesis. Nat. Commun. 10, 5774 (2019).

69. Wang, Y. et al. A 33-residue peptide tag increases solubility and stability of Escherichia coli produced single-chain antibody fragments. Nat. Commun. 13, 4614 (2022).

70. Rosace, A. et al. Automated optimisation of solubility and conformational stability of antibodies and proteins. Nat. Commun. 14, 1937 (2023).

71. Ojima-Kato, T., Nishikawa, Y., Furukawa, Y., Kojima, T. & Nakano, H. Nascent MSKIK peptide cancels ribosomal stalling by arrest peptides in Escherichia coli. J. Biol. Chem. 299, 104676 (2023).

72. Kobo, A., Taguchi, H. & Chadani, Y. Nonspecific N-terminal tetrapeptide insertions disrupt the translation arrest induced by ribosome-arresting peptide sequences. J. Biol. Chem. 300, 107360 (2024).

73. Pejenaute, A. et al. Improving the production and stability of nanobodies. Protein Eng. Des. Sel. 38, gzaf010 (2025).

74. Ojima-Kato, T., Yokoyama, G., Nakano, H., Hamada, M. & Motono, C. Screening and machine learning-based prediction of translation-enhancing peptides that reduce ribosomal stalling in Escherichia coli. *RSC Chem*. Biol. 7, 58–66 (2026).

75. Kudla, G., Murray, A. W., Tollervey, D. & Plotkin, J. B. Coding-sequence determinants of gene expression in Escherichia coli. Science 324, 255–258 (2009).

76. Goodman, D. B., Church, G. M. & Kosuri, S. Causes and effects of N-terminal codon bias in bacterial genes. Science 342, 475–479 (2013).

77. Boël, G. et al. Codon influence on protein expression in E. coli correlates with mRNA levels. Nature 529, 358–363 (2016).

78. Cambray, G., Guimaraes, J. C. & Arkin, A. P. Evaluation of 244,000 synthetic sequences reveals design principles to optimize translation in Escherichia coli. Nat. Biotechnol. 36, 1005–1015 (2018).

79. Nikolados, E.-M., Wongprommoon, A., Aodha, O. M., Cambray, G. & Oyarzún, D. A. Accuracy and data efficiency in deep learning models of protein expression. Nat. Commun. 13, 7755 (2022).

80. Broom, A., Jacobi, Z., Trainor, K. & Meiering, E. M. Computational tools help improve protein stability but with a solubility tradeoff. J. Biol. Chem. 292, 14349–14361 (2017).

81. Havlásek, M. et al. Decoding protein stabilization: Impact on aggregation, solubility, and unfolding mechanisms. J. Chem. Inf. Model. 65, 8688–8701 (2025).

82. Matreyek, K. A. et al. Multiplex assessment of protein variant abundance by massively parallel sequencing. Nat. Genet. 50, 874–882 (2018).

83. Waldo, G. S., Standish, B. M., Berendzen, J. & Terwilliger, T. C. Rapid protein-folding assay using green fluorescent protein. Nat. Biotechnol. 17, 691–695 (1999).

84. Cabantous, S. & Waldo, G. S. In vivo and in vitro protein solubility assays using split GFP. Nat. Methods 3, 845–854 (2006).

85. Wang, T., Badran, A. H., Huang, T. P. & Liu, D. R. Continuous directed evolution of proteins with improved soluble expression. Nat. Chem. Biol. 14, 972–980 (2018).

86. Zutz, A. et al. A dual-reporter system for investigating and optimizing protein translation and folding in E. coli. Nat. Commun. 12, 6093 (2021).

87. Niwa, T. et al. Bimodal protein solubility distribution revealed by an aggregation analysis of the entire ensemble of Escherichia coli proteins. Proc. Natl. Acad. Sci. U. S. A. 106, 4201–4206 (2009).

88. Xiao, R. et al. The high-throughput protein sample production platform of the Northeast Structural Genomics Consortium. J. Struct. Biol. 172, 21–33 (2010).

89. Helen M. Berman, Margaret J. Gabanyi, Andrei Kouranov, David I. Micallef, John Westbrook & Protein Structure Initiative network of investigators. Protein Structure Initiative - TargetTrack 2000-2017 - all data files. Zenodo 10.5281/zenodo.821654 (2017).

90. Ding, Z. et al. MPEPE, a predictive approach to improve protein expression in E. coli based on deep learning. Comput. Struct. Biotechnol. J. 20, 1142–1153 (2022).

91. Habibi, N., Mohd Hashim, S. Z., Norouzi, A. & Samian, M. R. A review of machine learning methods to predict the solubility of overexpressed recombinant proteins in Escherichia coli. BMC Bioinformatics 15, 134 (2014).

92. Sormanni, P., Aprile, F. A. & Vendruscolo, M. The CamSol method of rational design of protein mutants with enhanced solubility. J. Mol. Biol. 427, 478–490 (2015).

93. Hebditch, M., Carballo-Amador, M. A., Charonis, S., Curtis, R. & Warwicker, J. Protein-Sol: a web tool for predicting protein solubility from sequence. Bioinformatics 33, 3098–3100 (2017).

94. Khurana, S. et al. DeepSol: a deep learning framework for sequence-based protein solubility prediction. Bioinformatics 34, 2605–2613 (2018).

95. Hou, Q., Kwasigroch, J. M., Rooman, M. & Pucci, F. SOLart: a structure-based method to predict protein solubility and aggregation. Bioinformatics 36, 1445–1452 (2020).

96. Hon, J. et al. SoluProt: prediction of soluble protein expression in Escherichia coli. Bioinformatics 37, 23–28 (2021).

97. Thumuluri, V. et al. NetSolP: predicting protein solubility in E. coli using language models. Bioinformatics 38, 941–946 (2022).

98. Chen, J., Zheng, S., Zhao, H. & Yang, Y. Structure-aware protein solubility prediction from sequence through graph convolutional network and predicted contact map. J. Cheminform. 13, 7 (2021).

99. Li, B. & Ming, D. GATSol, an enhanced predictor of protein solubility through the synergy of 3D structure graph and large language modeling. BMC Bioinformatics 25, 204 (2024).

100. Li, G., Zhang, N. & Fan, L. ProG-SOL: Predicting protein solubility using protein embeddings and dual-graph convolutional networks. ACS Omega 10, 3910–3916 (2025).

101. Pimtawong, T., Ren, J., Lee, J., Lee, H.-M. & Na, D. A review on computational models for predicting protein solubility. J. Microbiol. 63, e.2408001 (2025).

102. Qian, J., Yang, L., Wang, R. & Qi, Y. Pro4S: Prediction of protein solubility by fusing sequence, structure, and surface. Protein Sci. 35, e70749 (2026).

103. Zambrano, R. et al. AGGRESCAN3D (A3D): server for prediction of aggregation properties of protein structures. Nucleic Acids Res. 43, W306–13 (2015).

104. Cagiada, M., Ovchinnikov, S. & Lindorff-Larsen, K. Predicting absolute protein folding stability using generative models. Protein Sci. 34, e5233 (2025).

105. Alley, E. C., Khimulya, G., Biswas, S., AlQuraishi, M. & Church, G. M. Unified rational protein engineering with sequence-based deep representation learning. Nat. Methods 16, 1315–1322 (2019).

106. Riesselman, A. J., Ingraham, J. B. & Marks, D. S. Deep generative models of genetic variation capture the effects of mutations. Nat. Methods 15, 816–822 (2018).

107. Miyamoto, K. et al. Peptide barcoding for establishment of new types of genotype-phenotype linkages. PLoS One 14, e0215993 (2019).

108. Matsuzaki, Y. et al. Peptide barcoding for one-pot evaluation of sequence-function relationships of nanobodies. Sci. Rep. 11, 21516 (2021).

109. Walter, J. D. et al. Flycodes enable simultaneous preclinical analysis for dozens of antibodies in single cassette-dosed mice. Proc. Natl. Acad. Sci. U. S. A. 122, e2426481122 (2025).

110. Schmirler, R., Heinzinger, M. & Rost, B. Fine-tuning protein language models boosts predictions across diverse tasks. Nat. Commun. 15, 7407 (2024).

111. McMahon, C. et al. Yeast surface display platform for rapid discovery of conformationally selective nanobodies. Nat. Struct. Mol. Biol. 25, 289–296 (2018).

112. Campbell, B. C. et al. mGreenLantern: a bright monomeric fluorescent protein with rapid expression and cell filling properties for neuronal imaging. Proc. Natl. Acad. Sci. U. S. A. 117, 30710–30721 (2020).

113. dos Reis, M., Savva, R. & Wernisch, L. Solving the riddle of codon usage preferences: a test for translational selection. Nucleic Acids Res. 32, 5036–5044 (2004).

114. Sharp, P. M. & Li, W. H. The codon Adaptation Index--a measure of directional synonymous codon usage bias, and its potential applications. Nucleic Acids Res. 15, 1281–1295 (1987).

115. Hsu, C. et al. Learning inverse folding from millions of predicted structures. in Proceedings of the 39th International Conference on Machine Learning (eds. Chaudhuri, K. et al.) vol. 162 8946–8970 (PMLR, 2022).

116. Tsuboyama, K. et al. Mega-scale experimental analysis of protein folding stability in biology and design. Nature 620, 434–444 (2023).

117. Cho, Y., et al. Accurate protein stability prediction for small domains using mega-scale experiments. bioRxivorg (2026) doi:10.64898/2026.05.19.726285.

118. Steinegger, M. & Söding, J. MMseqs2 enables sensitive protein sequence searching for the analysis of massive data sets. Nat. Biotechnol. 35, 1026–1028 (2017).

119. Chen, T. & Guestrin, C. XGBoost: A Scalable Tree Boosting System. in Proceedings of the 22nd ACM SIGKDD International Conference on Knowledge Discovery and Data Mining 785–794 (ACM, New York, NY, USA, 2016).

120. Wang, C. & Zou, Q. Prediction of protein solubility based on sequence physicochemical patterns and distributed representation information with DeepSoluE. BMC Biol. 21, 12 (2023).

121. Tankhilevich, E. et al. RP3Net: a deep learning model for predicting recombinant protein production in Escherichia coli. Bioinformatics 42, btag003 (2026).

122. Raimondi, D., Orlando, G., Fariselli, P. & Moreau, Y. Insight into the protein solubility driving forces with neural attention. PLoS Comput. Biol. 16, e1007722 (2020).

123. Su, J., et al. SaProt: Protein language modeling with structure-aware vocabulary. in The Twelfth International Conference on Learning Representations (2024).

124. Lin, Z. et al. Evolutionary-scale prediction of atomic-level protein structure with a language model. Science 379, 1123–1130 (2023).

125. Mille-Fragoso, L. S. et al. Efficient generation of epitope-targeted antibodies with Germinal. Nat. Biotechnol. 1–10 (2026).

126. Vincke, C. et al. General strategy to humanize a camelid single-domain antibody and identification of a universal humanized nanobody scaffold. J. Biol. Chem. 284, 3273–3284 (2009).

127. Kinoshita, S. et al. Molecular basis for thermal stability and affinity in a VHH: Contribution of the framework region and its influence in the conformation of the CDR3. Protein Sci. 31, e4450 (2022).

128. Wan, Y. et al. Hypervariable loop profiling decodes sequence determinants of antibody stability. Nat. Struct. Mol. Biol. 33, 998–1010 (2026).

129. Hu, E. J., et al. LoRA: Low-Rank Adaptation of large language models. in International Conference on Learning Representations (2022).

130. Corley, N., et al. Biomolecular design through latent reasoning with RFD4-Proteina. Preprint at https://rfd4-proteina.org/assets/paper/RFD4-Proteina.pdf (2026).

131. Ferrari, Á. J. R. et al. Large-scale discovery, analysis and design of protein energy landscapes. Nature 654, 1108–1118 (2026).

132. Martell, C. M. et al. Global analysis of aggregation determinants in small protein domains. bioRxivorg 2025.11.11.687847 (2025) doi:10.1101/2025.11.11.687847.

133. Markin, C. J. et al. Revealing enzyme functional architecture via high-throughput microfluidic enzyme kinetics. Science 373, eabf8761 (2021).

134. Hoover, D. M. & Lubkowski, J. DNAWorks: an automated method for designing oligonucleotides for PCR-based gene synthesis. Nucleic Acids Res. 30, e43 (2002).

135. Demichev, V., Messner, C. B., Vernardis, S. I., Lilley, K. S. & Ralser, M. DIA-NN: neural networks and interference correction enable deep proteome coverage in high throughput. Nat. Methods 17, 41–44 (2020).

136. Park, C. & Marqusee, S. Pulse proteolysis: a simple method for quantitative determination of protein stability and ligand binding. Nat. Methods 2, 207–212 (2005).

137. Tien, M. Z., Meyer, A. G., Sydykova, D. K., Spielman, S. J. & Wilke, C. O. Maximum allowed solvent accessibilites of residues in proteins. PLoS One 8, e80635 (2013).

138. Mirdita, M. et al. ColabFold: making protein folding accessible to all. Nat. Methods 19, 679–682 (2022).

139. Zhang, Y. & Skolnick, J. TM-align: a protein structure alignment algorithm based on the TM-score. Nucleic Acids Res. 33, 2302–2309 (2005).

